# Cell Type Specific DNA Signatures of Transcription Factor Binding

**DOI:** 10.1101/2022.07.15.500259

**Authors:** Aseel Awdeh, Marcel Turcotte, Theodore J. Perkins

## Abstract

Transcription factors (TFs) bind to different parts of the genome in different types of cells. These differences may be due to alterations in the DNA-binding preferences of a TF itself, or mechanisms such as chromatin accessibility, steric hindrance, or competitive binding, that result in a DNA “signature” of differential binding. We propose a method called SigTFB (Signatures of TF Binding), based on deep learning, to detect and quantify cell type specificity in a TF’s DNA-binding signature. We conduct a wide scale investigation of 194 distinct TFs across various cell types. We demonstrate the existence of cell type specificity in approximately 30% of the TFs. We stratify our analysis by different antibodies for the same TF, to rule out the possibility of certain technical artifacts, yet we find that cell type specificity estimates are largely consistent when the same TF is assayed with different antibodies. To further explain the biology behind a TF’s cell type specificity, or lack thereof, we conduct a wide scale motif enrichment analysis of all TFs in question. We show that the presence of alternate motifs correlates with a higher degree of cell type specificity in TFs, such as ATF7, while finding consistent motifs throughout is usually associated with the absence of cell type specificity in a TF, such as CTCF. In particular, we observe that several important TFs show distinct DNA binding signatures in different cancer cell types, which may point to important differences in modes of action. Moreover, we find that motif enrichment sometimes correlates with gene expression in TFs with higher cell type specificity. Our comprehensive investigation provides a basis for further study of the mechanisms behind differences in TF-DNA binding in different cell types.

## Introduction

Non-protein coding regions constitute approximately 98% of the human genome [1, 2]. These regions contain complex instructions to regulate gene expression. Differential binding of transcription factors (TFs) to regulatory sites drives differential gene expression, allowing one genome to give rise to a diversity of cell types and tissues. Conversely, some cell types are defined by one or a few master regulatory TFs that they express. However, the same TF may bind different sites or even different preferred DNA sequences across these multiple cell types [3, 4, 5, 6, 7, 8, 9].

Many studies have shown the cell type specific nature of TF binding locations across multiple cell lines and tissues [3, 4, 5, 6, 7, 8, 9, 10, 11]. For example, Lee et al. studied the binding of the TFs MYC and CCCTC binding factor (CTCF) across 11 different human cell types, and found that both showed some degree of cell type specificity in their binding locations [4]. However, MYC had a much greater degree of cell type specificity than CTCF. Fewer than 25% of CTCF binding sites were cell type specific, while more than 87% were cell type specific for MYC. Additionally, binding sites unique to the cell type were associated with cell type specific functions, such as endothelial cell differentiation and positive regulation of developmental growth in fibroblast cells. Due to its higher degree of cell type specificity, MYC is mainly involved with cell type specific functionalities, while CTCF has a more consistent role across cell types. Cell type specific binding patterns have also been observed for other TFs. Estrogen receptors α (ERs), for instance, bind to distinct DNA patterns in cancerous lines, such as breast cancer and endometrial cancer [5]. The cell type specific sites typically have lower affinity to ERs which bind in conjuction with other TFs, unlike the shared sites which have high affinity for ERs. SOX2 is another example that displays dual modalities of cell type specific binding in human embryonic stem cells (hESC) [12], where the co-binding of SOX2 with PAX6 leads to hECS neural differentiation, and the co-occurrence of SOX2 with OCT4 results in self-renewing hECS. In another study, Wang et al. show the cell type specificity of HOXB9 in K562 cells in comparison to other cell types such as HepG2 and H1-hECS [10]. However, there has yet to be a comprehensive and quantitative analysis of cell type specificity across a broad range of TFs.

There are many factors that cause cell type specificity in TF binding. With the exception of pioneer transcription factors [13], TF binding usually occurs in more accessible regions along the genome. Thus, cell type specific chromatin accessibility is one mechanism directing TFs to different parts of the genome [3, 11, 14, 15]. Another factor is direct protein-protein interactions between TFs which lead to the formation of stable regulatory complexes. All constituents of the complex may bind directly to the genome, or tethering interactions may lead to one element of the complex binding to the genome. Complexing with different DNA-binding partners may draw a TF to different parts of the genome. Alternatively, complexing with other proteins may alter the conformation of the protein and change its DNA-binding preferences [3, 9, 10, 16, 17, 18]. For example, PBX and MEIS paralogs have varying expressions across tissues, and their cooperative binding with HOX alters HOX binding specificity [8, 9]. An additional factor that influences the DNA-binding preferences of a TF protein is alternative splicing [19]. Different versions of the same TF, or TF isoforms, can either bind to different parts of the genome leading to differential gene expression, or bind to the same sites along the genome with different binding affinities resulting in varying amounts of gene expression [19, 20]. Other events, such as post-translational modifications, indirect cooperative binding or conversely steric hindrance, can also influence binding [21, 22].

While differential binding of TFs has been well established, here we seek to investigate in a deeper and more systematic way the possible mechanisms underlying differential binding. In particular, we seek to discriminate two major classes of mechanisms: those that are associated with DNA sequence binding signature and those that are not. Importantly, this is a distinct question from whether the same or different sites are bound in different cell types. For example, a TF might bind completely different sites in two cell types, and yet there may be no differentiating DNA sequence features at all. Mechanisms of differential binding that might not show a difference in DNA binding signature include: changes in TF expression level, so that fewer or more sites of the same type are detectably bound; changes in accessibility of sites of the same type; or even differences in data quality or depth, which alter our ability to detect binding. Conversely, mechanisms that would show a difference in DNA binding signature include: conformation-based changes in inherent TF-DNA preferences resulting from alternative splicing or post transcriptional modifications; complexing with different partners; or cooperation or hindrance with other TFs at regulatory sites.

A deeper understanding of genomics, including TF binding, has been achieved through the use of deep learning approaches. DeepBind is one of the first pivotal methods that used convolutional neural networks (CNNs) for the identification of protein binding sites in DNA and RNA sequences [23]. It used a single convolutional layer, and trained multiple single task models – one for each TF and cell line combination. Other methods, such as DeeperBind [24], DanQ [25] and DeepDRN [26], used a combination of both CNNs and recurrent neural networks (RNNs) for the prediction of TF binding sites. DeepSEA [27] and Basset [28] used CNNs to uncover regulatory features from genomic sequences to predict chromatin accessibility along the genome, and more specifically the impact of single nucleotide variants on regulatory regions, such as DNase hypersensitive sites, TF binding sites and histone marks.

Deep learning has also been used to generalize across cell types, where common non-specific preferences of a TF across cell types are used to predict TF binding activity in cell types with little or no data available [29, 30, 31, 32, 33]. This is possible because a TF may have the same binding preferences in multiple cell types [34, 35]. Indeed, databases such as JASPAR [35], HOCOMOCO [36] and Transfac [34], are established based on the largely successful assumption that TF-DNA binding preferences can be specified independent of context, and can even be derived in-vitro, from SELEX or PBM experiments [37, 38]. Yet, several studies, including [3, 4, 5, 6, 7, 8, 9, 10, 11, 39, 40, 41], show exceptions to the rule of invariant TF-DNA binding preferences, or demonstrate other differential DNA signatures in bound regions. In this paper, we propose a new deep learning approach for analyzing TF-DNA binding sites of a TF across a set of cell types. Although formulated as a binding site prediction problem, the purpose of the learning procedure is to detect and quantify the degree of cell type specificity in DNA-binding signatures for the TF across cell types. That is, given known bound regions for a TF across a potentially large number of different cell types (or tissues or conditions), we seek to quantify the extent to which the DNA sequences of those regions contain discriminative signals regarding cell type. This constitutes a first step towards understanding mechanisms that may underlie differential binding of TFs.

For a more comprehensive and thorough investigation of the cell type specificity of TFs, we develop a method called SigTFB (Signatures of TF Binding). We build upon previous deep learning studies in our own work. The network we use is similar to that used in DeepBind [23], augmenting it to differentiate and quantify cell type general versus cell type specific binding. Unlike most previous work, where the learning problem formulation is based on the discrimination of bound versus unbound regions, which may be genomically real unbound sites or sequence permuted sites, all the sites in our problem are bound. That is, sites are bound in different cell types, and our learning problem determines in which cell there is binding based on the DNA sequences. We focus on differences in “DNA signatures” of binding, rather than differences (or similarities) in the binding sites per se. We use the term DNA signatures to encompass the several factors (previously discussed) that could impact the DNA binding preferences of a TF. Our training procedure is inspired by that of ChromDragoNN [42], but uses a direct encoding of cell type in place of gene expression values. Moreover, similar to MTTFSite [31] and FactorNet [29], we utilize multi-task formulations to learn shared binding preferences of a TF in different cell types. Like Novakovsky et al. [43], which combines multi-task learning with transfer learning to predict TF binding in a specific cell type, we employ a multi-task transfer learning formulation for our learning problem. Our work is similar in intent to a recent study of the differential and cooperative binding nature of the two TFs, MEIS and HOXA2, in three mice tissues [9], but different in the approach we employ and much larger in scale.

We conduct a large scale investigation of 194 TFs assayed by one or more antibodies (AB) across various cell types (for a total of 230 distinct TF-AB pairs), identifying TFs that show a difference in DNA-binding preference across multiple cell lines, and quantifying their degree of cell type specificity. We also investigate the consistency in the degree of specificity across the different antibodies for a specific TF, and explore the correlation between the number of cell types available for a specific TF and the degree of cell type specificity. Finally, we further investigate this method to identify features that drive cell type specificity. More specifically, we analyze motifs that are learned by the model for cell type general versus cell type specific binding.

## Results

### A two-step deep learning model to study the differential binding of a transcription factor

To study the differential binding of a TF, we study one TF and antibody (AB) at a time – each combination requiring ChIP-seq data from a minimum of two different cell types. Data from different ABs for the same TF are not mixed,as we hypothesized that varying binding affinities and the off-target binding effects of the different ABs may contribute to “false” differences between cell types. We collect all the ChIP-seq peak sequences from the different cell types for a specific TF-AB from the ENCODE consortium [2]. We identified 194 unique TFs assayed by ChIP-seq using the same AB in at least two cell types. As some TFs were assayed by multiple ABs in different cell types, there was a total of 230 TF-AB pairs with ChIP-seq data in multiple cell lines/types (CLs). The full list of datasets can be found in SI Table 1. We obtained called peaks for each TF-AB-CL combination, and used human genome GRCh38/hg38 to extract 101 bp windows of DNA sequence around the summit of each peak (See Methods).

For each TF-AB pair, we define the following prediction problem: given the peak sequence and a one-hot encoding of cell type, can we predict whether that sequence was bound (i.e. was there a peak) in that cell type? We are not interested in performance in terms of peak prediction as the input peaks are already experimentally measured. Instead, we are interested in developing a supervised learning system to identify sequence features for a specific TF to distinguish binding in one cell type versus another.

We formulate the prediction of differential binding per TF-AB as a two step process using a deep learning architecture (see Figure 1). We trained a separate model for each of the 230 TF and AB combinations, covering 194 distinct TFs in a total of 35 human cell types. To construct the training set for each TF-AB, we merged ChIP-seq peaks that overlap by at least 30bp across the different corresponding cell types, adopting a similar approach to Basset [28]. The training set for each TF-AB corresponds to the collection of these peaks across the cell types. In stage 1 of training, each input instance comprises a 101 bp peak DNA sequence one-hot encoded into a 404-length binary vector. The output is a C length binary vector indicating in which cell types the peak is bound by the TF, with C being the number of cell types.

**Figure 1:**
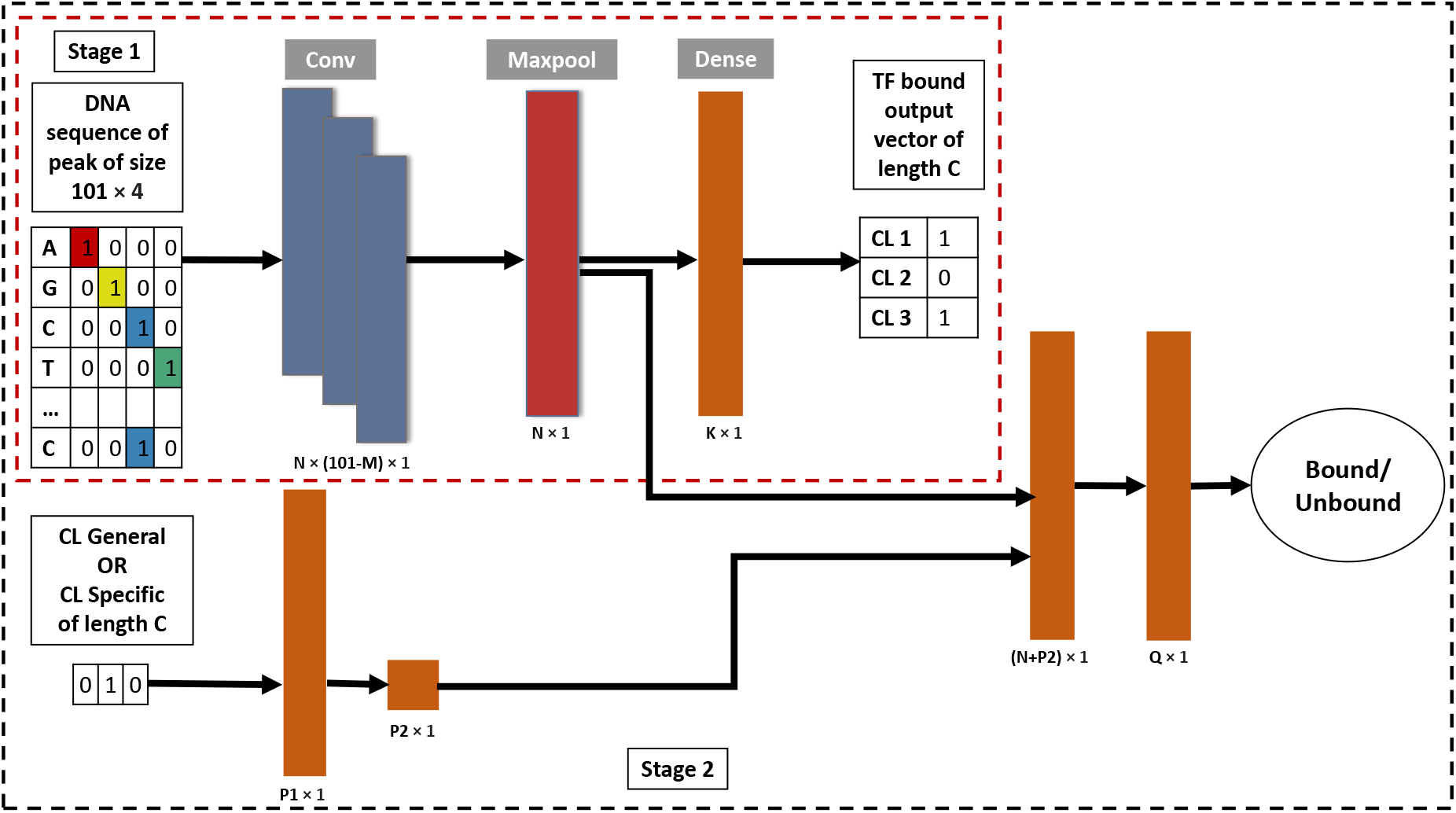
Simplified diagram of Stage 1 and Stage 2 Models. Stage 1 is shown in the red dashed box. Stage 2 is shown in the black dashed box. In Stage 1, the input instance is a one-hot encoded DNA sequence of size 101 × 4. This is passed through the convolutional layer (Conv) with *N* filters of size *M*, through a maxpool layer (Maxpool) of length *N*, a fully connected layer (Dense) of length *K*, then the output layer of length *C* to predict if the TF is bound or not in the different cell types. Stage 2 takes cell type information of length C as input as well. This is first passed through fully connected layers of lengths *P*1 and *P*2, and then concatenated with the output of the convolutional layer from Stage 1. The concatenated output is passed through a fully connected layer of length *Q* to predict whether the sequence is bound or not in that cell type.

The first stage focuses on the sequence part of the input, and is trained as a multi-task classifier for the simultaneous prediction of bound versus unbound regions across various cell types. A rectifier activated convolution layer first transforms the input matrix into an output matrix. The rows in the output matrix correspond to convolutional kernels, where each kernel is a motif detector. This is followed by the max pooling layer which reduces the convolutional matrix into a vector of length equal to the number of convolutional kernels (*N* in Figure 1). The output vector is then passed to a fully connected layer, where it is then compared via the Basset loss function to the true target vector. This constitutes the stage 1 model, and presents the shared features across all cell types for that specific TF-AB.

In stage 2, each unified peak instance is expanded into 2 × *C* instances, where in addition to the 404-length binary vector encoding the peak DNA sequence, there is a length C binary vector encoding cell type. In C of the instances, the cell type vector has one element set to 1 with the rest being zeros, specifying cell type, and a single binary output variable indicating binding or not in that cell type. These are referred to as “cell type specific” (CL Specific) instances. In the remaining instances, the cell type vector is set to all zeros, and the DNA input and binary output remain the same. This is a “cell type general” (CL General) instance, where the approximator is being essentially given a cell type specific instance, but the cell type information is hidden. Intuitively, the difference in prediction performance between the cell type specific and the cell type general cases conveys the degree to which the network is able to identify cell type specific DNA signatures that help in binding prediction.

The weights of the pre-trained multi-task model of stage 1 are used to initialize the parameters of the stage 2 models for the prediction of bound regions for a specific TF in a specified cell type. The cell type vector of length C defined previously is passed through a fully connected layer of length *P*1, and then concatenated with the convolutional output matrix from stage 1. The concatenated vector is passed through a fully connected layer of length *Q*, where the predicted output is then compared to the true target vector via the non-negative log likelihood loss function.

During both training and testing, instances are randomly chosen in mini-batches to have the same number of positive and negative instances from each cell type, and the same number of cell type specific and cell type general instances, avoiding any problems with class imbalance. We use the Ax hyperparameter optimization technique [44] along with 10-fold cross validation to produce an ensemble of 10 models. For each model, we compute a macro-average (across cell types) classification area under the receiver-operator characterstic curve (AUC) separately for cell type specific and cell type general instances. Finally, we take the difference of the two as a measure of learned cell type specificity, using a t-test across the 10 folds to assess statistical significance. A positive difference, where cell type specific predictions are more accurate than cell type general, is taken as evidence for the existence of a cell-type specific DNA signature in the binding sites of that TF.

### ATF7 binding shows cell type specific DNA binding signatures

To demonstrate our approach, we first focus on Activating Transcription Factor 7 (ATF7). As a member of the ATF family, ATF7 binds to the cyclic AMP response element (CRE) with the consensus DNA sequence “TGACGTCA” [45, 46]. Members of the ATF family are basic leucine zipper (bZIP) factors that complex with other bZIP factors to form homodimers or heterodimers [45, 46, 47, 48]. The ATF TFs exhibit varying functionalities in different tissues and cancerous cell lines, including tumour suppressive and oncogenic functions [46]. For instance, the deletion of ATF7 results in the spread of lymphoma [46]. Conversely, the activation of ATF7 in gastric or hepatocellular carcinoma promotes the proliferation of cancer cells. As such, ATF7 may be used as a biomarker for the early detection of tumours in liver and gastric cell lines. Due to the differences observed, we suspect ATF7 to bind to different places along the genome in different cell lines.

To investigate the activity of ATF7, we obtained ChIP-seq peaks from ENCODE [2] in the following four cell lines: GM12878, K562, HepG2 and MCF-7. The cancerous cell lines HepG2, MCF-7 and K562 correspond to liver hepatocellular carcinoma, breast cancer and myelogenous leukemia respectively. GM12878 is a non cancerous lymphoblastoid cell line. Figure 2a shows a Venn diagram of the peak overlaps between the four cell lines. The number of peaks per cell line are shown after the cell type name in brackets. We notice that a mere 1.36% of the total number of peaks across all four cell lines overlap, and the majority of the peaks are unique to one of the four cell lines. For example, 22.36% of the K562 peaks do not overlap with peaks from other cell lines. Moreover, we notice there is greater peak overlap between the pairs HepG2 and MCF-7, and GM12878 and K562.

**Figure 2:**
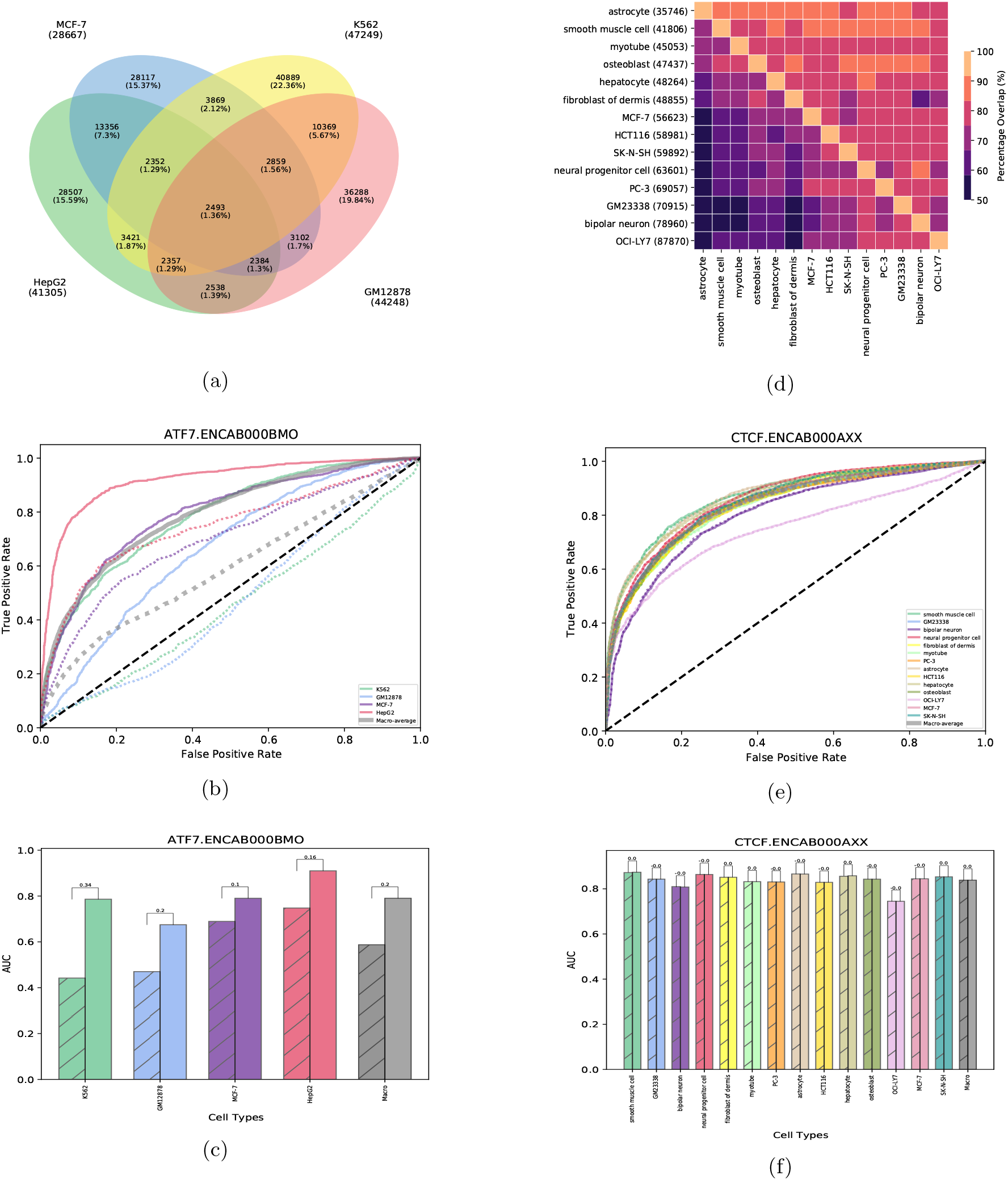
(a) Venn diagram of percentage overlap between cell lines for ATF7. (b) ROC curves per cell line cell line per condition: CL General (dashed line) and CL Specific (solid line) for ATF7. (c) AUC per cell line per condition: CL General (shaded) and CL Specific (not shaded) for ATF7. (d) Heatmap of percentage overlap between 14 cell lines in CTCF.ENCAB000AXX. (e) ROC curves per cell line cell line per condition: CL General (dashed line) and CL Specific (solid line) for CTCF. (f) AUC per cell line per condition: CL General (shaded) and CL Specific (not shaded) for CTCF.

The lack of overlap between peaks in the four cell lines does not imply cell type specificity in DNA binding preference, as sequences in those peaks may be very similar. Differences in output may be due to dissimilarities in terms of noise, bias or even the number of peaks of the ChIP-seq experiments. For instance, HepG2 has over 40,000 peaks while MCF-7 has fewer than 30,000. Therefore, no more than 75% of HepG2 peaks could possibly overlap with MCF-7 peaks

To determine if there are cell type specific DNA signatures in the ATF7 peaks, we applied our deep learning method, SigTFB, as described in the previous section. Figure 2b shows the receiver operating characteristic (ROC) curves for each cell line with and without the cell line identity being provided, as well as averaged performance across all cell lines. The plot shows high variability in site prediction across across cell lines. Predictions for HepG2 (solid red curve) are significantly better than for MCF-7 and K562 (solid purple and green), which are better than for GM12878 (solid blue). Consequently, predictions are more accurate when the network is informed of cell type than when it is not (e.g. solid red versus dashed red curves). This trend is also true for the macro-averaged ROC curve (gray color in Figure 2b). Figure 2c shows the area under the ROC curve (AUC) per cell line per condition for the ATF7 TF, where the shaded and unshaded bars are CL General and CL Specific cases respectively. For each cell line, as well as the macro-averaged result, there is a clear difference between the two conditions. CL Specific classification outperforms CL General classification with a macro-averaged AUC difference of 0.2 (*p* < 0.05; one-sample t-test on AUC difference). Thus, we can conclude that the network has detected DNA signatures discriminating peaks in different cell types. We return to the question of exactly what those signatures are below. First, we examine another transcription factor, CTCF, in detail.

### CTCF binding does not show cell type specific DNA binding signatures

We next examine another transcription factor (TF), CCCTC-binding factor (CTCF). The CTCF binding domain is defined by ll zinc fingers, and is believed to be invariant across cell types – meaning that although the actual binding locations across cell types may vary, the CTCF DNA binding preferences remain the same. CTCF can function as a transcriptional repressor, transcriptional activator, or as an insulator barrier between genomic domains [49, 50, 51]. It also plays a key role in regulating the three-dimensional structure of chromatin. The function of CTCF is greatly dependent on its DNA binding partners.

To test these prior findings, we obtained ChIP-seq data for CTCF in 14 different cell types. These cell types include smooth muscle cell, GM23338, bipolar neuron, neural progenitor cell, fibroblast of dermis, myotube, PC-3, astrocyte, HCT116, hepatocyte, osteoblast, OCI-LY7, MCF-7 and SK-N-SH. The percentage overlap of ChIP-seq peaks between each pair of cell types is shown in Figure 2d, where each entry of the heatmap shows the percentage of peaks of the row’s cell line overlapping peaks in the column’s cell line. Additionally, the number of peaks per cell type are shown in brackets after the row cell type label. Overlap percentages range from approximately 50% to 90%, with an average of 77%. Cell types with fewer peaks tend to be better covered by cell types with more peaks, suggesting an element of peak detection power is at play. For instance, the astrocyte dataset has the fewest peaks at ≈37,000, which are more than 90% covered by the CTCF peaks in every other cell type – even distantly related cell types such as osteoblasts or fibroblasts (first row in Figure 2d).

Figure 2d gives some intuition about the datasets. However, as seen for ATF7, a simple intersection analysis is not sufficient to determine cell type specificity. We further investigated the binding activity of CTCF by training SigTFB on CTCF and its 14 corresponding cell lines. Figure 2e shows the ROC curves for each of the cell types and the macro-averaged ROC across all cell types. Compared to ATF7, there is relatively little difference in binding site predictability across cell types and nearly no difference in predictability for a given cell type, with or without cell type identity information.

Cell types OCI-LY7 (lavender line) and bipolar neuron (indigo line) have the worst prediction performance, and also have the highest number of peaks. Possibly, a greater fraction of these peaks are not genuine, which would explain both inflated peak numbers and prediction difficulty. Figure 2f shows there is little too no difference in the area under the ROC curves (AUC) between CL Specific (solid line) and CL General (broken line) conditions for each cell line (p > 0.5; one-sample t-test on percentage differences). Consequently, these results illustrate the ubiquitous non cell type specific nature of CTCF DNA binding preferences. Importantly, they also demonstrate the specificity of SigTFB, in that it does not incorrectly report cell type specificity where there is none to be found.

### Determining the degree of the cell type specificity of the different transcription factors

Motivated by our results for ATF7 and CTCF, we expanded our study by investigating cell type specificity. Figure 3a displays a scatter plot of the mean AUC of prediction when the network is (y-axis) or is not (x-axis) told what cell type it is predicting for, with the color gradient depending on the negative log 10 p-values. Each point corresponds to a TF-AB combination. We observe a continuum of cell type specificity, where TFs with the least cell type specificity lie in the *x* = *y* diagonal of the scatter plot. For these TFs, the cell type information does not improve prediction. The position of a point along the diagonal may depend on the extent to which there are shared common motifs for the TF across cell types, or even the extent to which the peaks themselves overlap across cell types. Conversely, points lying above the diagonal indicate that the network better predicts binding when informed of the cell type; these are TFs with the most significant cell type specificity. In other words, there are signals in the DNA sequence to which the network responds differently, depending on the cell type for which binding is being predicting. Points in the the upper left corner correspond to TFs where cross-cell type prediction is virtually impossible, but is highly accurate for specific cell types. For such TFs, each cell type is expected to have specific DNA motifs that discriminate its binding sites.

**Figure 3:**
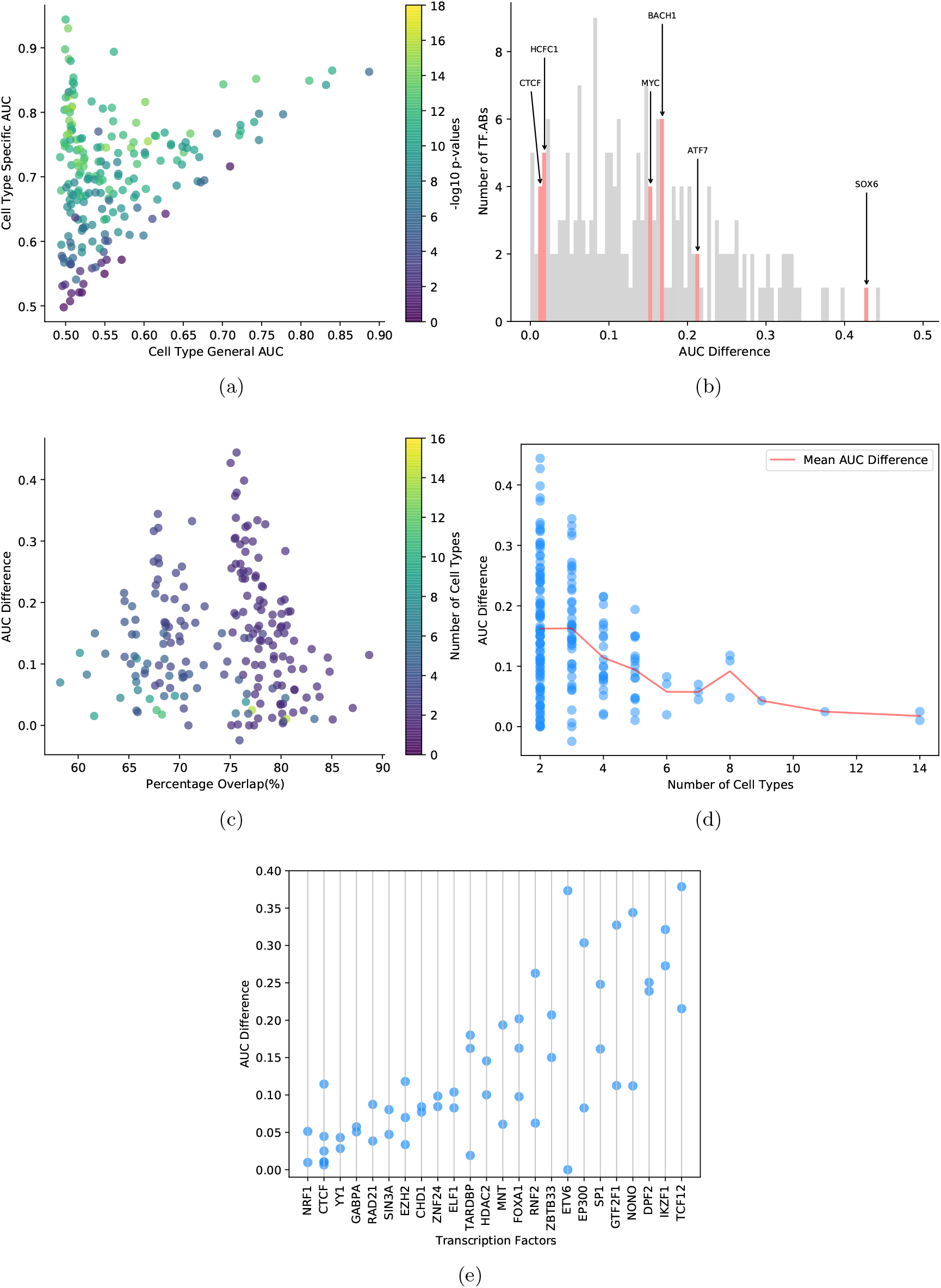
(a) Scatter plot of Cell Type General AUC versus Cell Type Specific AUC with the color gradient depending on the -log10 p-values. (b) Bar chart of AUC^1^differences. (c) Scatter plot of percentage overlap(%) and AUC differences, with the color gradient depending on number of cell types per TF-AB. (d) Scatter plot of number of cell types versus AUC difference, with red line depicting mean AUC difference per cell type count. (e) Scatter plot of AUC differences for TFs with more than two ABs.

Out of 230 TF-AB combinations, 204 TF-ABs have a p-value of less than 10^-5^, suggesting that a majority of TFs have some degree of cell type DNA signatures in their binding sites. We observe that many TFs have cell type specific binding patterns at differing degrees of specificity, where the majority of the points (approximately 143) show relatively little predictability in the cell type general formulation (Cell type general AUC<0.55). Of the 143, 134 have a statistically significant degree of cell type specificity, with 70 showing cell type specific AUC above 0.7. Figure 3b shows a histogram of the distribution of AUC differences between Cell Type General and Cell Type Specific across the different TF and AB combinations, with the bins containing ATF7, CTCF, HCFC1, BACH1, MYC and SOX6 highlighted in red. TFs that play a pivotal role in cancer either as oncogenes or suppressors, such as MYC [52, 53, 54, 55], BACH1 [56, 57, 58], ATF7 [45, 46, 47, 48], and SOX6 [59, 60], show a relatively higher cell type specificity than other TFs, such as CTCF [49, 50, 51] and HCFC1 [61, 62, 63], that are involved in chromatin regulation or other cellular processes. SI Table 1 shows the AUC differences obtained for each TF-AB.

As explained above, the lack of overlap between binding sites in different cell types is not evidence per se of any differential DNA signature. We next examined whether there is any association between the two. Figure 3c plots the mean percentage peak overlap versus the mean AUC difference for each TF-AB, with the color gradient depending on the number of cell types and each point being a TF-AB combination. No clear relationship between the two variables is seen (Spearman correlation r=-0.1). With either a high percentage overlap between 75% and 80% or a lower percentage overlap between 65% and 70%, the AUC difference ranges between 0.44 and −0.02. This suggests that mean percentage overlap is not an indicator of cell type specificity.

We also investigate whether the number of cell types available for a specific TF-AB pair contributes to the varying degrees of cell type specificity. With the cell type count per pair ranging from 2 to 14, the differences observed could be attributed to the number of cell types. A higher number of cell types may increase the likelihood of detecting a difference in AUC – as the condition that results in cell type specificity may more likely be there. Conversely, with more cell types, the number of false positives may increase, thus reducing the possibility of finding a difference. With less cell types, on the other hand, a difference in AUC may be a result of the quality of ChIP-seq experiments. The scatter plot in Figure 3d shows the relationship between the number of cell types and AUC difference, with the red line depicting the mean AUC difference per cell type count. For the 101 TF-ABs with exactly two cell types, the AUC differences varies between 0.44 and −0.02, suggesting that the degree of cell type specificity is not impacted by the number of cell types when the count is equal to two. Similar trends are observed with cell type counts less than six. However, with a higher number of cell type, there is a limited number of experiments available, making it impracticable to draw any conclusions.

Out of the 194 distinct TFs we studied, 24 were assayed with multiple ABs. Lack of consistency across ABs due to different off-target biases or binding affinities may impact the TF’s DNA signatures. Moreover, different ABs may have been used on different sets of cell types. Nevertheless, we may be reassured of the generality of our results if our measure of cell type specificity is consistent between different sets of experiments for the same TF. For most of the TFs, 170 out of 194, only 1 AB is used (SI Figure 1). However, there are 19 TFs with at least 2 ABs – all of which are polyclonal. Figure 3e shows a plot of the AUC differences between Cell Type Specific and Cell Type General for the 24 TFs with at least 2 ABs. Each point corresponds to a TF-AB combination, with TFs ordered by increasing average specificity. For example, CTCF was assayed with five different ABs, with a consistent result of very little cell type specificity – all five AUC differences are below 0.14. Conversely, several TFs show consistently high cell type specificity across multiple ABs, including: TCF12, SPI1, MNT, IKZF1 and DPF2. The least consistency is seen for the TF ETV6. Surprisingly, both datasets for ETV6 explore the same two cell lines GM12878 and K562, yet produce very differing results for cell type specificity: almost 0 for ENCAB000ABD and 0.37 for ENCAB997CJG. This may be due to differences in the actual ABs used, or could be a result of differences in the total number of peaks per dataset for each TF-AB combination.

### Determining the DNA signatures associated with cell type specificity for ATF7

Next, we look into the DNA signatures/motifs that cause the network to predict binding differently in general versus specific cell types for ATF7. Many approaches have been proposed for deriving important features from trained neural networks. In this section, we use several different methods to ensure consistency and for better interpretation of the model.

First, adopting a similar approach to AI-TAC [64], we use the t-SNE algorithm to present each ChIP-seq peak per cell type by its activation values in two dimensions across the 99 neurons of the final fully connected layer of the stage 2 model. Figures 4a and 4b show the ATF7 t-SNE plots for both cases of CL General and CL Specific respectively. Each point represents an instance, and is colored depending on which cell type to which it belongs. When predicting for general binding (Figure 4a), the DNA sequences, as encoded by the final network layer, appear as an undifferentiated mass with no obvious clustering structure, although the peaks from some cell lines do tend to be on one side or the other of the mass. Only bound instances per cell line are shown in Figure 4a. However, when CL information is provided, the network’s representation of the sequences groups them perfectly by cell type (Figure 4b). Similar trends were observed when applying t-SNE to other cell type specific TFs, such as CREM and SP1, as seen in SI Figures 3 and 4.

**Figure 4:**
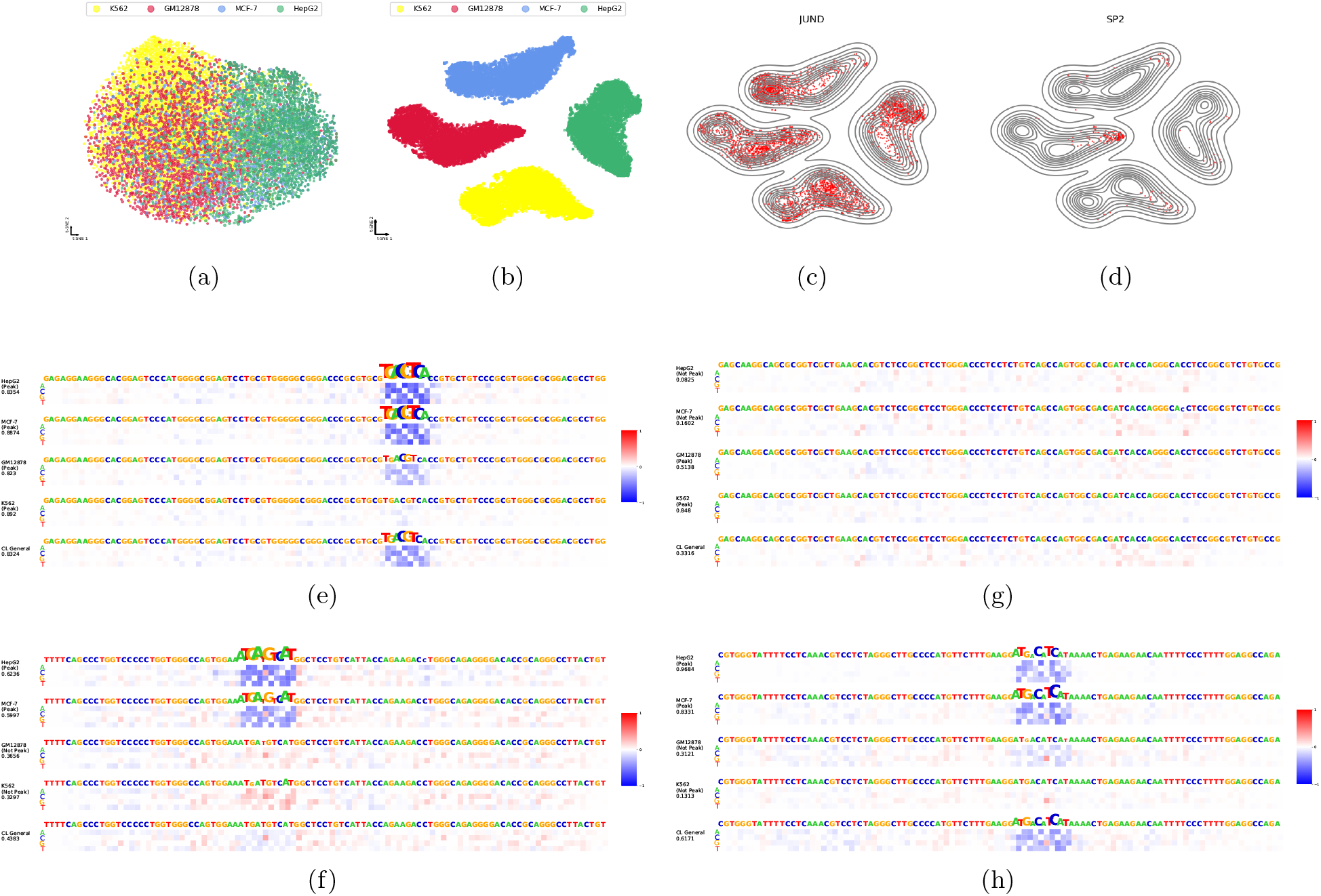
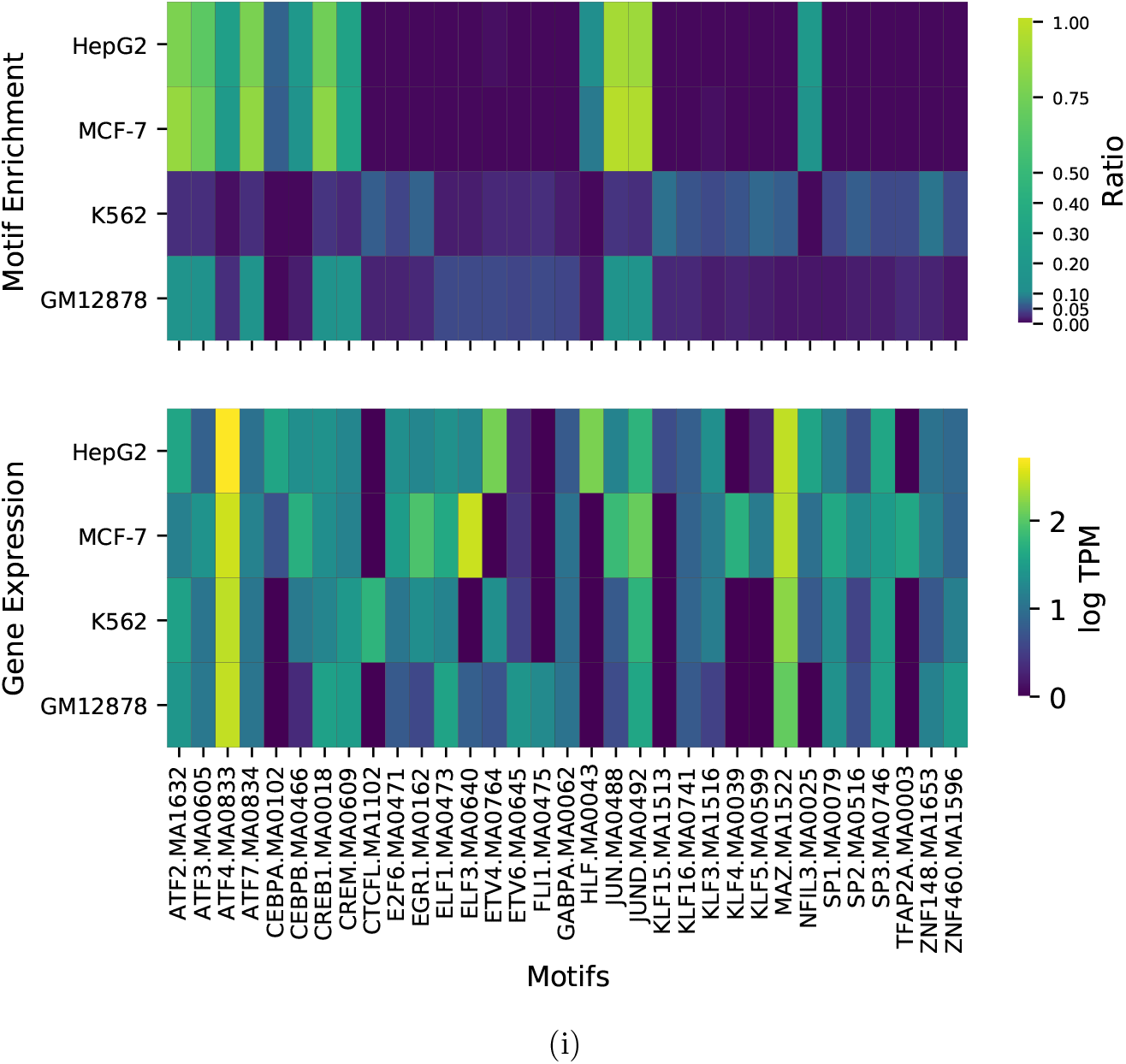
Motif enrichment for ATF7.ENCAB000BMO (a) t-SNE plot with cell type general concatenation. (b) t-SNE plot with cell type specific information. (c) and (d) are projections of the motifs JUND and SPI1 respectively onto the cell type specific t-SNE plots. (e), (f), (g) and (h) are mutation maps generated using in-silico mutagenesis on different test sequences. (i) Heatmaps of motif enrichment (top) and gene expression (bottom) per cell line across motifs.

We then explore the influence of the convolutional filters from the convolutional layer and their projections into this space. To do this, we converted the filters into PWMs and then used Tomtom [65] to search for the PWMs in the JASPAR database [35] (see Methods). Our emphasis was on filters with significant similarities to known motifs of TFs in JASPAR; we found that filters without significant matches usually captured partial or variant motifs for known TFs. We note that filters could have more than one significant motif match, however, we focused on the single best match for simplicity. For example, ≈40% of the filters matched best to the JUND motif, a basic leucine zipper factor. Figure 4c shows that the JUND motif is present in many bound sequences across the four cell lines, although it appears more enriched in subclusters of the MCF-7 and HepG2 peaks. ATF7 and JUND both have basic leucine zipper domains, and are known to physically interact [66]. Another set of filters matched the SP2 motif. Figure 4d shows this motif is found in many fewer peaks, and primarily a subgroup of GM12878 peaks. Notably, C’s and G’s are enriched in the SP2 binding site motifs in this cell type, although a link between SP2 and ATF7 has yet to be established.

To further explore the DNA signatures the network learns, we use another interpretation technique called in-silico mutagenesis (see Methods). We apply in-silico mutagenesis to the ATF7 test sequences to infer TF binding sites per cell type. We compute the differences in network output when altering each element of the DNA input to each other possible nucleotide, and obtain mutation maps per sequence per cell type for both cases of CL Specific and CL General. The white color in the mutation maps indicates no change, while blue or red refer to a drop or increase in predicted probability of binding respectively. Examples of mutation maps for different test sequences can be seen in Figures 4e, f, g, and h. The mutation maps labeled “CL General” visualize the influence of the perturbed nucleotides along the sequence with cell type general information, while those labeled with the cell type name refer to the cell type specific state. Below the cell type name in the mutation maps is an indication of whether or not the sequence is a peak in that specific cell type. The network-predicted peak probability is included below that.

The sequence in Figure 4e yields the same result for both cases of cell type specific and cell type general across all the four cell types. The mutation maps highlight the subsequence “TGACGTCA”, which is equivalent to the ATF7 motif profile in JASPAR. The network output is especially sensitive to the ATF7 motif when predicting for MCF-7 and HepG2, and less so for K562 and GM12878, although it still correctly predicts that peak for all cell types. Next, we focus on the sequence in Figure 4f. This sequence is a peak, and correctly predicted to be a peak, in MCF-7 and HepG2, and not a peak, and correctly predicted not to be a peak, in GM12878 and K562. The ATF2 motif of “ATGATGTCAT” is observed in HepG2 and MCF-7, but not detected as important in GM12878 and K562 nor cell type general. Indeed, altering some nucleotides in K562 and GM12878 actually increases the peak probability of the sequence. Other ATF7 motif variant patterns, such as “ATGACATCAT” and “TGATGCAAT”, are observed with cell specific information in SI Figures 2a and b respectively, where the motif pattern is clearly apparent in HepG2, barely apparent in MCF7, and non-existent in K562 and GM12878. Conversely, we observe alternate behavior in Figure 4g – where the sequence is considered to be a peak, and predicted to be a peak, in K562 and GM12878, and not a peak, and predicted not to be a peak, in HepG2 and MCF-7. No motif patterns are highlighted along the entire 101bp peak length, even though the network correctly predicts the peak for K562 and GM12878. It is not clear whether the network is predicting a peak by “default” and then suppressing that prediction in HepG2 and MCF-7 due to lack of a satisfactory motif or some competitive binding signal, or if there is some other multiply-present, redundant pattern that causes a positive prediction for GM12878 and K562–perhaps some co-binding factor with a motif very unlike the ATF family motifs.

To more systematically connect the important input sequence regions identified by in silico mutagenesis with known TF binding motifs, we extracted subsequences of length 31bp centered on the nucleotide with highest impact on prediction, and searched for enriched motifs using FIMO [67] (see Methods). Figure 4i (top) shows a heatmap of the ratio of the number significant motif hits to the total number of bound peaks per cell line per motif. We notice two main clusters of cell lines – each of which shows enrichment for a subset of motifs. As expected, the bZIP motifs, such as ATF2, ATF3, ATF4, ATF7, CREM and JUND are enriched in all four cell lines, with relatively more enrichment seen in HepG2 and MCF-7 as opposed to K562 and GM12878. Many TFs originate from TF families with similar DNA binding motifs, where motif enrichment is expected, but this does not necessarily denote involvement. Moreover, similar to the t-SNE output, we notice mild enrichment of SP2 (and other motifs, such as KLF3 and KLF4) in K562 and GM12878, and no enrichment in MCF-7 and HepG2.

In addition to scanning the test sequences using FIMO, we obtain the RNA-seq gene expression for the motifs across the four cell types (see Methods), as shown in Figure 4i (bottom). For many TFs, such as ATF4, CEBPA, CEBPB, CTCFL, JUN and JUND, there are similar patterns in both motif enrichment and gene expression – meaning motifs that are enriched across the four cell types are also expressed in the corresponding cell lines. However, there are TFs showing motif enrichment unrelated to expression. Results may be due to similar motifs showing enrichment for something not expressed (such as KLF4), co-expression with other TFs without binding, or competitive binding with other factors. SP2, for example, is expressed in HepG2 and MCF-7, but its motif is not enriched in those peaks.

### Determining the DNA signature associated with CTCF binding

We next explore the cell type specificity, or lack thereof, in CTCF binding. Similar to our previous analysis on ATF7, we first apply the t-SNE algorithm on the last fully connected layer produced per cell line from the pretrained stage 2 model. We compare both cases of CL General and CL Specific for the TF CTCF and AB ENCAB000AXX (Figures 5a and b). The plot in Figure 5a has no clear structure, similar to what was observed for ATF7. However, in Figure 5b, there are distinct clusters of DNA sequences – not from individual cell types, but from groups of cell types. For instance, the yellow/green/orange cluster towards the bottom comprises primarily peaks from smooth muscle cells, astrocytes, myotubes, and hepatocytes. Some clusters are divided into two parts. The cluster encompassing the yellow, green and orange points, for example, is found in two different locations in the t-SNE plot. This is also true for the top cluster of purple, blue and brown points. Some degree of cell type clustering is apparent, however the cell type groupings of sequences are not as clearly evident as in ATF7. Only positive instances per cell type are displayed in the plot, however. Thus, a separation of some positive instances does not necessarily indicate that the model is performing well on the negative instances.

**Figure 5:**
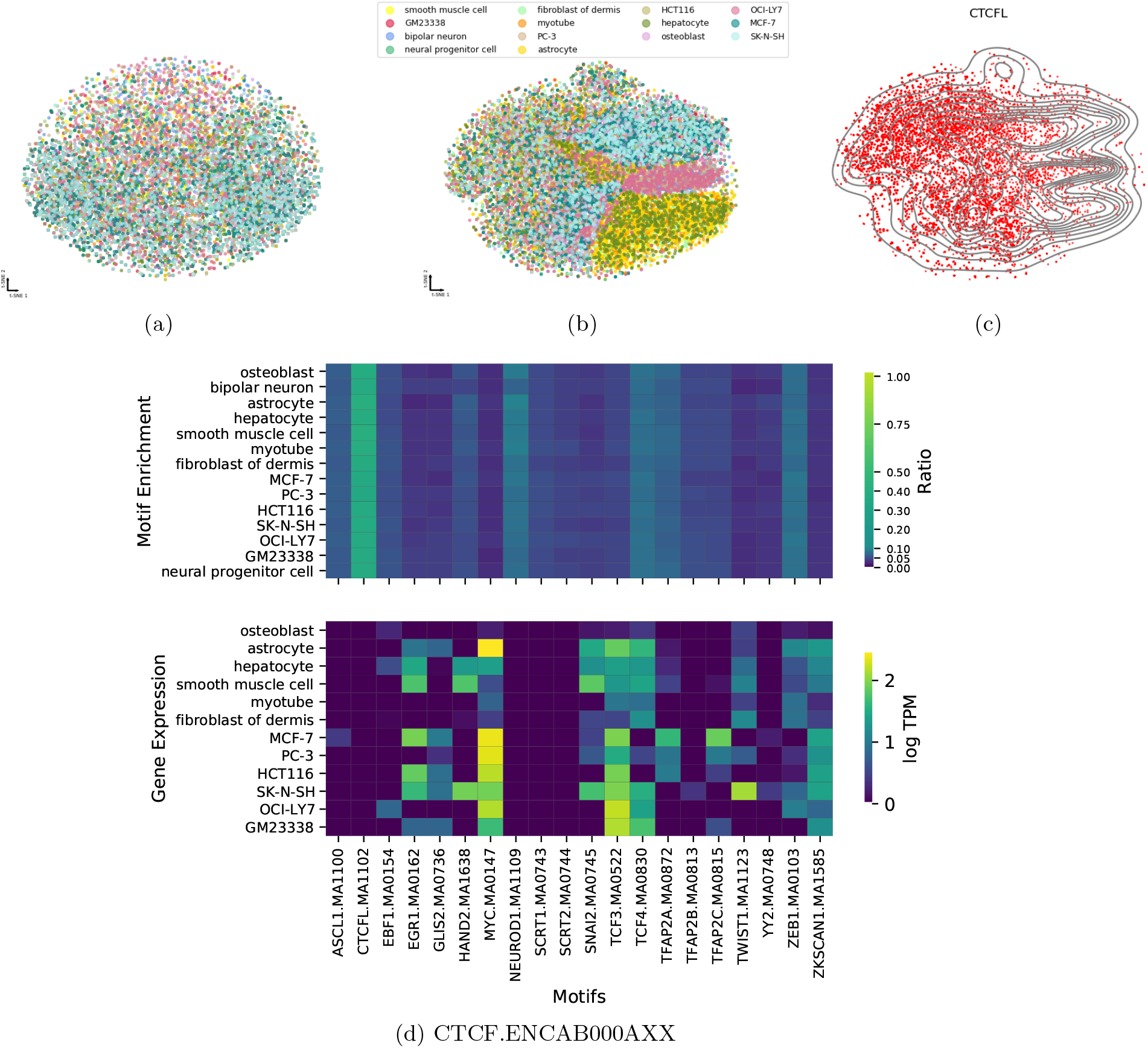
Motif enrichment for CTCF.ENCAB000AXX (a) t-SNE plot with cell type general concatenation. (b) t-SNE plot with cell type specific information. (c) Projections of CTCFL motif onto the cell type specific t-SNE plot. (d) Heatmaps of motif enrichment (top) and gene expression (bottom) per cell line across motifs.

We also examined the filters in the first convolutional layer of the network to discover what known TF PWMs they resembled. In Figure 5c, we show the non-uniform distribution of CTCFL. CTCFL (CCCTC-binding factor like) is a paralog of CTCF, and thus both share the highly conserved zinc finger DNA binding domain [68, 69]. However, unlike CTCF, CTCFL is known to be more cell type specific [68, 69]. We notice that CTCFL avoids the lower right yellow/green clusters and the upper center purple/blue/green cluster. The cell types in the yellow/green clusters include the smooth muscle cell, myotube, astrocyte and hepatocyte.

Next, we use in-silico mutagensis and FIMO to identify motifs enriched across the 14 cell types, as seen in Figure 5d (top). Unsurprisingly, different cell types show very similar patterns of motif enrichment, with the highest enrichment observed for CTCFL, and relatively high enrichment for other motifs, such as NEUROD1 and ZEB1. Additionally, similar to ATF7, we visualize gene expression per motif per cell line (Figure 5d (bottom)).

Analogous behavior is observed for CTCF using another AB ENCAB000AFR with a smaller number of cell types, as seen in SI Figure 5. With five cell types as opposed to 14, clustering using the cell type specific concatenation does not produce a distinct cluster for each cell type, rather it produces distinct subgroups of cell types that may be more cell type specific than others.

### Learned motifs across TF families

In this section, we examine the motif enrichment for various TF and antibodies (ABs). As previously discussed, we apply in-silico mutagenesis and FIMO on all TF-AB models (see Methods). TFs from the same families have identical DNA-binding domains [70, 71]. As a result, we focus on homologous TFs that are part of TF families, as provided by the hierarchical clustering of the DNA binding domains in the JASPAR database [43, 71, 72]. This gives a total of 261 TF-AB and cell type combinations.

Figure 6a visualizes the ratio of significant motif hits identified by FIMO to the total number of binding sites per TF-AB and cell type. The columns in the heatmap correspond to the ChIP-seq experiments in the form TF-AB-CL. The rows correspond to known motifs from JASPAR [35]. Each group of ChIP-seq experiments (columns) is part of a TF family, and is given a unique color as shown at the top of the heatmap. Different clusters of motifs are especially enriched in different clusters of ChIP-seq experiments (labeled A, B, C, D1, D2, E).

**Figure 6:**
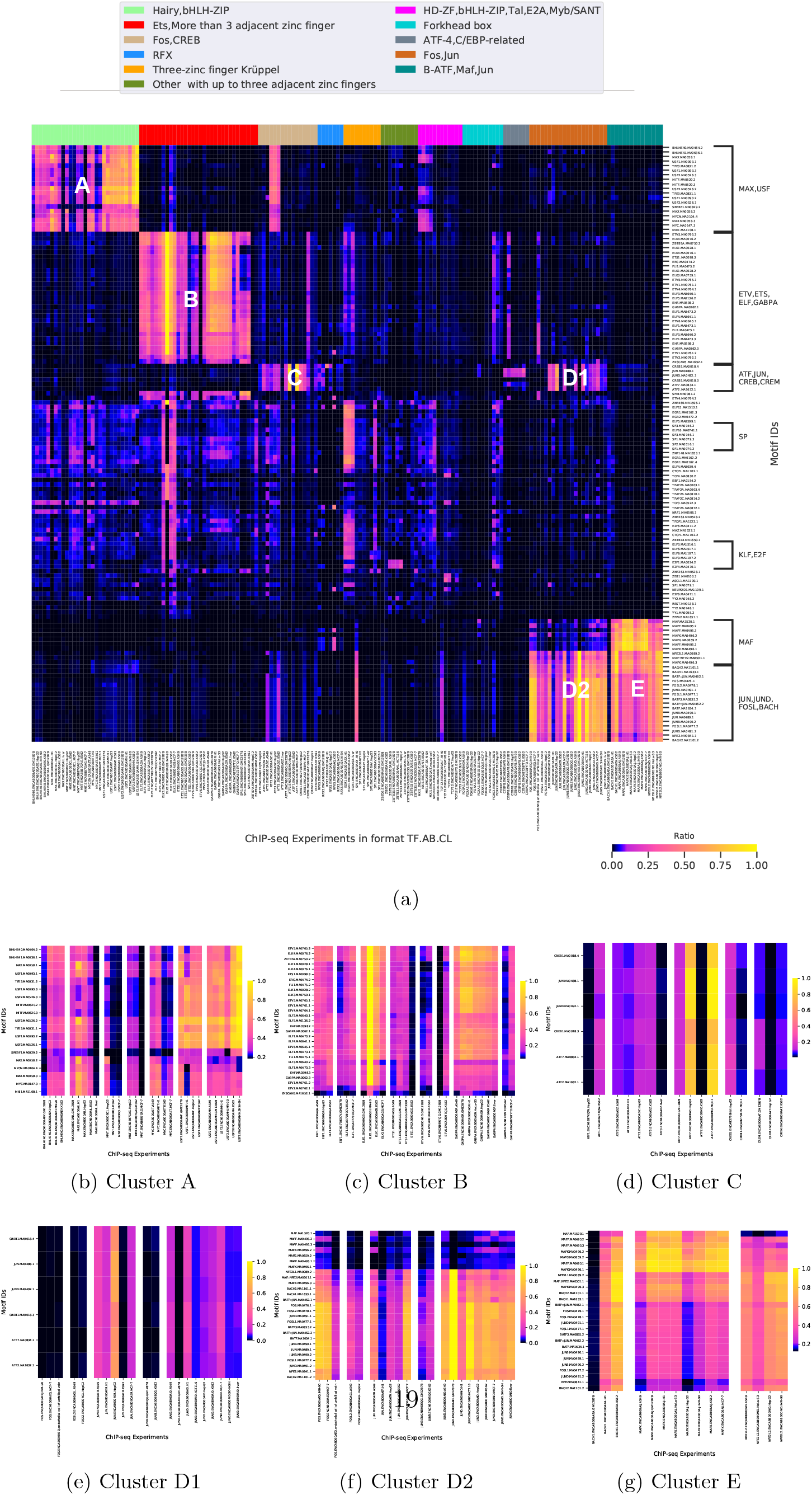
Overall motif enrichment. (a) Heatmap of motif enrichment. Each column corresponds to a TF-AB and cell type combination, and each row corresponds to a motif ID. Each TF family is given a unique color as seen at the top of the heatmap. (b), (c), (d), (e), (f) and (g) are zoomed in heatmaps for clusters A, B, C, D1, D2 and E.

To more clearly visualize the trends observed, Figures 6b-g display enlarged heatmaps of each cluster. Cluster A, for example, consists of the TFs BHLHE40, MAX, MNT, MYC, USF1 and USF2, which are part of the bHLH-ZIP and Hairy TF family. We notice uniform patterns of enrichment across motifs and cell types in USF1 and USF2, meaning the level of motif enrichment across their corresponding cell types does not change. This indicates a lack of cell type specificity, and is consistent with the low degrees of cell type specificity of less than 0.1 (SI Table 1). MAX, however, has a higher degree of cell type specificity of 0.15. This is due to the very low levels of enrichment motifs in liver, while the other four cell types have a higher and more constant level of motif enrichment. Similar behaviours can be seen for MNT with either antibody, and MYC. The TFs in this family, do not all show similar levels of enrichment across the motifs. For example, although the motif SREBF1 is enriched in all TFs, it is overall more enriched in BHLHE40, USF1 and USF2 in comparison to MAX, MNT and MYC. This may because the TFs MAX, MNT and MYX suppress SREBF1.

Similar trends of varying enrichment levels across different cell lines for the same TF and AB combination are also observed in the other clusters. For example, in cluster B for ETS1, we observe that most of the motifs (ELK1, ETV5, ETV4, GABPA, etc.) are enriched in the cell types A549, GM12878 and GM23338, but not in K562. As such, K562 has the highest difference in AUC between cell type specific (≈0.68) and cell type general (≈0.56) concatenation, in comparison to the other cell types.

The Fos,Jun TF family has two main clusters, D1 and D2. Cluster D2 (Figure 6f) shows varying levels of enrichment in most members of the Fos,Jun TF family across motifs from the MAF-related family, BACH, FOS/FOSL and JUN/JUND. For instance, the MAF motifs at the top of the heatmap show relatively little enrichment compared to the other motifs below, such as BACH1 and FOSL1, with some cell lines, such as JUN.ENCAB000AER.A549, having more enrichment for these motifs than others. Moreover, the cell lines, JUN.ENCAB000AER.H1 and JUNB.ENCAB000BQG.GM12878, have the lowest motif enrichment compared to other cell lines in the cluster. There are also observable differences between cells types for FOS, FOSL2 and JUN, which is consistent with the moderate level of cell type specificity obtained by SigTFB. Cluster D1 (Figure 6e), on the other hand, displays different trends for another set of motifs. Notably, JUN.ENCAB000AER.H1, which lacks enrichment for D2 motifs, shows significant enrichment for D1 motifs, including JUN and JUND motif variants that are different between blocks D1 and D2. Additionally, in all the cell types for FOS and FOSL2 in cluster D1, there is little to no enrichment for motifs CREB1, CREM, ATF7, ATF2 and ATF3.

Another group is cluster E, in Figure 6g, which consists of the TFs BACH1, MAFK and NFE2L2, part of the B-ATF,Jun,Maf TF family. For the ChIP-seq experiments MAFK and NFE2L2, we observe varying patterns of enrichment across their corresponding cell types. These results are consistent with the AUC differences obtained earlier of more than or equal to 0.1 – which indicates a higher degree of cell type specificity. MAFKs are part of the small MAF TF group, and are a member of the bZIP family. They are known for their role in gene expression regulation in multiple cellular processes. There is no sufficient evidence to suggest any cell type specific functionality, or lack that of, of MAFK [73]. Different, more apparent, trends are observed across cell types for BACH1. Cell types K562 and H1 show high levels of enrichment for all motifs, while cell type GM12878 displays no motif enrichment.

Ideally, ChIP-seq experiments with different ABs for the same TF identify similar peaks, and therefore similar DNA signatures in those peaks. Although Figure 6 shows consistency in motif enrichment across ABs for the same TF, exceptions to this notion do exist. Examples of consistent levels of enrichment in the same cell types for different ABs can be seen in MNT.ENCAB000BCL and MNT.ENCAB887GAG in Cluster A (Figure 6b), and ELF1.ENCAB000AGA and ELF1.ENCAB778OCV in Cluster B (Figure 6c). A lack of consistency in enrichment can be seen in Cluster B, however, more specifically in ETV6.ENCAB000ABD and ETV6.ENCAB997CJG for cell type GM12878. This can also seen in the drastic differences in AUC difference obtained for each of TF-AB, as discussed earlier.

Figure 7 provides a graphical representation of the most significant motif hits (ratio> 0.5) in cell types: A549, K562 and HepG2. Each node in the network represents a TF/motif. Directed arrows point from TFs representing ChIP-seq experiments (or TF-AB combinations) to known motifs enriched in that experiment. Enrichment values from different versions of the same motif are averaged. Arrows are colored according to which cell type(s) show enrichment. SI Figure 6 displays motif networks for other cell types (GM12878, MCF-7, liver, H1, HeLa-S3, HCT116, IMR-90, SK-N-SH).

**Figure 7:**
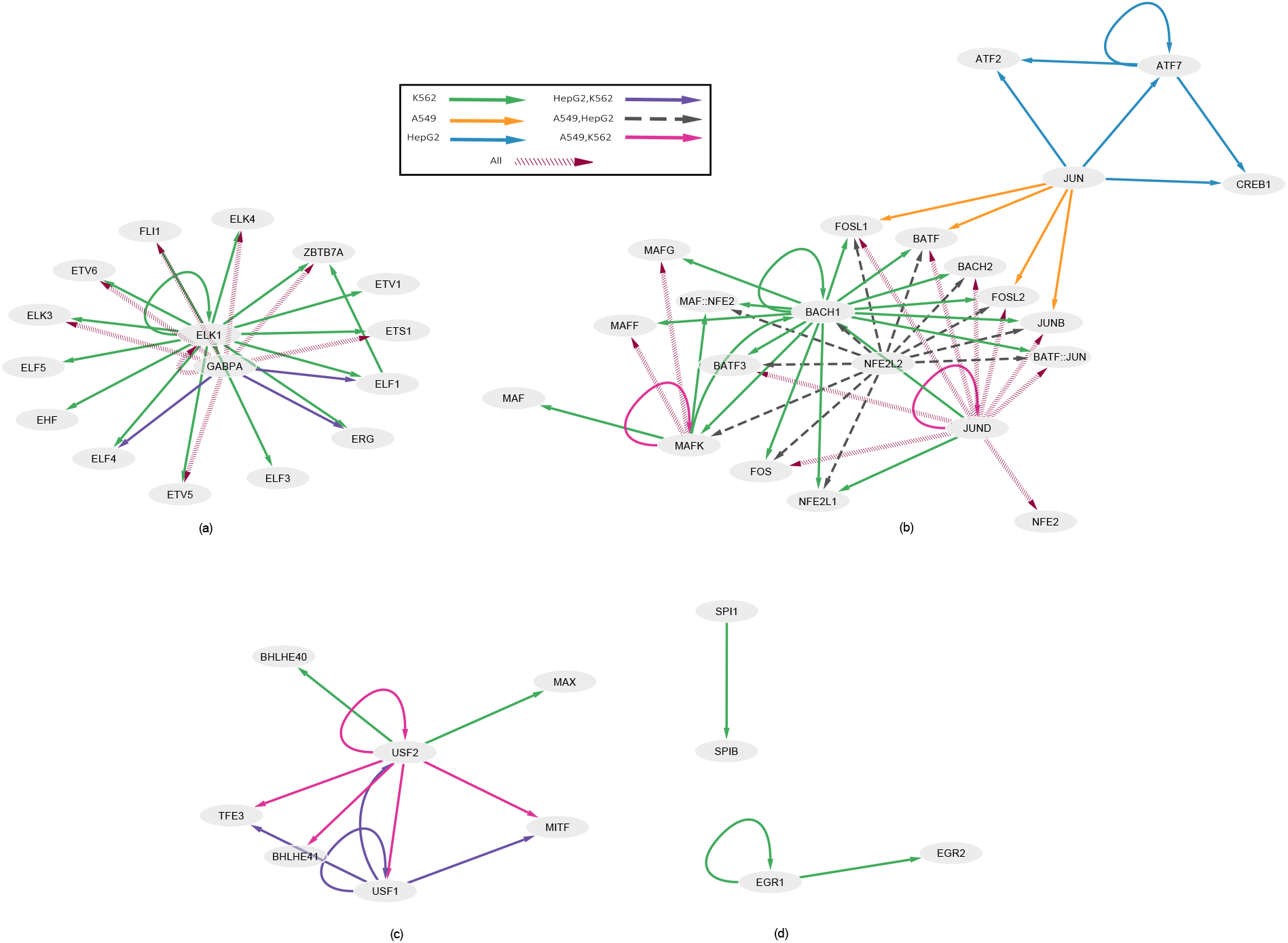
Motif enrichment network in the cancerous cell types: A549, K562 and HepG2. Each node represents a TF/motif, and the directed edges point from TFs of ChIP-seq experiments to known motifs enriched in that experiment. The various edge colors correspond to whether a motif is enriched in only one cell type, two cell types, or all three.

The three selected cell types are cancerous: A549 is lung carcinoma, HepG2 is hepatocellular carcinoma and K562 is leukemia. There are three main groups of TFs/motifs in Figure 7 (labeled a, b and c). Figure 7a, for example, is composed of TFs from the E26 transformation-specific (ETS) TF family. We notice enrichment of many ETS motifs, such as ETS1, ETV6 and ELK3, in all of three cell types for the GABPA TF (dashed maroon edges in Figure 7a). Abnormal levels of ETS motifs are often associated with cancer development and progression [74, 75]. Additionally, although many of the ETS motifs are enriched in all three cell types, ELF4, ELF1 and ERG are only enriched in K562 and HepG2, and not in A549 (purple edges in Figure 7a).

Figure 7b constitutes the FOS, JUN, MAF and ATF sub-families – part of the activator protein-1 (AP-1) TF family. We notice the enrichment of the AP-1 motifs: FOS, BATF3, FOSL1, BATF, FOSL2 and JUNB, in all three cell types for the JUND TF (dashed maroon edges in Figure 7b). AP-1 TFs play an important role in cancers and impact the proliferation, differentiation and apoptosis of cancerous cells [76]. While in most cancers such as breast [77, 78] and lung [79, 80] cancers, the AP-1 TFs act as oncogenes, in other cancers, such as leukemia, they may either act as oncogenes [81, 82] or tumor suppressors [83, 84, 85]. For instance, in spite of ChIP-seq data being available for the JUN TF in all cell types, we notice a difference in enrichment across the cell types. There is no significant enrichment of any motif for the JUN TF in K562, while in HepG2 and A549 the AP-1 TFs are enriched. This may be because JUN is suppressed in K562. Another example is BACH1, which shows enrichment for many of the AP-1 motifs, but only in K562.

Similar behaviours of variations in motif enrichment across cell types for a specific TF is shown in Figure 7c for the upstream stimulatory factors, USF1 and USF2. For example, the motifs TFE3, BHLHE41, USF1, USF2 and MITF are enriched in A549 and K562 for the TF USF2, while BHLHE40 and MAX are only enriched in K562 and not in A549. It is also important to keep in mind that ChIP-seq experiments may not be available for all three cell types. Thus, having enrichment only in one cell type may be due to ChIP-seq data being only available for that TF in that specific cell type, such as the case of TFs SPI1 and EGR1 in cell type K562 in Figure 7d.

## Discussion

The complex structural and biochemical nature of protein-DNA interactions has made it difficult to fully understand how various factors influence transcriptional regulation and differential binding. We conducted a wide-scale investigation of TF and AB combinations across various cell types to identify and quantify differential binding preferences of TFs. The “DNA signatures” constructed by our deep learning approach can account for many factors that could determine the binding preferences of a TF, such as its intrinsic binding preference, chromatin accessibility or co-binding factors. Different TFs display varying degrees of cell type specificity in their binding preference. For instance, our close examination of ATF7 found substantial cell type specificity, whereas we saw virtually none for CTCF. Such cell type specificity, or lack thereof, is also reflected in learned DNA sequence representations and motif enrichment analyses.

Differential binding analysis is one of the many key methods used for uncovering the relationship between TFs and the human genome in mechanisms, for example, leading to differential gene expression in tissue development or disease occurrence, or cooperative binding of TFs with other proteins and TFs. Binding site identification, or peak calling, of ChIP-seq can tell us whether a TF is binding the same or different sites in different cell types. However, we have shown here that peak overlap has negligible statistical association to the presence or absence of a DNA signature of differential binding.

Other deep learning approaches, such as MTTFSite [31] and Phuycharoen et al. [9], have also explored differential binding of TFs across cell types. While MTTFSite and Phuycharoen et al. adopt a similar learning framework to SigTFB in stage 1 training, in terms of using a multi-task model, their problem formulation and objective fundamentally differ. In MTTFSite, for example, prior to training, shared non-unique cell type instances are defined as bound regions across cell types that overlap by at least 100bp, while the remaining bound instances that do not overlap are cell type specific. In SigTFB, however, the model is given all instances as input and learns to differentiate non-specific versus cell type specific instances. The negative instances for a specific cell type in SigTFB are bound regions in other cell types, while in MTTFSite and Phuycharoen et al. negative instances are unbound regions in all cell types. SigTFB essentially learns to differentiate between shared and unique motifs in cell types from only bound regions. Additionally, the scale of the study differs. MTTFSite and Phuycharoen et al. investigate TFs in a total of 5 and 3 cell types respectively, while SigTFB explores all TFs in ENCODE with at least more than 2 cell types available, resulting in a total of 35 cell types across all TFs. Moreover, while MTTFSite trains a shared multi-task model on all cell types, then evaluates the network on previously defined cell type specific features, SigTFB trains a private network for each cell type in stage 2, using the weights from pretrained multi-task model of stage 1, to learn the cell type specific features of a cell type via the concatenation of one-hot cell type specific encoding. Through this approach, SigTFB demonstrates the varying degrees of TF specificity across cell types on wide scale, and shows the common non-specific preferences of a TF, as well as the unique cell type specific ones.

Similar to Novakovsky et al [43] and ChromDragoNN [42], we display the effectiveness of transfer learning in a multi-task deep learning framework for the prediction of binding profiles genome wide. Unlike these approaches, however, which mainly focus on cross cell type prediction, where models are trained on some cell types and tested on other cell types with limited data, we use transfer learning to acquire exclusive features per cell type. The multi-task setting in the first stage of learning allows the model to learn generalizable shared and unique features across cell types. Then, with the use of transfer learning, the model is constrained to learn cell type specific features, allowing the learning of a set of motifs that cause cell type specificity. In addition to the type of learning used, data representation, the criteria chosen for model evaluation, and the hyperparameters selected are important factors we account for during the learning phase to achieve a more accurate prediction of binding profiles at cell type resolution.

Most deep learning approaches, such as DeepBind [23], MTTFSite [31] and DanQ [25], do not investigate the differences in ABs for the same TF when analyzing ChIP-seq experiments. We hypothesized that ABs could greatly influence the quality of ChIP-seq experiments. The polyclonal nature of ABs in ENCODE, for example, may result in ABs targeting the same protein to have different specificities, affinities and off-target binding. As a result, due to the lack of study on the consistency of functionality and performance of ABs across TFs ENCODE wide, we separate experiments from different ABs for a particular TF, and investigate the consistency in binding preference across different ABs for the same TF. Overall, we find consistency across ABs for most TFs (≈ 88%), while for others the consistency is less apparent. Still, any lack of consistency in output may be due to other factors, such as the quality of the ChIP-seq dataset, the controls selected for peak calling, or the cell types available.

The fundamental reasoning behind our study is that, by understanding how a deep neural network can code for cis-regulatory regions in differentially bound cell types, we can deduce the extent of cell type specificity of various regulatory proteins, and ultimately study their impact on downstream genomic analysis. Nonetheless, we acknowledge some limitations. First, the fact that a TF does not show cell type specificity in the cell types available from ENCODE does not imply that it will not show cell type specificity in other cell types. The human genome contains almost 1400 TFs [86], and despite the enormous effort of the ENCODE consortium, we found only 194 distinct TFs assayed in more than one cell type meeting our data set criteria. It is thus impossible to detect cell type specific binding for the vast majority of TFs, and it is uncertain whether other TFs may show specificity in other cell types. This underlines the importance of continued empirical study of TF binding in a wide range of cell types. A second limitation is that, despite best efforts, deep learning can at times fail to solve a prediction problem, even when a solution is possible in principle. There may be TFs for which we failed to detect a cell type specific signal, even when one is present. On the other hand, our careful checks against overfitting suggest that when a cell type specific signal is present, it is likely genuine, especially when it is backed up by additional motif enrichment analyses. Thus, our results are best viewed as providing evidence for cell type specific DNA signatures in many TFs, while providing evidence against the same, without ruling it out, for other TFs. Thirdly, assumptions made regarding the network architecture, such as the 101 bp input sequence or fixed filter widths, may limit the learning capabilities of SigTFB. For instance, its inability to detect widely spaced motifs or motif pairs with fixed spacing, suggests that some DNA signatures relevant for cell type specificity may be possibly missing. Furthermore, in this work, we use ChIP-seq data due to its high availability and accessibility for multiple TFs and cell types. While ENCODE has many standards in place to ensure high data quality, other experimental approaches, such native ChIP [87] or ChIP-exo [88], may provide less noisy, higher resolution and more precise estimates of TF-DNA binding, and thus may ultimately improve the search for DNA signatures. Finally, it is important to note that our confirmatory motif enrichment analysis is limited by the current state of knowledge. Like ENCODE, JASPAR includes data on a relatively small fraction of all human TFs. Not all motifs are in JASPAR, and not having matches may not necessarily be because there are not any significant matches, but because there are not any matches in the JASPAR database for this motif, which emphasizes the importance of further effort in that domain.

While the degree of cell type specificity may be inferred from the TF-AB models, one can not directly infer the cause of such results. Thus, future work may further explain the rational behind such cell type specificity with a more in depth analysis of the different factors that cause cell type specificity in TF binding. Our study provides the starting point for such more detailed mechanistic investigations.

## Conclusion

Many TFs are known to bind to different genomic sites in different cell types. Here, we demonstrated that for some of these TFs, different binding sites are associated with different DNA signatures, while others are not. We developed a deep learning prediction framework that is capable of detecting such DNA signatures, and used explanation techniques to elucidate signatures from the trained networks. Our results provide a basis for future research into more specific molecular mechanisms that are reflected by these signatures.

## Methods

### Data accession and preprocessing

To identify the regions of enrichment along the genome for various TFs, we turned to the ENCODE project [2]. Because different antibodies for a TF can have different specificities or biases, we chose not to mix data from different antibodies. We identified 230 transcription factor-antibody (TF-AB) combinations that were generated using the ENCODE uniform processing pipeline for replicated ChIP-seq experiments [2]. We required experiments with data in at least two cell types. The pipeline maps pair-ended reads for each ChIP-seq experiment to GRCh38 and obtains filtered alignments. It then takes as input the filtered alignment BAM files from the replicates and the controls of the ChIP-seq experiment to detect peaks that pass the irreproducible discovery rate (IDR) at a threshold of 2% [2]. We used these peaks as input for our analyses.

For each TF-AB combination, we obtained the “experiment” level peaks across its various corresponding cell types. Then, similar to the greedy approach used by Basset [28] for chromatin accessibility, we repeatedly merged peaks from different experiments across various cell types if they overlapped by at least 30bp. The center of each merged peak is taken and extended by 50bp in each direction, such that the length of the intervals are 101bp [28]. There can be multiple experiments for the same TF-AB combination.

### Training and test set construction

We train a different model for each of the 230 TF and AB combinations (see Figure 1). In stage one, described further below, the dataset per TF-AB pair is a multi-class dataset, where each of the C cell types assayed for that TF-AB is one of the possible classes. Each instance corresponds to a unified peak, as described in the previous section. A positive instance is a sequence which is bound in a specific cell type by the TF under study, meaning they are regions that have passed the IDR threshold of 2%. Due to the merging of the peaks, a negative instance is a sequence not bound by that TF in a specified cell type. Because there can be multiple experiments for the same TF-AB pair in the same cell type, the target output is deemed positive if a peak was detected for at least one of the experiments in that cell type. The output is a binary vector of length C encoding which cell types the unified peak is bound, or not bound, in. There is at least one element that is bound (has a value of 1) in the output vector, and as many as all the elements bound, if that peak is found in all cell types.

For stage 2 training, there are 2 × *C* instances per unified peak. For *C* of those instances, the input is a one-hot encoded DNA sequence representing the unified peak, as well as a binary vector one-hot encoding one of the *C* cell types. The output is 1 or 0 depending on whether that instance is bound in that cell type or not. This is the cell type specific case, where the model is informed about which cell type is being explored. For the remaining *C* instances, a vector of all zeros is used, instead of a one-hot encoded vector, with the same output and DNA sequence as before. This is the cell type general case, where we are not informing the model which cell type we are investigating. Here, in the concatenation step, we use a direct encoding of cell type, instead of the gene expression values used in ChromDragoNN [42].

We split each TF-AB dataset into 90% training and 10% testing. At both stages 1 and 2, the training dataset is then repeatedly divided into training and validation at a ratio of 8:2. At each iteration (per stage), we train the model on the training subset and observe the performance on the validation subset. We use Ax for optimal hyperparameter selection [44]. This is repeated 10 times for each stage for each TF-AB dataset. The hyperparameters for stage 1 training includes learning rate, weight decay, initial weight scales for the convolutional and fully connected layers, number of channels, batch size, and the number of epochs. PyTorch 1.5.0 (GPU) with the Adam optimizer is used for stage 1 training.

Many datasets have an unequal distribution of instances across cell types for a specific TF-AB. As a result, we observe poor predictive performance for the minority cell types – although the overall predictive performance across all cell types for that model may be high. To address the class imbalance issue, we develop a multi-label balanced sampler. During training or testing, each mini-batch selected has the same number of total instances per cell type, and each cell type has the same number of positive and negative instances. Thus, we ensure that high overall predictive performance is not biased towards certain cell types because of class imbalance across or within cell lines. Models with very poor predictive performance per cell type per TF-AB (with specificity or sensitivity less than 0.2), even after addressing the class imbalance issue, are filtered out.

We also filter out ChIP-seq experiments with fewer than 100 positive or 100 negative test sequences. If the size of the dataset is too small, then enrichment values may be too optimistic.

### Neural network training

We formulate the training of differential binding per TF as a two step process (see Figure 1).

#### Stage 1

Stage 1 is formulated as a multi-task classification problem, such that for each input DNA sequence we predict whether or not its bound across multiple cell lines for a specific TF-AB. We use a modified version of DeepBind [23] of one hidden layer CNN with 404 binary inputs followed by a fully connected layer (Figure 1). Unlike DeepBind [23], the number of channels in our model is set as a hyperparameter. We also investigated the use of more complex models with more than one convolutional layer. However, the one convolutional layer gave the best results in terms of validation accuracy and loss. We use the Basset loss function, which was adopted by Basset [28] and ChromDragoNN [42], both of which predict chromatin accessibility across various cell types. Moreover, the 230 datasets obtained, with a total of of 194 unique TFs, are those that passed the filtering step in stage 1. That is, we discarded 20 datasets with a specificity or sensitivity of less than 0.2 in stage 1.

#### Stage 2

We then adopt a transfer learning multi-modal approach for stage 2, inspired by ChromDragoNN [42] (see Figure 1). In this stage, not only do we consider the genomic sequence of the peak, we also include cell type specific and cell type general information as input to the model. In stage 2 training, the genomic sequence is first input to the convolutional layer from stage 1, which is initialized using weights of the best pre-trained model with the optimized hyperparameters from stage 1. The convolutional layer from the stage 1 model is then concatenated with the output generated by the fully connected layer for either the cell type specific or cell type general data (refer to “CL General OR CL Specific” in Figure 1). The concatenated vector is then passed through a fully connected layer. Finally, the output node then determines whether or not this genomic sequence is bound in that specific cell type.

The negative log likelihood loss function with the SGD optimizer is used for stage 2. We use the same multi-label balanced sampler in stage 1 to address the class imbalance issue. Moreover, in addition to the stage 1 hyperparameters, stage 2 hyperparameters include number of neurons per layer before concatenation, number of neurons per layer after concatenation, freeze pretrained model, dropout probability and momentum rate. We use Ax for hyperparameter tuning [44]. The area under precision-recall curve (AURPC) is used to evaluate the primary performance of the stage 2 models.

### Feature attribution with in-silico mutagensis and FIMO

We apply in-silico mutagensis on test sequences per cell type for each of the 230 TF-AB pre-trained models. Using in-silico mutagensis, we change each nucleotide to every other possible value and observe the output of each perturbed sequence. From the mutation map per sequence, we then extract a 31bp subsequence with the largest negative impact on output.

To analyze the subsequences per cell type per TF-AB model, we then use FIMO 5.0.3 to search for motifs in the subsequences using known JASPAR human motifs [35] that are based on at least 1000 sites and had log p-values of at least 100. This gives us a total of 400 JASPAR motifs. For each cell type per TF-AB, and each motif, we find the ratio of the number of significant motif hits identified by FIMO to the number of total peaks for that cell type. We then find the top 20 motifs with the highest ratio. By using this approach, we account for enrichment as well as the number of peaks per cell type per TF-AB. We finally take the union of the these motifs across the cell types to construct the enrichment and expression heatmaps in Figures 4, 5 and 6.

Moreover, we obtain RNA-seq expression for the 35 distinct cell types across the 230 TF-AB pairs. The same cell type may appear in more than one TF-AB pair. We extract gene level expression data, both polyA and total RNAseq, in the form of transcripts per million (TPM) per cell type from ENCODE [2] – data that was generated using the uniform processing pipeline on unperturbed cell types. In the event that there is more than one RNA-seq experiment, be it polyA or total RNA-seq, available per cell type, the average TPM per gene per cell type is taken.

### Software availability

The source code for SigTFB involving the pre-processing of ChIP-seq data, classification and downstream genomic analysis is available at https://github.com/aawdeh/SigTFB.

## Acknowledgments

We thank members of the Perkins lab, and Compute Canada for granting us access to their cluster to store data and run our computational analyses.

## Authors’ contributions

AA and TJP conceived and designed the analysis. AA developed the tool, performed analysis/computations and wrote the manuscript with input from TJP. TJP edited the manuscript. TJP and MT supervised the project. All authors provided critical feedback and helped shape the research, analysis and manuscript. All authors read and approved the final manuscript.

## Funding

We acknowledge the support of the Natural Sciences and Engineering Research Council of Canada (NSERC), [funding reference number RGPIN-2019-06604]. This research was enabled by support provided by a Queen Elizabeth II Graduate Scholarship in Science and Technology (QEII-GSST) to AA, and a Compute Canada (www.computecanada.ca) Resources-for-Research-Groups grant to TJP. None of the agencies that funded this work had any role in the design of the study, in the collection, analysis, and interpretation of data, or in writing the manuscript.

## Availability of data and materials

The data used is available at https://doi.org/10.20383/103.0605.

## Ethics approval and consent to participate

Not applicable.

## Consent for publication

Not applicable.

## Competing interests

The authors declare that they have no competing interests.

## Notes

https://doi.org/10.20383/103.0605

https://github.com/aawdeh/SigTFB

### Competing Interest Statement

The authors have declared no competing interest.

## References

[1] Mattick, J.S.: Non-coding RNAs: the architects of eukaryotic complexity. EMBO reports 2(11), 986–991 (2001)

[2] Consortium, E.P., et al.: An integrated encyclopedia of DNA elements in the human genome. Nature 489(7414), 57 (2012)

[3] Srivastava, D., Mahony, S.: Sequence and chromatin determinants of transcription factor binding and the establishment of cell type-specific binding patterns. Biochimica et Biophysica Acta (BBA)-Gene Regulatory Mechanisms 1863(6), 194443 (2020)

[4] Lee, B.-K., Bhinge, A.A., Battenhouse, A., McDaniell, R.M., Liu, Z., Song, L., Ni, Y., Birney, E., Lieb, J.D., Furey, T.S., et al.: Cell-type specific and combinatorial usage of diverse transcription factors revealed by genome-wide binding studies in multiple human cells. Genome research 22(1), 9–24 (2012)

[5] Gertz, J., Savic, D., Varley, K.E., Partridge, E.C., Safi, A., Jain, P., Cooper, G.M., Reddy, T.E., Crawford, G.E., Myers, R.M.: Distinct properties of cell-type-specific and shared transcription factor binding sites. Molecular cell 52(1), 25–36 (2013)

[6] Heinz, S., Romanoski, C.E., Benner, C., Glass, C.K.: The selection and function of cell type-specific enhancers. Nature reviews Molecular cell biology 16(3), 144–154 (2015)

[7] Porcelli, D., Fischer, B., Russell, S., White, R.: Chromatin accessibility plays a key role in selective targeting of HOX proteins. Genome biology 20(1), 1–19 (2019)

[8] Bridoux, L., Zarrineh, P., Mallen, J., Phuycharoen, M., Latorre, V., Ladam, F., Losa, M., Baker, S.M., Sagerstrom, C., Mace, K.A., et al.: HOX paralogs selectively convert binding of ubiquitous transcription factors into tissue-specific patterns of enhancer activation. PLoS genetics 16(12), 1009162 (2020)

[9] Phuycharoen, M., Zarrineh, P., Bridoux, L., Amin, S., Losa, M., Chen, K., Bobola, N., Rattray, M.: Uncovering tissue-specific binding features from differential deep learning. Nucleic acids research 48(5), 27–27 (2020)

[10] Wang, J., Zhuang, J., Iyer, S., Lin, X., Whitfield, T.W., Greven, M.C., Pierce, B.G., Dong, X., Kundaje, A., Cheng, Y., et al.: Sequence features and chromatin structure around the genomic regions bound by 119 human transcription factors. Genome research 22(9), 1798–1812 (2012)

[11] Neph, S., Vierstra, J., Stergachis, A.B., Reynolds, A.P., Haugen, E., Vernot, B., Thurman, R.E., John, S., Sandstrom, R., Johnson, A.K., et al.: An expansive human regulatory lexicon encoded in transcription factor footprints. Nature 489(7414), 83–90 (2012)

[12] Zhang, S., Bell, E., Zhi, H., Brown, S., Imran, S.A., Azuara, V., Cui, W.: OCT4 and PAX6 determine the dual function of SOX2 in human ESCs as a key pluripotent or neural factor. Stem cell research & therapy 10(1), 1–14 (2019)

[13] Zaret, K.S., Carroll, J.S.: Pioneer transcription factors: establishing competence for gene expression. Genes & development 25(21), 2227–2241 (2011)

[14] Srivastava, D., Aydin, B., Mazzoni, E.O., Mahony, S.: An interpretable bimodal neural network characterizes the sequence and preexisting chromatin predictors of induced transcription factor binding. Genome biology 22(1), 1–25 (2021)

[15] Vierstra, J., Lazar, J., Sandstrom, R., Halow, J., Lee, K., Bates, D., Diegel, M., Dunn, D., Neri, F., Haugen, E., et al.: Global reference mapping of human transcription factor footprints. Nature 583(7818), 729–736 (2020)

[16] Filtz, T.M., Vogel, W.K., Leid, M.: Regulation of transcription factor activity by interconnected post-translational modifications. Trends in pharmacological sciences 35(2), 76–85 (2014)

[17] Brand, M., Ranish, J.A., Kummer, N.T., Hamilton, J., Igarashi, K., Francastel, C., Chi, T.H., Crabtree, G.R., Aebersold, R., Groudine, M.: Dynamic changes in transcription factor complexes during erythroid differentiation revealed by quantitative proteomics. Nature structural & molecular biology 11(1), 73–80 (2004)

[18] Nie, Y., Shu, C., Sun, X.: Cooperative binding of transcription factors in the human genome. Genomics 112(5), 3427–3434 (2020)

[19] Lopez, A.J.: Developmental role of transcription factor isoforms generated by alternative splicing. Developmental biology 172(2), 396–411 (1995)

[20] Lowen, M., Scott, G., Zwollo, P.: Functional analyses of two alternative isoforms of the transcription factor pax-5. Journal of Biological Chemistry 276(45), 42565–42574 (2001)

[21] Pilpel, Y., Sudarsanam, P., Church, G.M.: Identifying regulatory networks by combinatorial analysis of promoter elements. Nature genetics 29(2), 153–159 (2001)

[22] Banerjee, N., Zhang, M.Q.: Identifying cooperativity among transcription factors controlling the cell cycle in yeast. Nucleic acids research 31(23), 7024–7031 (2003)

[23] Alipanahi, B., Delong, A., Weirauch, M.T., Frey, B.J.: Predicting the sequence specificities of DNA-and RNA-binding proteins by deep learning. Nature biotechnology 33(8), 831–838 (2015)

[24] Hassanzadeh, H.R., Wang, M.D.: Deeperbind: Enhancing prediction of sequence specificities of DNA binding proteins. In: 2016 IEEE International Conference on Bioinformatics and Biomedicine (BIBM), pp. 178–183 (2016). IEEE

[25] Quang, D., Xie, X.: DanQ: a hybrid convolutional and recurrent deep neural network for quantifying the function of DNA sequences. Nucleic acids research 44(11), 107–107 (2016)

[26] Chen, C., Hou, J., Shi, X., Yang, H., Birchler, J.A., Cheng, J.: DeepGRN: prediction of transcription factor binding site across cell-types using attention-based deep neural networks. BMC bioinformatics 22(1), 1–18 (2021)

[27] Zhou, J., Troyanskaya, O.G.: Predicting effects of noncoding variants with deep learning–based sequence model. Nature methods 12(10), 931–934 (2015)

[28] Kelley, D.R., Snoek, J., Rinn, J.L.: Basset: learning the regulatory code of the accessible genome with deep convolutional neural networks. Genome research 26(7), 990–999 (2016)

[29] Quang, D., Xie, X.: FactorNet: a deep learning framework for predicting cell type specific transcription factor binding from nucleotide-resolution sequential data. Methods 166, 40–47 (2019)

[30] Li, H., Quang, D., Guan, Y.: Anchor: trans-cell type prediction of transcription factor binding sites. Genome research 29(2), 281–292 (2019)

[31] Zhou, J., Lu, Q., Gui, L., Xu, R., Long, Y., Wang, H.: MTTFsite: cross-cell type TF binding site prediction by using multi-task learning. Bioinformatics 35(24), 5067–5077 (2019)

[32] Li, H., Guan, Y.: Fast decoding cell type–specific transcription factor binding landscape at single-nucleotide resolution. Genome Research 31(4), 721–731 (2021)

[33] Qin, Q., Feng, J.: Imputation for transcription factor binding predictions based on deep learning. PLoS computational biology 13(2), 1005403 (2017)

[34] Matys, V., Kel-Margoulis, O.V., Fricke, E., Liebich, I., Land, S., Barre-Dirrie, A., Reuter, I., Chekmenev, D., Krull, M., Hornischer, K., et al.: TRANSFAC® and its module TRANSCompel®: transcriptional gene regulation in eukaryotes. Nucleic acids research 34(suppl_1), 108–110 (2006)

[35] Castro-Mondragon, J.A., Riudavets-Puig, R., Rauluseviciute, I., Berhanu Lemma, R., Turchi, L., Blanc-Mathieu, R., Lucas, J., Boddie, P., Khan, A., Manosalva Pérez, N., et al.: Jaspar 2022: the 9th release of the open-access database of transcription factor binding profiles. Nucleic acids research 50(D1), 165–173 (2022)

[36] Kulakovskiy, I.V., Vorontsov, I.E., Yevshin, I.S., Sharipov, R.N., Fedorova, A.D., Rumynskiy, E.I., Medvedeva, Y.A., Magana-Mora, A., Bajic, V.B., Papatsenko, D.A., et al.: HOCO-MOCO: towards a complete collection of transcription factor binding models for human and mouse via large-scale ChIP-seq analysis. Nucleic acids research 46(D1), 252–259 (2018)

[37] Stormo, G.D., Zhao, Y.: Determining the specificity of protein–DNA interactions. Nature Reviews Genetics 11(11), 751–760 (2010)

[38] Jolma, A., Yan, J., Whitington, T., Toivonen, J., Nitta, K.R., Rastas, P., Morgunova, E., Enge, M., Taipale, M., Wei, G., et al.: DNA-binding specificities of human transcription factors. Cell 152(1-2), 327–339 (2013)

[39] Badis, G., Berger, M.F., Philippakis, A.A., Talukder, S., Gehrke, A.R., Jaeger, S.A., Chan, E.T., Metzler, G., Vedenko, A., Chen, X., et al.: Diversity and complexity in DNA recognition by transcription factors. Science 324(5935), 1720–1723 (2009)

[40] Rohs, R., Jin, X., West, S.M., Joshi, R., Honig, B., Mann, R.S.: Origins of specificity in protein-DNA recognition. Annual review of biochemistry 79, 233 (2010)

[41] Jolma, A., Kivioja, T., Toivonen, J., Cheng, L., Wei, G., Enge, M., Taipale, M., Vaquerizas, J.M., Yan, J., Sillanpää, M.J., et al.: Multiplexed massively parallel SELEX for characterization of human transcription factor binding specificities. Genome research 20(6), 861–873 (2010)

[42] Nair, S., Kim, D.S., Perricone, J., Kundaje, A.: Integrating regulatory DNA sequence and gene expression to predict genome-wide chromatin accessibility across cellular contexts. Bioinformatics 35(14), 108–116 (2019)

[43] Novakovsky, G., Saraswat, M., Fornes, O., Mostafavi, S., Wasserman, W.W.: Biologically relevant transfer learning improves transcription factor binding prediction. Genome Biology 22(1), 1–25 (2021)

[44] Balandat, M., Karrer, B., Jiang, D.R., Daulton, S., Letham, B., Wilson, A.G., Bakshy, E.: Botorch: Programmable bayesian optimization in pytorch. arxiv e-prints, 1910 (2019)

[45] Maekawa, T., Kim, S., Nakai, D., Makino, C., Takagi, T., Ogura, H., Yamada, K., Chatton, B., Ishii, S.: Social isolation stress induces ATF-7 phosphorylation and impairs silencing of the 5-HT 5B receptor gene. The EMBO journal 29(1), 196–208 (2010)

[46] Chen, M., Liu, Y., Yang, Y., Qiu, Y., Wang, Z., Li, X., Zhang, W.: Emerging roles of activating transcription factor (ATF) family members in tumourigenesis and immunity: Implications in cancer immunotherapy. Genes & Diseases (2021)

[47] Gozdecka, M., Breitwieser, W.: The roles of ATF2 (activating transcription factor 2) in tumorigenesis. Biochemical Society Transactions 40(1), 230–234 (2012)

[48] Meijer, B.J., Giugliano, F.P., Baan, B., van der Meer, J.H., Meisner, S., van Roest, M., Koelink, P.J., de Boer, R.J., Jones, N., Breitwieser, W., et al.: ATF2 and ATF7 are critical mediators of intestinal epithelial repair. Cellular and molecular gastroenterology and hepatology 10(1), 23–42 (2020)

[49] Kim, S., Yu, N.-K., Kaang, B.-K.: CTCF as a multifunctional protein in genome regulation and gene expression. Experimental & molecular medicine 47(6), 166–166 (2015)

[50] Chen, H., Tian, Y., Shu, W., Bo, X., Wang, S.: Comprehensive identification and annotation of cell type-specific and ubiquitous CTCF-binding sites in the human genome. PloS one 7(7), 41374 (2012)

[51] Holwerda, S.J.B., de Laat, W.: CTCF: the protein, the binding partners, the binding sites and their chromatin loops. Philosophical Transactions of the Royal Society B: Biological Sciences 368(1620), 20120369 (2013)

[52] Vishnoi, K., Viswakarma, N., Rana, A., Rana, B.: Transcription factors in cancer development and therapy. Cancers 12(8), 2296 (2020)

[53] Dang, C.V.: MYC on the path to cancer. Cell 149(1), 22–35 (2012)

[54] Xu, J., Chen, Y., Olopade, O.I.: MYC and breast cancer. Genes & cancer 1(6), 629–640 (2010)

[55] Stine, Z.E., Walton, Z.E., Altman, B.J., Hsieh, A.L., Dang, C.V.: MYC, metabolism, and cancer. Cancer discovery 5(10), 1024–1039 (2015)

[56] Davudian, S., Mansoori, B., Shajari, N., Mohammadi, A., Baradaran, B.: BACH1, the master regulator gene: A novel candidate target for cancer therapy. Gene 588(1), 30–37 (2016)

[57] Zhang, X., Guo, J., Wei, X., Niu, C., Jia, M., Li, Q., Meng, D.: BACH1: function, regulation, and involvement in disease. Oxidative medicine and cellular longevity 2018 (2018)

[58] Padilla, J., Lee, J.: A novel therapeutic target, BACH1, regulates cancer metabolism. Cells 10(3), 634 (2021)

[59] Guo, X., Yang, M., Gu, H., Zhao, J., Zou, L.: Decreased expression of SOX6 confers a poor prognosis in hepatocellular carcinoma. Cancer epidemiology 37(5), 732–736 (2013)

[60] Jiang, W., Yuan, Q., Jiang, Y., Huang, L., Chen, C., Hu, G., Wan, R., Wang, X., Yang, L.: Identification of SOX6 as a regulator of pancreatic cancer development. Journal of Cellular and Molecular Medicine 22(3), 1864–1872 (2018)

[61] Wysocka, J., Reilly, P.T., Herr, W.: Loss of HCF-1–chromatin association precedes temperature-induced growth arrest of tsBN67 cells. Molecular and cellular biology 21(11), 3820–3829 (2001)

[62] Julien, E., Herr, W.: Proteolytic processing is necessary to separate and ensure proper cell growth and cytokinesis functions of HCF-1. The EMBO journal 22(10), 2360–2369 (2003)

[63] Khurana, B., Kristie, T.M.: A protein sequestering system reveals control of cellular programs by the transcriptional coactivator HCF-1. Journal of Biological Chemistry 279(32), 33673–33683 (2004)

[64] Maslova, A., Ramirez, R.N., Ma, K., Schmutz, H., Wang, C., Fox, C., Ng, B., Benoist, C., Mostafavi, S., et al.: Deep learning of immune cell differentiation. Proceedings of the National Academy of Sciences 117(41), 25655–25666 (2020)

[65] Gupta, S., Stamatoyannopoulos, J.A., Bailey, T.L., Noble, W.S.: Quantifying similarity between motifs. Genome biology 8(2), 1–9 (2007)

[66] De Graeve, F., Bahr, A., Sabapathy, K.T., Hauss, C., Wagner, E.F., Kedinger, C., Chatton, B.: Role of the ATFa/JNK2 complex in jun activation. Oncogene 18(23), 3491–3500 (1999)

[67] Grant, C.E., Bailey, T.L., Noble, W.S.: FIMO: scanning for occurrences of a given motif. Bioinformatics 27(7), 1017–1018 (2011)

[68] Bergmaier, P., Weth, O., Dienstbach, S., Boettger, T., Galjart, N., Mernberger, M., Bartkuhn, M., Renkawitz, R.: Choice of binding sites for CTCFL compared to CTCF is driven by chromatin and by sequence preference. Nucleic acids research 46(14), 7097–7107 (2018)

[69] Debaugny, R.E., Skok, J.A.: CTCF and CTCFL in cancer. Current opinion in genetics & development 61, 44–52 (2020)

[70] Ambrosini, G., Vorontsov, I., Penzar, D., Groux, R., Fornes, O., Nikolaeva, D.D., Ballester, B., Grau, J., Grosse, I., Makeev, V., et al.: Insights gained from a comprehensive all-against-all transcription factor binding motif benchmarking study. Genome biology 21(1), 1–18 (2020)

[71] Fornes, O., Castro-Mondragon, J.A., Khan, A., Van der Lee, R., Zhang, X., Richmond, P.A., Modi, B.P., Correard, S., Gheorghe, M., Baranašić, D., et al.: JASPAR 2020: update of the open-access database of transcription factor binding profiles. Nucleic acids research 48(D1), 87–92 (2020)

[72] Castro-Mondragon, J.A., Jaeger, S., Thieffry, D., Thomas-Chollier, M., Van Helden, J.: RSAT matrix-clustering: dynamic exploration and redundancy reduction of transcription factor binding motif collections. Nucleic Acids Research 45(13), 119–119 (2017)

[73] Blank, V.: Small MAF proteins in mammalian gene control: mere dimerization partners or dynamic transcriptional regulators? Journal of molecular biology 376(4), 913–925 (2008)

[74] Sizemore, G.M., Pitarresi, J.R., Balakrishnan, S., Ostrowski, M.C.: The ETS family of oncogenic transcription factors in solid tumours. Nature Reviews Cancer 17(6), 337–351 (2017)

[75] Hsu, T., Trojanowska, M., Watson, D.K.: ETS proteins in biological control and cancer. Journal of cellular biochemistry 91(5), 896–903 (2004)

[76] Wu, Z., Nicoll, M., Ingham, R.J.: AP-1 family transcription factors: a diverse family of proteins that regulate varied cellular activities in classical hodgkin lymphoma and alk+ alcl. Experimental Hematology & Oncology 10(1), 1–12 (2021)

[77] Trop-Steinberg, S., Azar, Y.: AP-1 expression and its clinical relevance in immune disorders and cancer. The American journal of the medical sciences 353(5), 474–483 (2017)

[78] Rohini, M., Menon, A.H., Selvamurugan, N.: Role of activating transcription factor 3 and its interacting proteins under physiological and pathological conditions. International journal of biological macromolecules 120, 310–317 (2018)

[79] Jiang, X., Xie, H., Dou, Y., Yuan, J., Zeng, D., Xiao, S.: Expression and function of FRA1 protein in tumors. Molecular Biology Reports 47(1), 737–752 (2020)

[80] Reddy, S.P., Mossman, B.T.: Role and regulation of activator protein-1 in toxicant-induced responses of the lung. American Journal of Physiology-Lung Cellular and Molecular Physiology 283(6), 1161–1178 (2002)

[81] Blonska, M., Zhu, Y., Chuang, H.H., You, M.J., Kunkalla, K., Vega, F., Lin, X.: JUN-regulated genes promote interaction of diffuse large b-cell lymphoma with the microenvironment. Blood, The Journal of the American Society of Hematology 125(6), 981–991 (2015)

[82] Fan, F., Bashari, M., Morelli, E., Tonon, G., Malvestiti, S., Vallet, S., Jarahian, M., Seckinger, A., Hose, D., Bakiri, L., et al.: The AP-1 transcription factor JUNB is essential for multiple myeloma cell proliferation and drug resistance in the bone marrow microenvironment. Leukemia 31(7), 1570–1581 (2017)

[83] Szremska, A.P., Kenner, L., Weisz, E., Ott, R.G., Passegué, E., Artwohl, M., Freissmuth, M., Stoxreiter, R., Theussl, H.-C., Parzer, S.B., et al.: JUNB inhibits proliferation and transformation in b-lymphoid cells. Blood 102(12), 4159–4165 (2003)

[84] Ott, R.G., Simma, O., Kollmann, K., Weisz, E., Zebedin, E.-M., Schorpp-Kistner, M., Heller, G., Zöchbauer, S., Wagner, E.F., Freissmuth, M., et al.: JUNB is a gatekeeper for b-lymphoid leukemia. Oncogene 26(33), 4863–4871 (2007)

[85] Passegué, E., Jochum, W., Schorpp-Kistner, M., Möhle-Steinlein, U., Wagner, E.F.: Chronic myeloid leukemia with increased granulocyte progenitors in mice lacking JUNB expression in the myeloid lineage. Cell 104(1), 21–32 (2001)

[86] Pechenick, D.A., Payne, J.L., Moore, J.H.: Phenotypic robustness and the assortativity signature of human transcription factor networks. PLoS computational biology 10(8), 1003780 (2014)

[87] Ren, B., Robert, F., Wyrick, J.J., Aparicio, O., Jennings, E.G., Simon, I., Zeitlinger, J., Schreiber, J., Hannett, N., Kanin, E., et al.: Genome-wide location and function of DNA binding proteins. science 290(5500), 2306–2309 (2000)

[88] Rhee, H.S., Pugh, B.F.: Comprehensive genome-wide protein-DNA interactions detected at single-nucleotide resolution. Cell 147(6), 1408–1419 (2011)

