## Supplementary material for "Cell Type Specific DNA Signatures of Transcription Factor Binding"

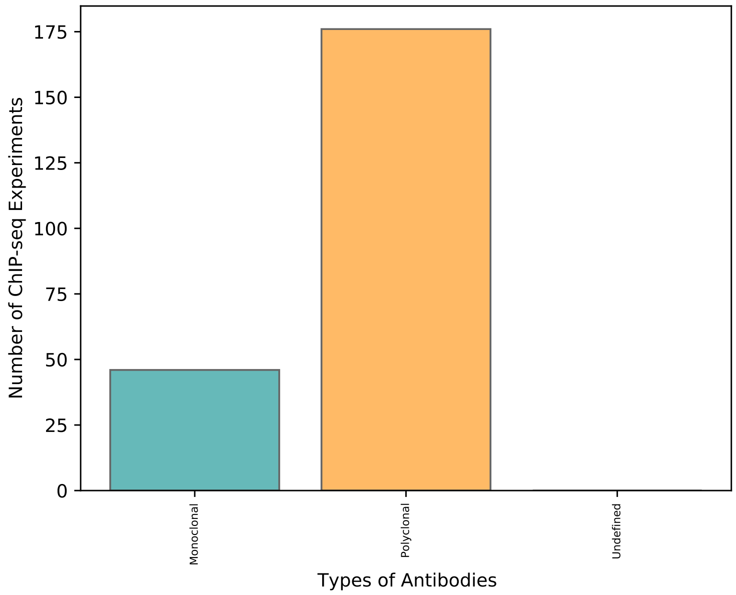


Figure 1: Number of ChIP-seq experiments using polyclonal vs monoclonal antibodies.

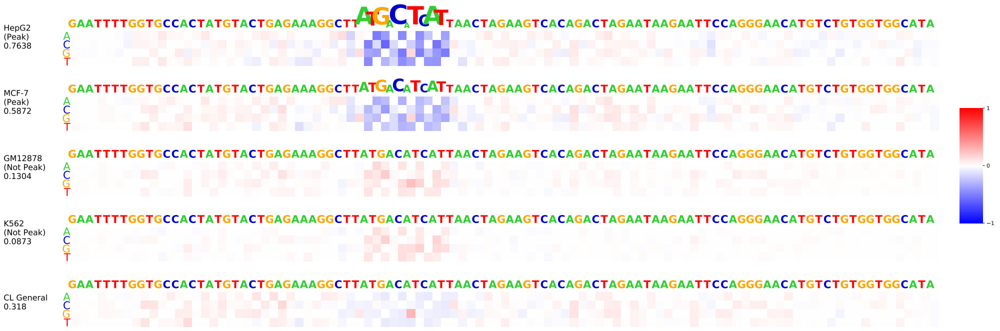


(a)

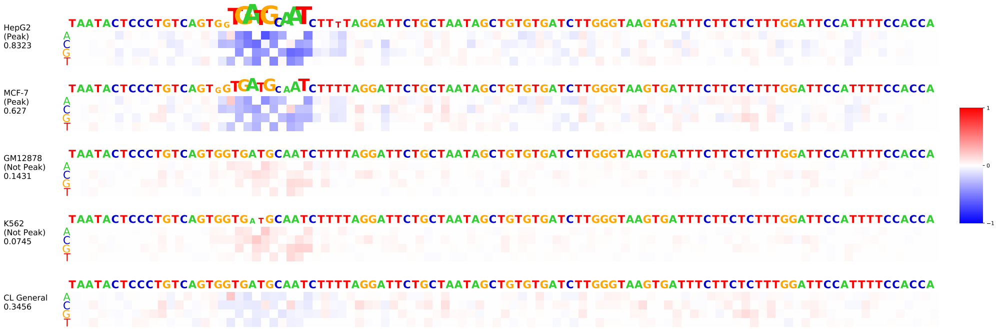


(b)

Figure 2: Mutation maps generated using in-silico mutagenesis on two different ATF7 test sequences. The instances are peaks, and considered to be peaks, in the first two cell lines: HepG2 and MCF-7. They are not peaks, and not considered to be peaks, in the other two cell lines: K562 and GM12878. The addition of cell type general information does not assist in prediction.

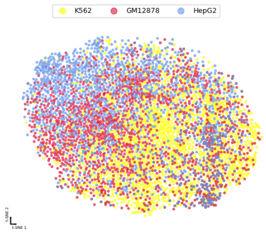


(a)

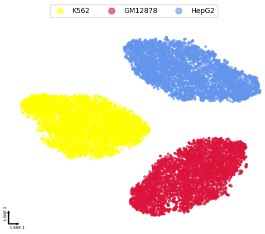


(b)

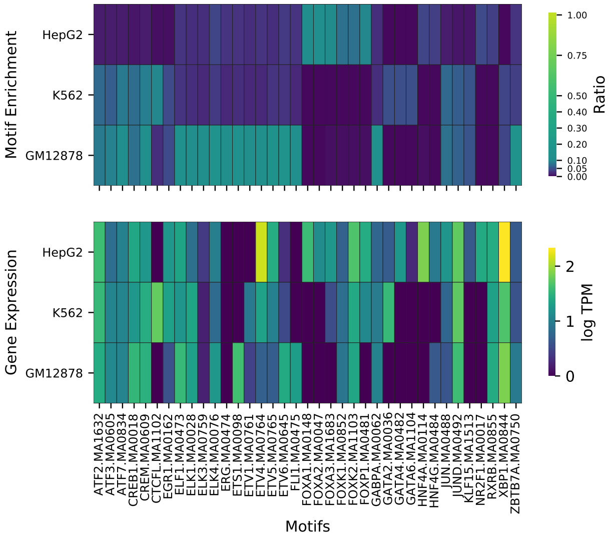


(c) CREM.ENCAB000AAT

Figure 3: Motif enrichment of CREM.ENCAB000AAT. t-SNE plots for both (a) CL General and (b) CL Specific cases. (c) Heatmaps of motif enrichment (top) and gene expression (bottom).

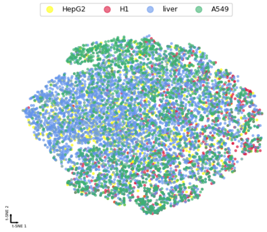


(a)

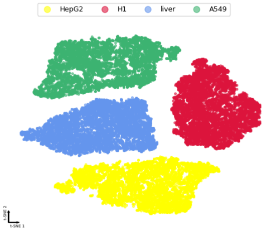


(b)

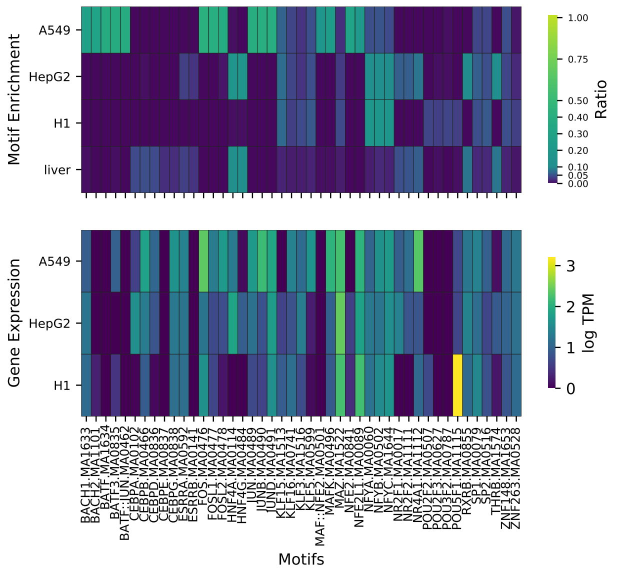


(c) SP1.ENCAB000AKY

Figure 4: Motif enrichment of SP1.ENCAB000AKY. t-SNE plots for both (a) CL General and (b) CL Specific cases. (c) Heatmaps of motif enrichment (top) and gene expression (bottom).

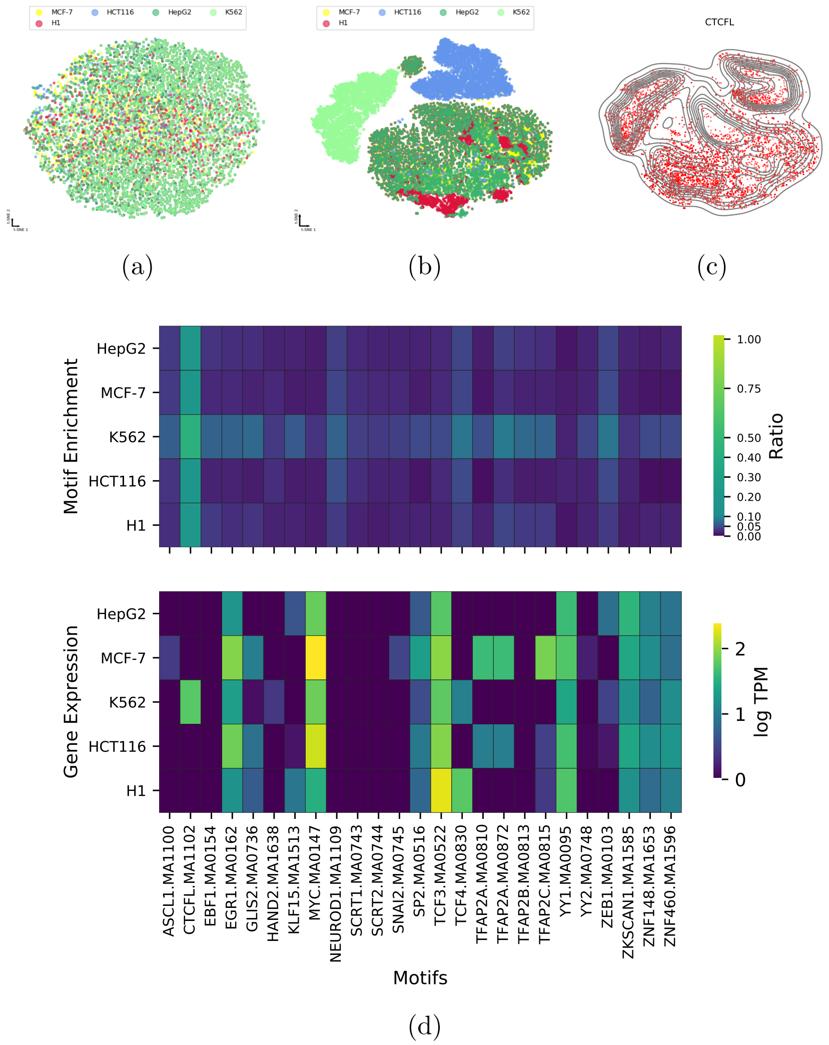


Figure 5: Motif enrichment of CTCF.ENCAB000AFR. t-SNE plots for both (a) CL General and (b) CL Specific cases. (c) Heatmaps of motif enrichment (top) and gene expression (bottom).

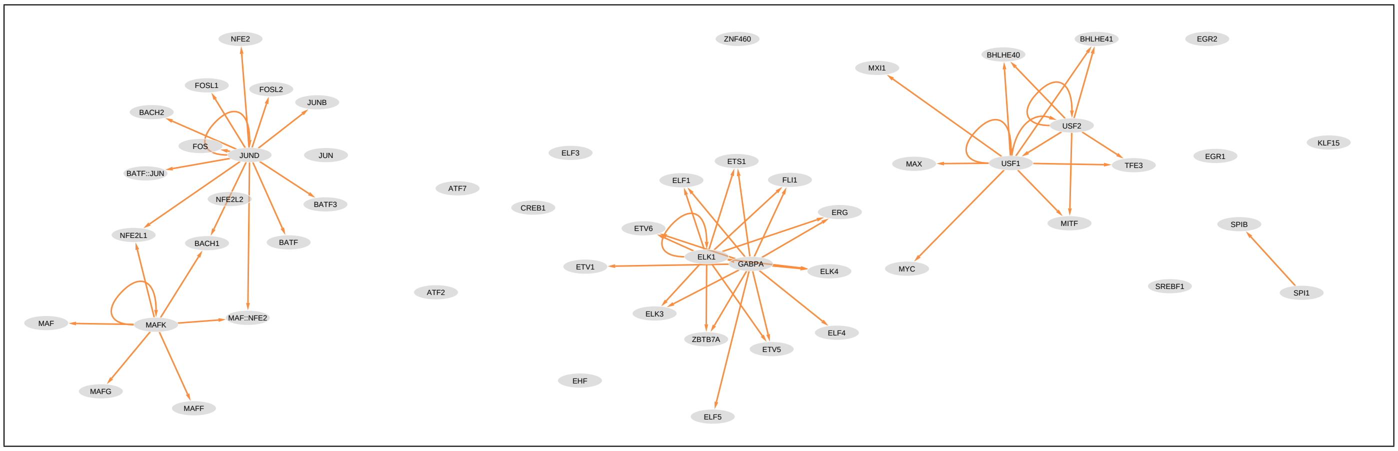


(a) GM12878

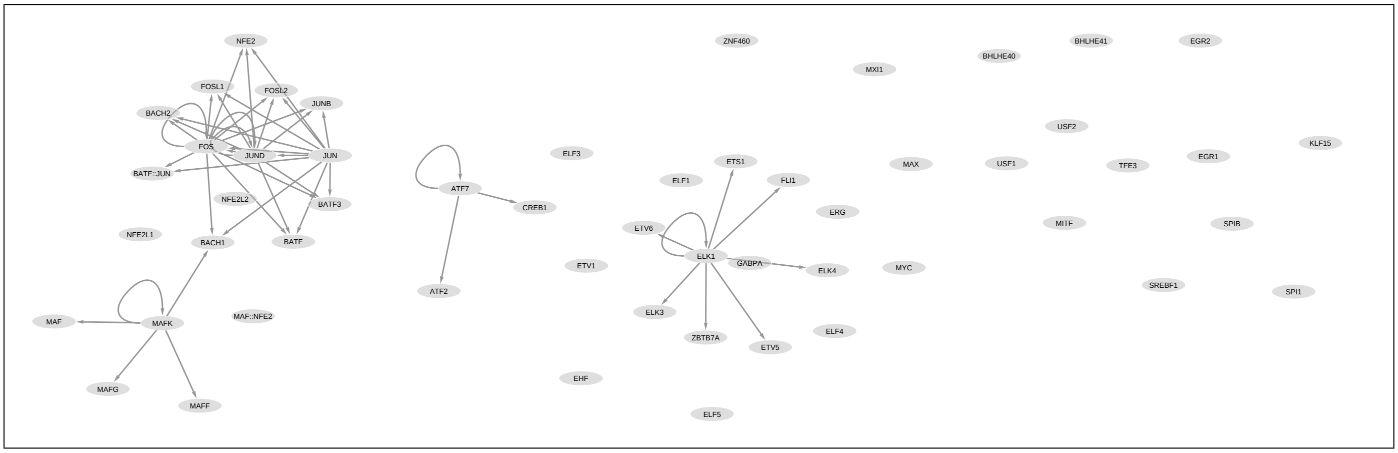


(b) MCF-7

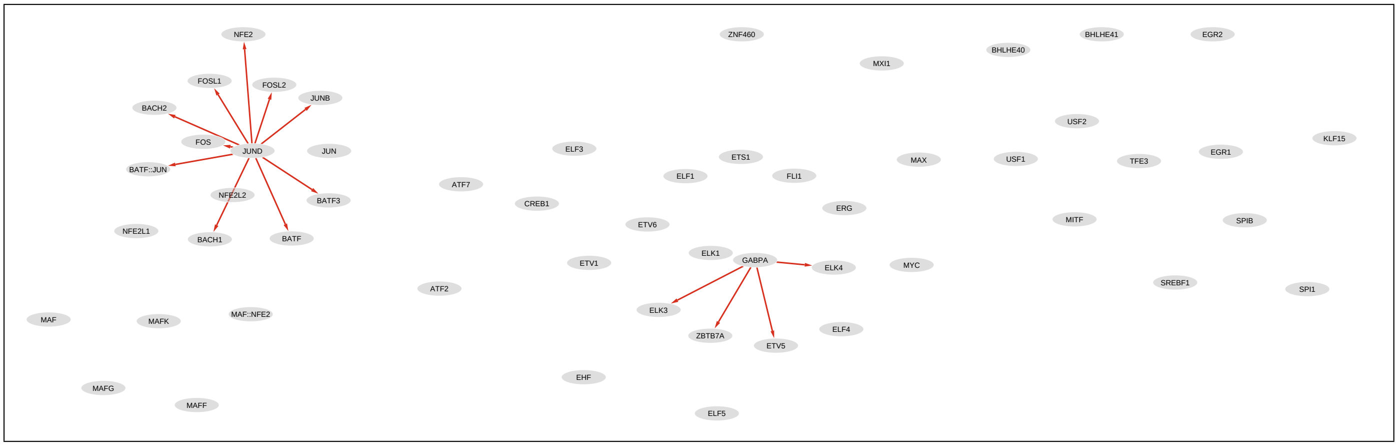


(c) Liver

Figure 6: Motif enrichment networks for each of cell types (a) GM12878, (b) MCF-7 and (c) Liver.

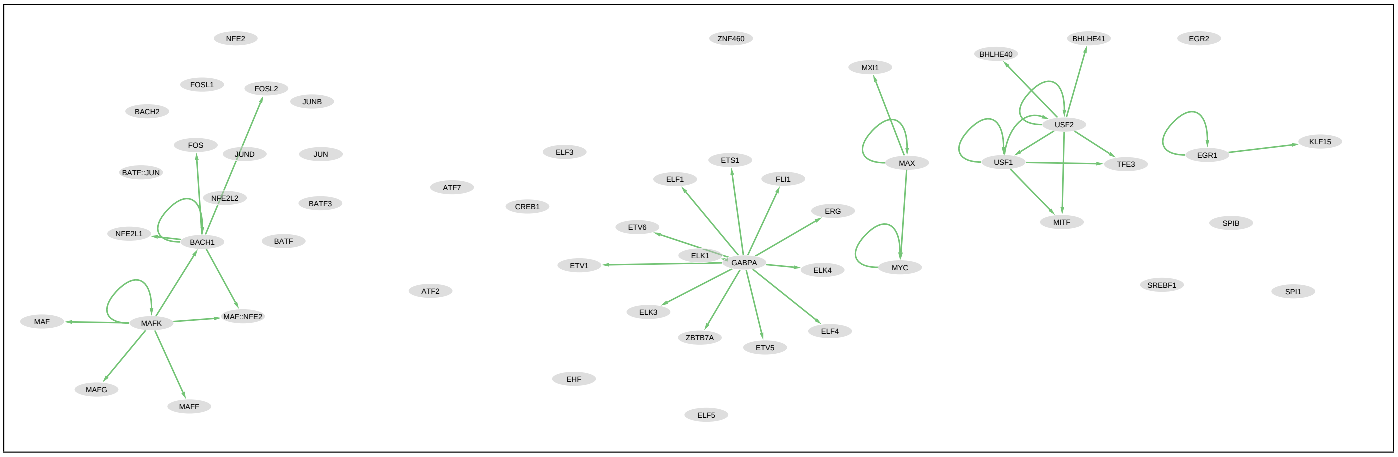


(d) H1

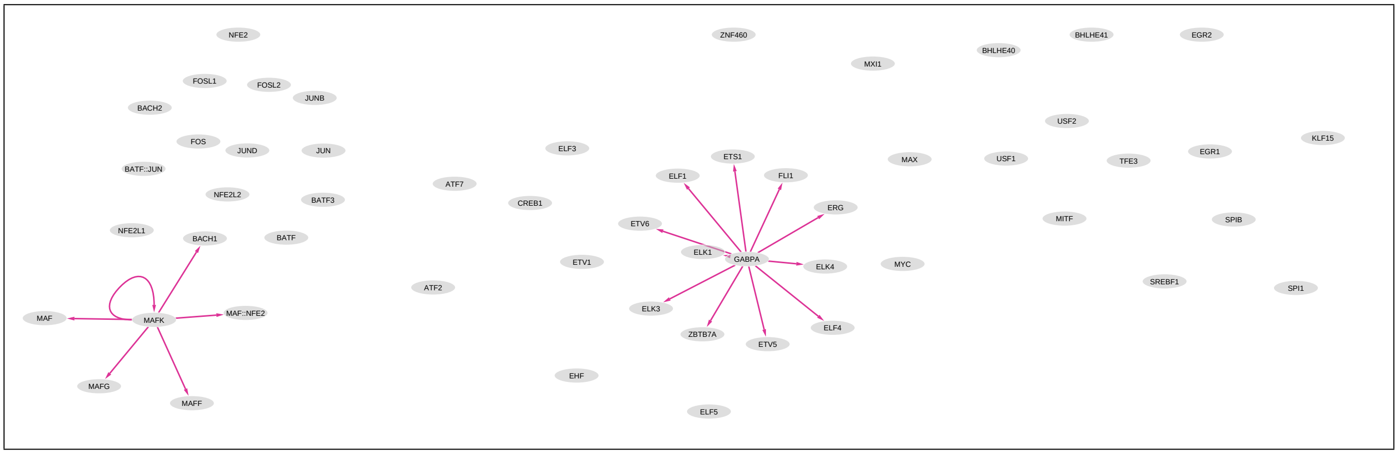


(e) HeLa-S3

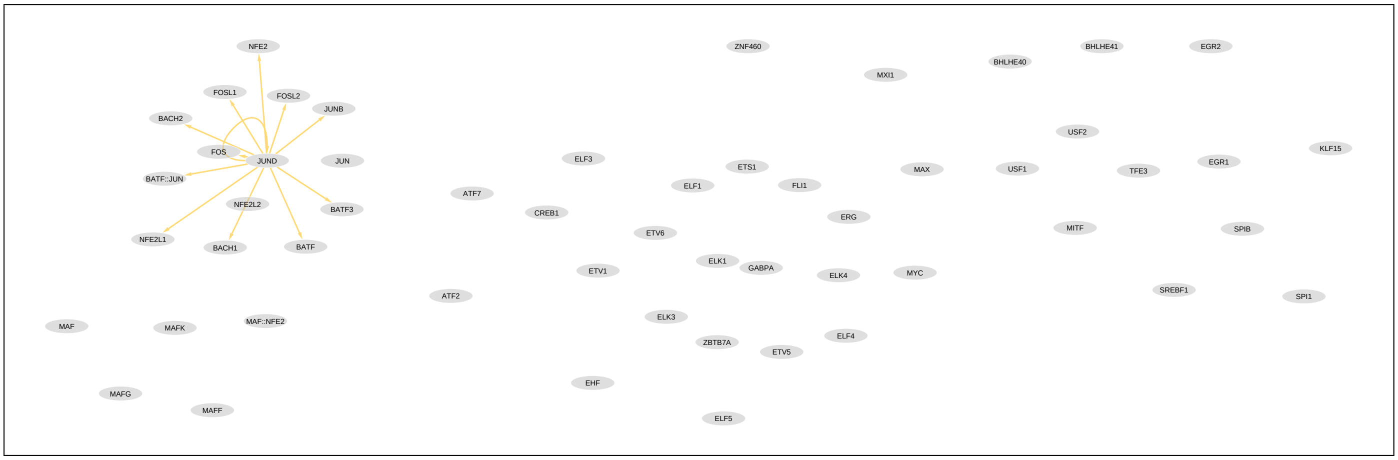


(f) HCT116

Figure 6: (continued) Motif enrichment networks for each of cell types (d) H1, (e) HeLa-S3 and (f) HCT116.

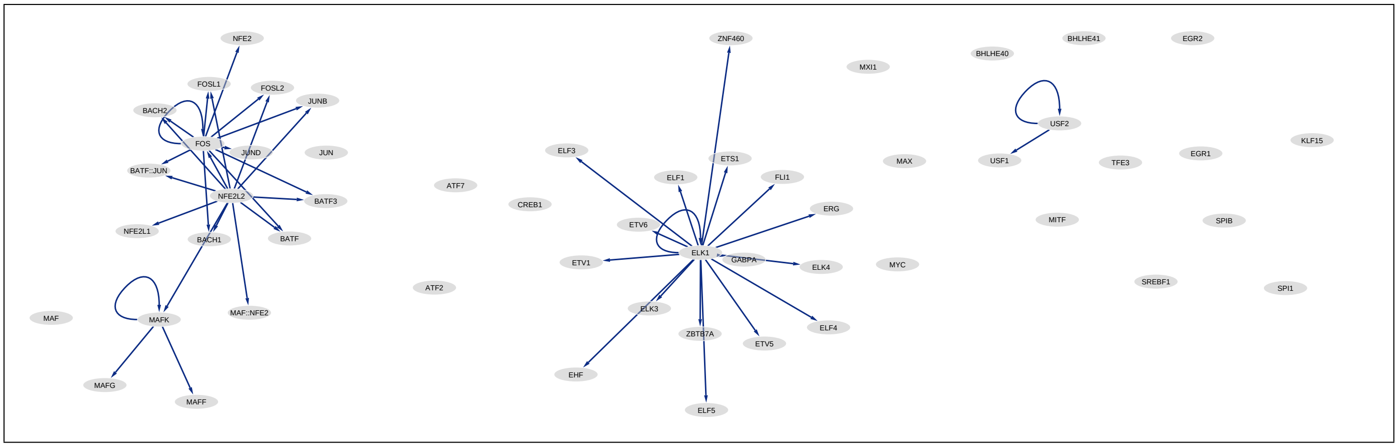


(g) IMR-90

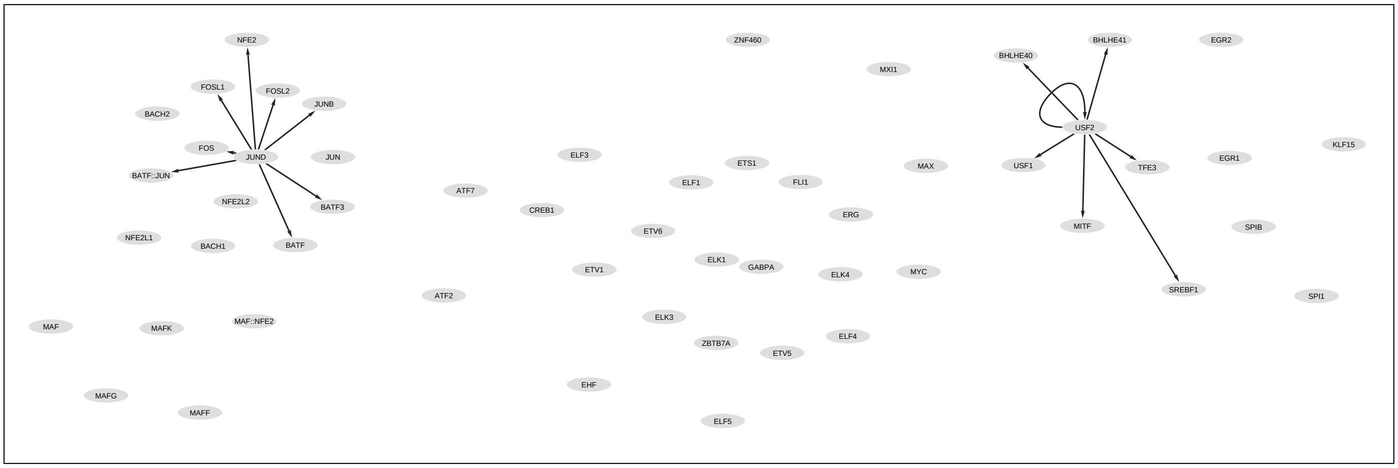


(h) SK-N-SH

Figure 6: (continued) Motif enrichment networks for each of cell types (g) IMR-90 and (h) SK-N-SH.

Table 1: TF-AB combinations and their corresponding AUC differences (score), cell types (CL ID) and actual ENCODE experiment used (ENCODE Experiments). Refer to SI Table 2 for the cell type names corresponding to the "CL ID".

| TF | AB | Score | CL ID | ENCODE Experiments |
|---|---|---|---|---|
| AGO1 | ENCAB189GDE | 0.06 | EFO:0002067<br>EFO:0001187 | ENCFF627BHP<br>ENCFF100VYA |
| ARNT | ENCAB608UKB | 0.06 | EFO:0002067<br>EFO:0002784 | ENCFF758RQJ<br>ENCFF655EFA |
| ASH2L | ENCAB947YBH | 0.01 | EFO:0001187<br>EFO:0002784 | ENCFF096XRG<br>ENCFF638IUM |
| ATF1 | ENCAB697XQW | 0.2 | EFO:0002067<br>EFO:0001187 | ENCFF914IWA<br>ENCFF715PLB |
| ATF2 | ENCAB605GOL | 0.14 | EFO:0002067<br>EFO:0001187<br>EFO:0002784 | ENCFF210HTZ<br>ENCFF803FHN<br>ENCFF089BQU |
| ATF3 | ENCAB000ADZ | 0.19 | EFO:0001086<br>EFO:0002067<br>EFO:0001187<br>UBERON:0002107<br>EFO:0003042 | ENCFF487GLV<br>ENCFF137OEY<br>ENCFF851UTY<br>ENCFF467WOR<br>ENCFF782SGI<br>ENCFF146URA |
| ATF7 | ENCAB000BMO | 0.21 | EFO:0002067<br>EFO:0001187<br>EFO:0002784<br>EFO:0001203 | ENCFF760ZVI<br>ENCFF371SJR<br>ENCFF495PWL<br>ENCFF498YGH |
| BACH1 | ENCAB000AEA | 0.17 | EFO:0002067<br>EFO:0002784<br>EFO:0003042 | ENCFF725YZH<br>ENCFF543FNN<br>ENCFF851YHG |
| BCL3 | ENCAB000AEG | 0.18 | EFO:0001086<br>EFO:0002784 | ENCFF093ZAB<br>ENCFF247MHT |
| BCLAF1 | ENCAB012FUX | 0.23 | EFO:0002067<br>EFO:0002784 | ENCFF381ZDU<br>ENCFF054DTJ |

| | | | | |
|---|---|---|---|---|
| BHLHE40 | ENCAB000AEK | 0.08 | EFO:0002067<br>EFO:0001187<br>EFO:0002784<br>EFO:0001196 | ENCFF567GON<br>ENCFF863ATX<br>ENCFF477JTV<br>ENCFF370ZNL<br>ENCFF622HGF |
| BMI1 | ENCAB000BDV | 0.14 | EFO:0002067<br>EFO:0002784<br>EFO:0001203 | ENCFF352DRR<br>ENCFF414LXZ<br>ENCFF592LPO |
| BRD4 | ENCAB782ZNQ | 0.01 | EFO:0002067<br>EFO:0001187 | ENCFF806CQB<br>ENCFF736GHL |
| CBX5 | ENCAB150NID | 0.0 | EFO:0002067<br>EFO:0002784 | ENCFF403TAE<br>ENCFF417SVR |
| CBX8 | ENCAB000AEX | 0.08 | EFO:0001086<br>EFO:0002067 | ENCFF330OCU<br>ENCFF210GJE |
| CEBPB | ENCAB000AFB | 0.11 | EFO:0001086<br>EFO:0002067<br>EFO:0001187<br>EFO:0001196<br>EFO:0002784 | ENCFF786YYI<br>ENCFF047UIF<br>ENCFF757KYL<br>ENCFF862DXR<br>ENCFF915ZYE<br>ENCFF813LOW<br>ENCFF321KQD |
| CEBPG | ENCAB728YTO | 0.11 | EFO:0002067<br>EFO:0001203 | ENCFF930PBH<br>ENCFF086CSF<br>ENCFF797MRW |
| CHD1 | ENCAB000AFE | 0.08 | EFO:0002784<br>EFO:0003042 | ENCFF863CTN<br>ENCFF806HXY<br>ENCFF549ODQ |
| CHD1 | ENCAB000AFF | 0.08 | EFO:0001196<br>EFO:0001203 | ENCFF510QXG<br>ENCFF730UAD |
| CHD2 | ENCAB000AFG | 0.1 | EFO:0001086<br>EFO:0001187<br>EFO:0003072<br>EFO:0002784<br>EFO:0003042 | ENCFF310IDS<br>ENCFF546AYN<br>ENCFF181XMM<br>ENCFF245UXM<br>ENCFF068MEO |
| CHD4 | ENCAB276UJU | 0.07 | EFO:0001086<br>EFO:0001187 | ENCFF766YPH<br>ENCFF148ABR |
| CREB1 | ENCAB279BHV | 0.06 | EFO:0001187<br>EFO:0001203 | ENCFF495PCJ<br>ENCFF550TXR |

| | | | | |
|---|---|---|---|---|
| CREM | ENCAB000AAT | 0.17 | EFO:0002067<br>EFO:0001187<br>EFO:0002784 | ENCFF091YID<br>ENCFF290UGF<br>ENCFF021XJN |
| CTBP1 | ENCAB000BAY | 0.23 | EFO:0002067<br>EFO:0001203 | ENCFF456MGR<br>ENCFF349UTF |
| CTCF | ENCAB000AFQ | 0.01 | EFO:0001086<br>EFO:0002067<br>EFO:0003072<br>EFO:0001196<br>EFO:0002784 | ENCFF396BZQ<br>ENCFF307XFM<br>ENCFF960ZGP<br>ENCFF540DWT<br>ENCFF646TUX |
| CTCF | ENCAB000AFR | 0.04 | EFO:0002824<br>EFO:0001203<br>EFO:0002067<br>EFO:0001187<br>EFO:0003042 | ENCFF549PGC<br>ENCFF543WTP<br>ENCFF821AQO<br>ENCFF119XFJ<br>ENCFF942TCG |
| CTCF | ENCAB000AXU | 0.01 | UBERON:0002107<br>CL:0002319<br>EFO:0003072 | ENCFF049UCF<br>ENCFF143HEE<br>ENCFF372JOV |
| CTCF | ENCAB000AXX | 0.01 | CL:0000103<br>NTR:0000711<br>CL:0000192<br>CL:0002551<br>CL:0002372<br>EFO:0002824<br>CL:0000062<br>EFO:0003072<br>CL:0000182<br>EFO:0006711<br>EFO:0002074<br>EFO:0007950<br>CL:0000127<br>EFO:0001203 | ENCFF203ZIS<br>ENCFF798RFA<br>ENCFF719TNH<br>ENCFF560GGY<br>ENCFF232FXZ<br>ENCFF186NOM<br>ENCFF476DVJ<br>ENCFF685KTA<br>ENCFF141MTA<br>ENCFF518MQA<br>ENCFF744PXO<br>ENCFF148BSH<br>ENCFF960XTR<br>ENCFF846FYU |

| | | | | |
|---|---|---|---|---|
| CTCF | ENCAB000AXY | 0.02 | EFO:0001086<br>CL:0000236<br>EFO:0002067<br>CL:0001054<br>EFO:0001187<br>CL:0000312<br>UBERON:0002106<br>UBERON:0001264<br>EFO:0002791<br>UBERON:0002048<br>CL:0002327<br>CL:0002553<br>EFO:0002784<br>EFO:0001203 | ENCFF505MGI<br>ENCFF068YLN<br>ENCFF139RCX<br>ENCFF502CZS<br>ENCFF861NDU<br>ENCFF954FAQ<br>ENCFF028IIR<br>ENCFF356LIU<br>ENCFF612ZUY<br>ENCFF519CXF<br>ENCFF300XXC<br>ENCFF910TER<br>ENCFF785NTC<br>ENCFF535MZG<br>ENCFF628EUU<br>ENCFF685HMV<br>ENCFF615GTV |
| CTCF | ENCAB247TYO | 0.11 | CL:0000103<br>EFO:0007950 | ENCFF322WKG<br>ENCFF904CNB |
| DDX20 | ENCAB398QAO | 0.16 | EFO:0002067<br>EFO:0001203 | ENCFF536LKB<br>ENCFF089GNH |
| DPF2 | ENCAB076EMA | 0.24 | EFO:0002067<br>EFO:0001203 | ENCFF217ZTP<br>ENCFF042AWM |
| DPF2 | ENCAB384ZPY | 0.25 | EFO:0002067<br>EFO:0002784 | ENCFF537VKZ<br>ENCFF771IAW |
| E2F4 | ENCAB000AFV | 0.2 | EFO:0002067<br>EFO:0002784 | ENCFF687SFB<br>ENCFF225TLP |
| E2F8 | ENCAB224FFQ | 0.1 | EFO:0002067<br>EFO:0002784<br>EFO:0001203 | ENCFF171WWF<br>ENCFF412GFI<br>ENCFF072VGV |

| | | | | |
|---|---|---|---|---|
| E4F1 | ENCAB944PGR | 0.23 | EFO:0002067<br>EFO:0002784<br>EFO:0001203 | ENCFF347USC<br>ENCFF752KNU<br>ENCFF035GFS |
| EGR1 | ENCAB000ASX | 0.0 | UBERON:0002107<br>EFO:0002067<br>EFO:0003042 | ENCFF477ANT<br>ENCFF561OGS<br>ENCFF617JQS<br>ENCFF808WST |
| EHMT2 | ENCAB282XQE | 0.13 | EFO:0001086<br>EFO:0002067<br>EFO:0001187 | ENCFF413RQL<br>ENCFF682XPD<br>ENCFF199OOU |
| ELF1 | ENCAB000AGA | 0.1 | EFO:0001086<br>EFO:0002067<br>EFO:0001187 | ENCFF935ZUW<br>ENCFF463GCH<br>ENCFF840RWO |
| ELF1 | ENCAB778OCV | 0.08 | EFO:0002067<br>EFO:0002784<br>EFO:0001203 | ENCFF617ZLL<br>ENCFF020UCD<br>ENCFF948CPI |
| ELK1 | ENCAB000AGB | 0.02 | EFO:0002067<br>EFO:0002784<br>EFO:0001203 | ENCFF119SCQ<br>ENCFF408TWV<br>ENCFF432AQP |
| EP300 | ENCAB000AJM | 0.08 | EFO:0002067<br>CL:0002319<br>EFO:0001187<br>EFO:0002791<br>EFO:0002784<br>EFO:0003042 | ENCFF755HCK<br>ENCFF924KFU<br>ENCFF459ARL<br>ENCFF674QCU<br>ENCFF834UVX<br>ENCFF865UDD<br>ENCFF510FUM |
| EP300 | ENCAB000AJO | 0.3 | EFO:0001187<br>EFO:0002784 | ENCFF080HJX<br>ENCFF806JJS |
| ESRRA | ENCAB000AGE | 0.15 | EFO:0001086<br>EFO:0002067<br>EFO:0002784<br>EFO:0001203 | ENCFF722LJP<br>ENCFF592GWM<br>ENCFF541DRZ<br>ENCFF558UWY |
| ETS1 | ENCAB000AGG | 0.06 | EFO:0007950<br>EFO:0001086<br>EFO:0002067<br>EFO:0002784 | ENCFF896WFR<br>ENCFF461PRP<br>ENCFF511AZU<br>ENCFF980VOD |

| | | | | |
|---|---|---|---|---|
| ETV6 | ENCAB000ABD | -0.0 | EFO:0002067<br>EFO:0002784 | ENCFF426GSY<br>ENCFF745ANU |
| ETV6 | ENCAB997CJG | 0.37 | EFO:0002067<br>EFO:0002784 | ENCFF658SGJ<br>ENCFF116AMK |
| EZH2 | ENCAB000AGH | 0.12 | CL:0002551<br>EFO:0001187<br>CL:0000515<br>EFO:0002791<br>CL:0002327<br>CL:0002553<br>EFO:0002784<br>CL:0000236 | ENCFF340LPI<br>ENCFF260KLJ<br>ENCFF504QZJ<br>ENCFF279MNV<br>ENCFF420KMT<br>ENCFF615NYO<br>ENCFF128OWK<br>ENCFF434OEY |
| EZH2 | ENCAB913HCF | 0.03 | CL:0000182<br>EFO:0005723 | ENCFF976SAN<br>ENCFF324UNA |
| EZH2 phospho T487 | ENCAB000BKV | 0.07 | NTR:0000711<br>EFO:0002074<br>CL:0000103<br>EFO:0005723 | ENCFF314ZKR<br>ENCFF320REA<br>ENCFF689GPW<br>ENCFF687VHB |
| FIP1L1 | ENCAB511DBP | 0.12 | EFO:0002067<br>EFO:0001187 | ENCFF084DTV<br>ENCFF031LBW |
| FOS | ENCAB000AEQ | 0.13 | CL:0002618<br>EFO:0001196<br>EFO:0001203 | ENCFF217ZMF<br>ENCFF327GZX<br>ENCFF170POB |
| FOSL2 | ENCAB000AGL | 0.16 | EFO:0001086<br>EFO:0001187 | ENCFF808RWZ<br>ENCFF054ESU |
| FOXA1 | ENCAB000AGM | 0.16 | EFO:0001086<br>EFO:0001187 | ENCFF152BOT<br>ENCFF297HAX<br>ENCFF167BKY |
| FOXA1 | ENCAB000AGN | 0.2 | EFO:0001187<br>UBERON:0002107 | ENCFF872MGU<br>ENCFF324QGE<br>ENCFF951VPZ |
| FOXA1 | ENCAB301QLE | 0.1 | EFO:0001187<br>EFO:0001203 | ENCFF160RLI<br>ENCFF367TQC |
| FOXA2 | ENCAB000AGO | 0.11 | EFO:0001187<br>UBERON:0002107 | ENCFF184NAC<br>ENCFF168JLI<br>ENCFF293LRQ |

| | | | | |
|---|---|---|---|---|
| FOXK2 | ENCAB625ERS | 0.17 | EFO:0002067<br>EFO:0001187<br>EFO:0002784<br>EFO:0001203 | ENCFF899MQW<br>ENCFF990MTR<br>ENCFF490EQR<br>ENCFF315CHX |
| FUS | ENCAB598VSJ | 0.03 | EFO:0002067<br>EFO:0001187 | ENCFF216YZI<br>ENCFF688ARM |
| GABPA | ENCAB000AGR | 0.06 | EFO:0001086<br>EFO:0002067<br>EFO:0001187<br>UBERON:0002107<br>EFO:0002791<br>EFO:0002784<br>EFO:0003042 | ENCFF520GJC<br>ENCFF124HAC<br>ENCFF225GFQ<br>ENCFF054HJA<br>ENCFF091UDB<br>ENCFF946ACA<br>ENCFF280YAF<br>ENCFF344XWK |
| GABPA | ENCAB728YTO | 0.05 | EFO:0002067<br>EFO:0001203 | ENCFF620TOF<br>ENCFF535BMV |
| GATA2 | ENCAB000AGT | 0.3 | EFO:0002067<br>CL:0002618 | ENCFF173TXA<br>ENCFF987YIJ |
| GATAD2B | ENCAB939ONI | 0.17 | EFO:0002067<br>EFO:0002784<br>EFO:0001203 | ENCFF046BRP<br>ENCFF298AIX<br>ENCFF569CMJ |
| GTF2F1 | ENCAB000AHE | 0.11 | EFO:0002067<br>EFO:0001187<br>EFO:0002791<br>EFO:0003042<br>EFO:0001203 | ENCFF876GXQ<br>ENCFF394JHN<br>ENCFF493PRB<br>ENCFF478HYJ<br>ENCFF343QQE |
| GTF2F1 | ENCAB108WPN | 0.33 | EFO:0002067<br>EFO:0001187 | ENCFF843UHP<br>ENCFF599TWF |
| HCFC1 | ENCAB000AHG | 0.02 | EFO:0002067<br>EFO:0001187<br>EFO:0002784<br>EFO:0001203 | ENCFF722QBB<br>ENCFF167RXK<br>ENCFF401IAI<br>ENCFF485SRU |
| HDAC1 | ENCAB649LYF | 0.16 | EFO:0002067<br>EFO:0001187 | ENCFF069KPS<br>ENCFF557WXK |

| | | | | |
|---|---|---|---|---|
| HDAC2 | ENCAB000AHI | 0.1 | EFO:0001086<br>EFO:0002067<br>EFO:0003042 | ENCFF363GSV<br>ENCFF497YNJ<br>ENCFF814DAF |
| HDAC2 | ENCAB000AHJ | 0.15 | EFO:0002067<br>EFO:0001187<br>EFO:0003042 | ENCFF589GSN<br>ENCFF009IVJ<br>ENCFF741IMY |
| HDGF | ENCAB173GGB | 0.17 | EFO:0002067<br>EFO:0002784<br>EFO:0001203 | ENCFF442WRJ<br>ENCFF161SFU<br>ENCFF575WFB |
| HES1 | ENCAB724OQW | 0.31 | EFO:0002067<br>EFO:0001203 | ENCFF010OOE<br>ENCFF144OPN |
| HNF4A | ENCAB000AHP | 0.15 | EFO:0001187<br>UBERON:0002107 | ENCFF072CXB<br>ENCFF837QHJ<br>ENCFF905JAC |
| HNF4G | ENCAB000AHQ | 0.09 | EFO:0001187<br>UBERON:0002107 | ENCFF497MUF<br>ENCFF086CTA |
| HNRNPK | ENCAB000BFN | 0.09 | EFO:0002067<br>EFO:0001187 | ENCFF828KXG<br>ENCFF984QUV |
| HNRNPL | ENCAB000AVZ | 0.23 | EFO:0002067<br>EFO:0001187 | ENCFF984ESZ<br>ENCFF039CUI |
| HNRNPLL | ENCAB698BTS | 0.12 | EFO:0002067<br>EFO:0001187 | ENCFF662WPN<br>ENCFF890KTX |
| IKZF1 | ENCAB144WPX | 0.32 | EFO:0002067<br>EFO:0001187<br>EFO:0002784 | ENCFF969BZA<br>ENCFF968NOG<br>ENCFF994OQH |
| IKZF1 | ENCAB590IRI | 0.27 | EFO:0002067<br>EFO:0002784 | ENCFF018NNF<br>ENCFF785BTP |
| JUN | ENCAB000AER | 0.12 | EFO:0002067<br>EFO:0003042<br>EFO:0001187<br>EFO:0001203 | ENCFF907UNK<br>ENCFF331IUK<br>ENCFF312GEN<br>ENCFF394CEC<br>ENCFF167WUZ<br>ENCFF881AVX<br>ENCFF672LKE<br>ENCFF032UMW |

| | | | | |
|---|---|---|---|---|
| JUNB | ENCAB000BQG | 0.0 | EFO:0002067<br>EFO:0002784 | ENCFF478XNA<br>ENCFF739XTO |
| JUND | ENCAB000AID | 0.02 | EFO:0001086<br>EFO:0002824<br>EFO:0002067<br>EFO:0001187<br>UBERON:0002107<br>EFO:0003072<br>EFO:0003042<br>EFO:0002784<br>EFO:0001203 | ENCFF587VEY<br>ENCFF569ZCY<br>ENCFF213EYD<br>ENCFF873DJD<br>ENCFF998KDQ<br>ENCFF246HKM<br>ENCFF539GRW<br>ENCFF420PED<br>ENCFF187QQB<br>ENCFF430PEI<br>ENCFF443HNU<br>ENCFF646IUA<br>ENCFF229COM |
| KDM1A | ENCAB000AIH | 0.11 | EFO:0001086<br>EFO:0001187<br>EFO:0003042 | ENCFF768FGG<br>ENCFF316CBQ<br>ENCFF562OAN |
| KDM5A | ENCAB914RSH | 0.06 | EFO:0001086<br>EFO:0001187 | ENCFF149INM<br>ENCFF334HKG |
| LARP7 | ENCAB568DRS | 0.32 | EFO:0002067<br>EFO:0002784 | ENCFF305SLO<br>ENCFF668ZTJ |
| MAFF | ENCAB717EYF | 0.14 | EFO:0002067<br>EFO:0002791<br>EFO:0001187 | ENCFF498MGH<br>ENCFF493TIR<br>ENCFF672LKL |
| MAFK | ENCAB000AIJ | 0.11 | EFO:0001086<br>EFO:0001203<br>EFO:0002067<br>EFO:0001187<br>EFO:0002791<br>EFO:0001196<br>EFO:0002784<br>EFO:0003042 | ENCFF813WJW<br>ENCFF171OJF<br>ENCFF873SVI<br>ENCFF351VGZ<br>ENCFF186AWV<br>ENCFF712RIS<br>ENCFF328IZQ<br>ENCFF893SCL |

| | | | | |
|---|---|---|---|---|
| MAX | ENCAB000AIL | 0.15 | EFO:0002067<br>EFO:0001187<br>UBERON:0002107<br>EFO:0002784<br>EFO:0003042 | ENCFF762LKG<br>ENCFF140PUO<br>ENCFF270NAL<br>ENCFF493ZMX<br>ENCFF618VMC<br>ENCFF669BQN<br>ENCFF900NVQ |
| MAZ | ENCAB000AIM | 0.05 | EFO:0001086<br>EFO:0001187<br>EFO:0001196<br>EFO:0002784<br>EFO:0001203 | ENCFF666YGQ<br>ENCFF144TBQ<br>ENCFF661NNJ<br>ENCFF916AJB<br>ENCFF348STZ |
| MBD2 | ENCAB000BQP | 0.13 | EFO:0002067<br>EFO:0001203 | ENCFF464QAL<br>ENCFF617QSK |
| MLLT1 | ENCAB650PBW | 0.16 | EFO:0002067<br>EFO:0002784<br>EFO:0001203 | ENCFF578NMN<br>ENCFF125MEN<br>ENCFF010AIG |
| MNT | ENCAB000BCL | 0.19 | EFO:0002067<br>EFO:0001187<br>EFO:0001203 | ENCFF562FMQ<br>ENCFF432GSK<br>ENCFF459DYU |
| MNT | ENCAB887GAG | 0.06 | EFO:0002067<br>EFO:0001187 | ENCFF454QQD<br>ENCFF482JSR |
| MTA1 | ENCAB000BCN | 0.24 | EFO:0002067<br>EFO:0001203 | ENCFF801KEW<br>ENCFF225VFR |
| MTA3 | ENCAB000BML | 0.24 | EFO:0002067<br>EFO:0001203 | ENCFF083AZM<br>ENCFF459XLR |
| MXI1 | ENCAB000AIT | 0.08 | EFO:0002067<br>CL:0002319<br>EFO:0002784<br>EFO:0003072 | ENCFF255WJM<br>ENCFF199HGX<br>ENCFF243QTL<br>ENCFF116RCK |

| | | | | |
|---|---|---|---|---|
| MYC | ENCAB000AET | 0.15 | EFO:0001086<br>EFO:0002067<br>EFO:0003042<br>EFO:0001203 | ENCFF392JJN<br>ENCFF542GMN<br>ENCFF339AQP<br>ENCFF605WXD<br>ENCFF370EQJ<br>ENCFF492XUU<br>ENCFF300OKR<br>ENCFF527EGF<br>ENCFF658XME<br>ENCFF700TLG |
| NANOG | ENCAB000AIX | 0.28 | EFO:0007950<br>EFO:0003042 | ENCFF621PFM<br>ENCFF794GVQ |
| NCOR1 | ENCAB805KAO | 0.26 | EFO:0002067<br>EFO:0001187 | ENCFF638IIC<br>ENCFF616RSZ |
| NEUROD1 | ENCAB000BDX | 0.28 | EFO:0002067<br>EFO:0001203 | ENCFF059LJD<br>ENCFF755APC |
| NFATC3 | ENCAB375FHW | 0.15 | EFO:0002067<br>EFO:0002784 | ENCFF430JFH<br>ENCFF704PDA |
| NFE2L2 | ENCAB800OND | 0.1 | EFO:0001086<br>EFO:0002791<br>EFO:0001187<br>EFO:0001196 | ENCFF882YLO<br>ENCFF305KIK<br>ENCFF474PPT<br>ENCFF418TUX |
| NFXL1 | ENCAB208FGX | 0.26 | EFO:0002067<br>EFO:0002784<br>EFO:0001203 | ENCFF329STX<br>ENCFF860IXB<br>ENCFF927DIO |
| NFYB | ENCAB000AJD | 0.11 | EFO:0002067<br>EFO:0002784 | ENCFF510NDO<br>ENCFF009NXK |
| NKRF | ENCAB893CIV | 0.14 | EFO:0002067<br>EFO:0002784 | ENCFF084NXU<br>ENCFF520QSR |
| NONO | ENCAB097MPS | 0.11 | EFO:0002067<br>EFO:0001187 | ENCFF823CQK<br>ENCFF420QKI |
| NONO | ENCAB349QGP | 0.34 | EFO:0002067<br>EFO:0001203 | ENCFF515YFU<br>ENCFF800CDQ |
| NR2C1 | ENCAB324ARN | 0.18 | EFO:0002067<br>EFO:0002784 | ENCFF462AKP<br>ENCFF023XHV |
| NR2F1 | ENCAB000ATZ | 0.2 | EFO:0002067<br>EFO:0002784 | ENCFF363IQN<br>ENCFF531KOV |

| | | | | |
|---|---|---|---|---|
| NR2F2 | ENCAB000AJH | 0.24 | EFO:0002067<br>UBERON:0002107 | ENCFF118HUH<br>ENCFF819WNB<br>ENCFF379TVQ |
| NR2F6 | ENCAB854ATP | 0.19 | EFO:0002067<br>EFO:0001187 | ENCFF350CKI<br>ENCFF194VBK |
| NRF1 | ENCAB000AJI | 0.05 | EFO:0001203<br>EFO:0001187<br>EFO:0002784<br>EFO:0003042 | ENCFF652BRY<br>ENCFF407IVS<br>ENCFF418DKQ<br>ENCFF269RME |
| NRF1 | ENCAB000BLM | 0.01 | EFO:0002067<br>EFO:0001187 | ENCFF313RFR<br>ENCFF626VDA<br>ENCFF543STN |
| PAX5 | ENCAB000AJS | 0.18 | EFO:0002785<br>EFO:0002786<br>EFO:0002784 | ENCFF997VAB<br>ENCFF196JGP<br>ENCFF987CQF |
| PAX8 | ENCAB000BOS | 0.34 | EFO:0002784<br>EFO:0001203 | ENCFF473UHQ<br>ENCFF992JWY |
| PBX2 | ENCAB697XQW | 0.11 | EFO:0002067<br>EFO:0001187 | ENCFF925OBR<br>ENCFF709YWO |
| PCBP1 | ENCAB000BFX | 0.0 | EFO:0002067<br>EFO:0001187 | ENCFF467RYH<br>ENCFF487WAN |
| PCBP2 | ENCAB000BFY | 0.33 | EFO:0002067<br>EFO:0001187 | ENCFF642XRH<br>ENCFF941XZW |
| PHF8 | ENCAB757LUG | 0.06 | EFO:0001086<br>EFO:0001187 | ENCFF907WHF<br>ENCFF202WIO |

| | | | | |
|---|---|---|---|---|
| POLR2A | ENCAB000AOC | 0.02 | EFO:0001086<br>EFO:0002824<br>EFO:0001203<br>EFO:0002067<br>EFO:0001187<br>EFO:0002786<br>EFO:0002791<br>CL:0002618<br>EFO:0002785<br>EFO:0002784<br>EFO:0003042 | ENCFF246QVY<br>ENCFF271RGE<br>ENCFF403ZEO<br>ENCFF422HDN<br>ENCFF964EVA<br>ENCFF565SUC<br>ENCFF387VGY<br>ENCFF455ZLJ<br>ENCFF021HUZ<br>ENCFF099NYA<br>ENCFF741JES<br>ENCFF798PUX<br>ENCFF730DLS<br>ENCFF664KTN<br>ENCFF668VIK<br>ENCFF915LKZ<br>ENCFF182YZG |
| POLR2AphosphoS2 | ENCAB000AOB | 0.02 | EFO:0001086<br>EFO:0002067<br>EFO:0001187<br>EFO:0002784 | ENCFF652YVO<br>ENCFF156MIR<br>ENCFF266OPF<br>ENCFF847DXY |
| POLR2G | ENCAB392AZR | 0.19 | EFO:0002067<br>EFO:0001187 | ENCFF283CUY<br>ENCFF551IJP |
| PRPF4 | ENCAB624FBH | 0.4 | EFO:0002067<br>EFO:0001187 | ENCFF417RQZ<br>ENCFF908QCS |
| PTBP1 | ENCAB553IRN | 0.25 | EFO:0002067<br>EFO:0001187 | ENCFF875ZPV<br>ENCFF917HXV |

| | | | | |
|---|---|---|---|---|
| RAD21 | ENCAB000AKG | 0.04 | EFO:0001086<br>CL:0002319<br>EFO:0001187<br>UBERON:0002107<br>EFO:0003072<br>EFO:0001196<br>EFO:0002784<br>EFO:0003042 | ENCFF557OCR<br>ENCFF897QCA<br>ENCFF654EGO<br>ENCFF454TRL<br>ENCFF895JAW<br>ENCFF295GOD<br>ENCFF255FRL<br>ENCFF874VFZ<br>ENCFF060IVS<br>ENCFF229WFR<br>ENCFF093XOJ<br>ENCFF315BSV |
| RAD21 | ENCAB697XQW | 0.09 | EFO:0001187<br>EFO:0001203 | ENCFF330VPL<br>ENCFF081TVG |
| RAD51 | ENCAB406ZKT | 0.13 | EFO:0002067<br>EFO:0001187<br>EFO:0002784<br>EFO:0001203 | ENCFF859MBC<br>ENCFF996NBR<br>ENCFF091AYX<br>ENCFF740OPF |
| RB1 | ENCAB000BFB | 0.02 | EFO:0002067<br>EFO:0002784 | ENCFF034OSV<br>ENCFF328QZM |
| RBBP5 | ENCAB000AKH | 0.03 | EFO:0002067<br>EFO:0003042 | ENCFF607WCG<br>ENCFF666PCE |
| RBFOX2 | ENCAB592TEY | 0.05 | EFO:0002067<br>EFO:0001187 | ENCFF871YRG<br>ENCFF232ASB |
| RBM22 | ENCAB476CFS | 0.32 | EFO:0002067<br>EFO:0001187 | ENCFF420IBN<br>ENCFF305WYD |
| RBM39 | ENCAB975CWB | 0.25 | EFO:0002067<br>EFO:0001187 | ENCFF420ALF<br>ENCFF503DIK |
| RCOR1 | ENCAB000AFK | 0.15 | EFO:0003072<br>EFO:0001187<br>EFO:0002784<br>EFO:0001196 | ENCFF073ADA<br>ENCFF470ZMK<br>ENCFF987VKU<br>ENCFF139EBY |

| | | | | |
|---|---|---|---|---|
| REST | ENCAB000AJK | 0.05 | EFO:0001086<br>EFO:0002067<br>EFO:0001187<br>EFO:0003072<br>UBERON:0002107<br>EFO:0002791<br>EFO:0002784<br>EFO:0003042 | ENCFF023ZUW<br>ENCFF208NUB<br>ENCFF107EWI<br>ENCFF403CAJ<br>ENCFF540FXB<br>ENCFF313CII<br>ENCFF669XCW<br>ENCFF986RRJ<br>ENCFF796YFZ<br>ENCFF274BBE<br>ENCFF288XHG<br>ENCFF178WRO |
| RFX5 | ENCAB000AKJ | 0.04 | EFO:0001086<br>EFO:0001203<br>EFO:0002067<br>EFO:0001187<br>EFO:0003072<br>EFO:0002784<br>EFO:0003042 | ENCFF502JJJ<br>ENCFF062WBN<br>ENCFF259LNG<br>ENCFF201YKU<br>ENCFF179WDI<br>ENCFF059GWW<br>ENCFF103MPW |
| RNF2 | ENCAB000BEA | 0.06 | EFO:0002067<br>EFO:0001187 | ENCFF380SYL<br>ENCFF820LKT |
| RNF2 | ENCAB790JUW | 0.26 | EFO:0002067<br>EFO:0003042 | ENCFF462AZY<br>ENCFF283MNG |
| RXRA | ENCAB000AKN | 0.2 | EFO:0003042<br>EFO:0001187<br>EFO:0002784<br>UBERON:0002107 | ENCFF313BDA<br>ENCFF105TFM<br>ENCFF430SIE<br>ENCFF572MCI<br>ENCFF201KGJ |
| SAP30 | ENCAB000AKO | 0.15 | EFO:0002067<br>EFO:0003042 | ENCFF103RHL<br>ENCFF193TFR |

| | | | | |
|---|---|---|---|---|
| SIN3A | ENCAB000AKR | 0.05 | EFO:0001086<br>EFO:0001203<br>EFO:0002067<br>EFO:0002784<br>EFO:0003042 | ENCFF514BGQ<br>ENCFF802JAN<br>ENCFF050CYK<br>ENCFF567BJI<br>ENCFF220RUS |
| SIN3A | ENCAB000AKS | 0.08 | EFO:0001086<br>EFO:0002067<br>EFO:0001187<br>EFO:0003072<br>EFO:0003042 | ENCFF663RUS<br>ENCFF407VGB<br>ENCFF635YMI<br>ENCFF708HTR<br>ENCFF905VZD |
| SIN3B | ENCAB194LUB | 0.19 | EFO:0002067<br>EFO:0001187 | ENCFF193DQZ<br>ENCFF543INR |
| SIX5 | ENCAB000AKV | 0.07 | EFO:0001086<br>EFO:0002067<br>EFO:0002784<br>EFO:0003042 | ENCFF864TFH<br>ENCFF189NMX<br>ENCFF644BNN<br>ENCFF247LOF |
| SKIL | ENCAB000BNW | 0.3 | EFO:0002067<br>EFO:0002784 | ENCFF254QDM<br>ENCFF903KEI |
| SMAD1 | ENCAB000AYH | 0.16 | EFO:0002067<br>EFO:0002784 | ENCFF987PGY<br>ENCFF084BUP |
| SMAD5 | ENCAB000AAG | 0.02 | EFO:0002067<br>EFO:0002784 | ENCFF855SJG<br>ENCFF069AAY |
| SMARCA4 | ENCAB000AEO | 0.26 | CL:0000103<br>EFO:0002067 | ENCFF703NAE<br>ENCFF482JUI |
| SMARCA5 | ENCAB528BVW | 0.21 | EFO:0002067<br>EFO:0002784<br>EFO:0001203 | ENCFF052STI<br>ENCFF481TNF<br>ENCFF618JNX |
| SMARCC2 | ENCAB313DWJ | 0.18 | EFO:0002067<br>EFO:0001187 | ENCFF150NHK<br>ENCFF751ZVX |
| SMARCE1 | ENCAB550CKA | 0.27 | EFO:0002067<br>EFO:0001203 | ENCFF435SZS<br>ENCFF761NKP |
| SMC3 | ENCAB000AKX | 0.02 | EFO:0001086<br>EFO:0002067<br>CL:0002319<br>EFO:0001187<br>EFO:0001196<br>EFO:0002784 | ENCFF035YWE<br>ENCFF572RPI<br>ENCFF944KJO<br>ENCFF380ZXB<br>ENCFF256LDD<br>ENCFF175UEE |

| | | | | |
|---|---|---|---|---|
| SNIP1 | ENCAB027LVC | 0.14 | EFO:0002067<br>EFO:0001203 | ENCFF529BDW<br>ENCFF455HWV |
| SOX6 | ENCAB000BOG | 0.43 | EFO:0002067<br>EFO:0001187 | ENCFF944LNI<br>ENCFF431STY |
| SP1 | ENCAB000AKY | 0.16 | UBERON:0002107<br>EFO:0001086<br>EFO:0001187<br>EFO:0003042 | ENCFF404OSB<br>ENCFF500JFI<br>ENCFF175VXL<br>ENCFF433EFF<br>ENCFF978TMH |
| SP1 | ENCAB000BAV | 0.25 | EFO:0002067<br>EFO:0001187<br>EFO:0001203 | ENCFF577EMC<br>ENCFF452LDK<br>ENCFF735WMX |
| SPI1 | ENCAB000AKF | 0.15 | EFO:0002067<br>EFO:0002785<br>EFO:0002784 | ENCFF414ECK<br>ENCFF744AGB<br>ENCFF071ZMW |
| SREBF1 | ENCAB000ALC | 0.14 | EFO:0001086<br>EFO:0002067<br>EFO:0001203 | ENCFF275WAD<br>ENCFF777MYW<br>ENCFF624DDK |
| SRF | ENCAB000ALE | 0.14 | EFO:0002784<br>EFO:0003042 | ENCFF345IDL<br>ENCFF182IFE<br>ENCFF829SEJ |
| STAT1 | ENCAB000ALF | 0.16 | EFO:0002067<br>EFO:0002784 | ENCFF323QQU<br>ENCFF747ICD<br>ENCFF431NLF<br>ENCFF646MXG |
| STAT5A | ENCAB000ALI | 0.29 | EFO:0002067<br>EFO:0002784 | ENCFF383YEA<br>ENCFF517IXK |
| SUZ12 | ENCAB000BEB | -0.02 | EFO:0002067<br>EFO:0001187 | ENCFF856HYC<br>ENCFF239LRW |

| | | | | |
|---|---|---|---|---|
| TAF1 | ENCAB000ALM | 0.02 | EFO:0001086<br>EFO:0001203<br>EFO:0002067<br>EFO:0001187<br>EFO:0002786<br>EFO:0003072<br>UBERON:0002107<br>EFO:0002791<br>EFO:0002785<br>EFO:0002784<br>EFO:0003042 | ENCFF033PLJ<br>ENCFF762MGC<br>ENCFF234TBW<br>ENCFF471NIK<br>ENCFF453TIB<br>ENCFF278XOE<br>ENCFF540AAP<br>ENCFF870SFJ<br>ENCFF886KDK<br>ENCFF214OJW<br>ENCFF423CTO |
| TARDBP | ENCAB000AUF | 0.02 | EFO:0002067<br>EFO:0002784 | ENCFF641AXD<br>ENCFF871LZM |
| TARDBP | ENCAB000BAX | 0.16 | EFO:0002067<br>EFO:0002784<br>EFO:0001203 | ENCFF909RMQ<br>ENCFF668JHK<br>ENCFF233RBO |
| TARDBP | ENCAB057RGG | 0.18 | EFO:0002067<br>EFO:0001187 | ENCFF448YOS<br>ENCFF696QPP |
| TBL1XR1 | ENCAB000ALP | 0.1 | EFO:0002067<br>EFO:0001187<br>EFO:0002784 | ENCFF126KGW<br>ENCFF392JWA<br>ENCFF239WFN |
| TBP | ENCAB000ALR | 0.08 | EFO:0002067<br>EFO:0001187<br>EFO:0002791<br>EFO:0002784<br>EFO:0003042 | ENCFF370YGS<br>ENCFF748YXF<br>ENCFF302RQH<br>ENCFF534GKQ<br>ENCFF896UZB |
| TCF12 | ENCAB000ALT | 0.22 | EFO:0001086<br>EFO:0001187<br>EFO:0002784<br>EFO:0003042 | ENCFF740HPV<br>ENCFF768VSH<br>ENCFF299JYV<br>ENCFF228CDD |
| TCF12 | ENCAB506UYG | 0.38 | EFO:0002067<br>EFO:0002784 | ENCFF897RYA<br>ENCFF912LXU |

| | | | | |
|---|---|---|---|---|
| TCF7 | ENCAB000ACR | 0.27 | EFO:0002067<br>EFO:0001187<br>EFO:0002784 | ENCFF152RNE<br>ENCFF512IAI<br>ENCFF928MIN |
| TOE1 | ENCAB755SML | 0.19 | EFO:0002067<br>EFO:0001187<br>EFO:0001203 | ENCFF539FVQ<br>ENCFF014WCO<br>ENCFF144VMM |
| TRIM22 | ENCAB000BNM | 0.33 | EFO:0001187<br>EFO:0002784<br>EFO:0001203 | ENCFF063GDN<br>ENCFF452VLA<br>ENCFF830TFU<br>ENCFF552WAH |
| U2AF1 | ENCAB298WVV | 0.33 | EFO:0002067<br>EFO:0001187 | ENCFF034KUO<br>ENCFF482DRO |
| USF1 | ENCAB000AMF | 0.08 | EFO:0002067<br>EFO:0001187<br>EFO:0002784<br>EFO:0003042 | ENCFF701QXK<br>ENCFF717KGR<br>ENCFF914IFQ<br>ENCFF699HXL |
| USF2 | ENCAB000AMH | 0.07 | EFO:0001086<br>EFO:0002067<br>EFO:0001196<br>EFO:0002784<br>EFO:0003042 | ENCFF514SWA<br>ENCFF425FVY<br>ENCFF593EOW<br>ENCFF938BOJ<br>ENCFF710JBU |
| XRCC5 | ENCAB308AOH | 0.04 | EFO:0002067<br>EFO:0001187 | ENCFF929TWP<br>ENCFF790ZAQ |
| YBX1 | ENCAB493UWX | 0.17 | EFO:0001187<br>EFO:0002784<br>EFO:0001203 | ENCFF332FUE<br>ENCFF247VVK<br>ENCFF500RBO |
| YY1 | ENCAB000ANS | 0.03 | EFO:0002067<br>EFO:0002786 | ENCFF072IHJ<br>ENCFF635XCI<br>ENCFF024TJO |

| | | | | |
|---|---|---|---|---|
| YY1 | ENCAB000ANT | 0.04 | EFO:0001086<br>EFO:0002824<br>EFO:0002067<br>EFO:0001187<br>EFO:0003072<br>UBERON:0002107<br>EFO:0002785<br>EFO:0002784<br>EFO:0003042 | ENCFF953BTB<br>ENCFF613DTQ<br>ENCFF363UWP<br>ENCFF094BQZ<br>ENCFF177YDT<br>ENCFF509GYP<br>ENCFF538VYU<br>ENCFF223MUF<br>ENCFF838VFX<br>ENCFF459TWF |
| ZBED1 | ENCAB000AAK | 0.21 | EFO:0002067<br>EFO:0002784 | ENCFF388TYU<br>ENCFF630FLK |
| ZBTB33 | ENCAB000AML | 0.15 | EFO:0001086<br>EFO:0002824<br>EFO:0001187<br>UBERON:0002107<br>EFO:0002784 | ENCFF422MCZ<br>ENCFF593ZJA<br>ENCFF943WRA<br>ENCFF773OQL<br>ENCFF727ZIT<br>ENCFF882UHR |
| ZBTB33 | ENCAB292USO | 0.21 | EFO:0002067<br>EFO:0002784<br>EFO:0001203 | ENCFF556STK<br>ENCFF780WLS<br>ENCFF475DID |
| ZBTB40 | ENCAB373DVF | 0.08 | EFO:0002067<br>EFO:0001187<br>EFO:0002784<br>EFO:0001203 | ENCFF932XEU<br>ENCFF624WDI<br>ENCFF088LZZ<br>ENCFF084IUW |
| ZBTB7A | ENCAB000AMM | 0.07 | EFO:0002067<br>EFO:0001187 | ENCFF953JQD<br>ENCFF245LRG |
| ZC3H11A | ENCAB000AMN | 0.44 | EFO:0001086<br>EFO:0002067 | ENCFF478PGJ<br>ENCFF415SIS |

| | | | | |
|---|---|---|---|---|
| ZFP36 | ENCAB118PND | 0.09 | EFO:0001086<br>EFO:0002067<br>EFO:0001187<br>EFO:0002791<br>EFO:0002784 | ENCFF166GKK<br>ENCFF429XQI<br>ENCFF137JHO<br>ENCFF224WII<br>ENCFF763HPQ |
| ZFX | ENCAB657HDP | 0.05 | EFO:0001203<br>EFO:0002824 | ENCFF215SIC<br>ENCFF775BWJ |
| ZHX1 | ENCAB361RPF | 0.04 | EFO:0002067<br>EFO:0002791 | ENCFF267DZF<br>ENCFF495BPY |
| ZHX2 | ENCAB000ATW | 0.17 | EFO:0001187<br>EFO:0001203 | ENCFF694ZRC<br>ENCFF964KDQ |
| ZKSCAN1 | ENCAB000AMP | 0.26 | EFO:0002067<br>EFO:0001187<br>EFO:0001203 | ENCFF687REM<br>ENCFF704VDI<br>ENCFF721NEC |
| ZMYM3 | ENCAB426WVA | 0.19 | EFO:0002067<br>EFO:0001187 | ENCFF195IFB<br>ENCFF769SEZ |
| ZNF143 | ENCAB000AMR | 0.06 | EFO:0002067<br>EFO:0002784<br>EFO:0003042 | ENCFF933WSP<br>ENCFF700GZI<br>ENCFF193POQ<br>ENCFF153TQR |
| ZNF207 | ENCAB000BNU | 0.24 | EFO:0002784<br>EFO:0001203 | ENCFF676BIG<br>ENCFF621ZSK |
| ZNF217 | ENCAB182PZR | 0.15 | EFO:0002784<br>EFO:0001203 | ENCFF200SLC<br>ENCFF620RPM |
| ZNF24 | ENCAB060JJI | 0.1 | EFO:0002067<br>EFO:0001187 | ENCFF858WPR<br>ENCFF723JDW |
| ZNF24 | ENCAB198YAJ | 0.08 | EFO:0002067<br>EFO:0001187<br>EFO:0002784<br>EFO:0001203 | ENCFF619BFO<br>ENCFF313HBL<br>ENCFF260CBQ<br>ENCFF904QAD |
| ZNF282 | ENCAB503NQV | 0.0 | EFO:0002067<br>EFO:0001187 | ENCFF596JDS<br>ENCFF482XNG |
| ZNF592 | ENCAB438BKV | 0.1 | EFO:0002067<br>EFO:0001203 | ENCFF972UGK<br>ENCFF541HRT |
| ZNF687 | ENCAB146FZU | 0.14 | EFO:0002784<br>EFO:0001203 | ENCFF137BRA<br>ENCFF329QYZ |
| ZSCAN29 | ENCAB211EDR | 0.1 | EFO:0002067<br>EFO:0002784 | ENCFF214NJL<br>ENCFF979GFF |

| | | | | |
|---|---|---|---|---|
| AGO1 | ENCAB189GDE | 0.06 | EFO:0002067<br>EFO:0001187 | ENCFF627BHP<br>ENCFF100VYA |
| ARNT | ENCAB608UKB | 0.06 | EFO:0002067<br>EFO:0002784 | ENCFF758RQJ<br>ENCFF655EFA |
| ASH2L | ENCAB947YBH | 0.01 | EFO:0001187<br>EFO:0002784 | ENCFF096XRG<br>ENCFF638IUM |
| ATF1 | ENCAB697XQW | 0.2 | EFO:0002067<br>EFO:0001187 | ENCFF914IWA<br>ENCFF715PLB |
| ATF2 | ENCAB605GOL | 0.14 | EFO:0002067<br>EFO:0001187<br>EFO:0002784 | ENCFF210HTZ<br>ENCFF803FHN<br>ENCFF089BQU |
| ATF3 | ENCAB000ADZ | 0.19 | EFO:0001086<br>EFO:0002067<br>EFO:0001187<br>UBERON:0002107<br>EFO:0003042 | ENCFF487GLV<br>ENCFF137OEY<br>ENCFF851UTY<br>ENCFF467WOR<br>ENCFF782SGI<br>ENCFF146URA |
| ATF7 | ENCAB000BMO | 0.21 | EFO:0002067<br>EFO:0001187<br>EFO:0002784<br>EFO:0001203 | ENCFF760ZVI<br>ENCFF371SJR<br>ENCFF495PWL<br>ENCFF498YGH |
| BACH1 | ENCAB000AEA | 0.17 | EFO:0002067<br>EFO:0002784<br>EFO:0003042 | ENCFF725YZH<br>ENCFF543FNN<br>ENCFF851YHG |
| BCL3 | ENCAB000AEG | 0.18 | EFO:0001086<br>EFO:0002784 | ENCFF093ZAB<br>ENCFF247MHT |
| BCLAF1 | ENCAB012FUX | 0.23 | EFO:0002067<br>EFO:0002784 | ENCFF381ZDU<br>ENCFF054DTJ |
| BHLHE40 | ENCAB000AEK | 0.08 | EFO:0002067<br>EFO:0001187<br>EFO:0002784<br>EFO:0001196 | ENCFF567GON<br>ENCFF863ATX<br>ENCFF477JTV<br>ENCFF370ZNL<br>ENCFF622HGF |
| BMI1 | ENCAB000BDV | 0.14 | EFO:0002067<br>EFO:0002784<br>EFO:0001203 | ENCFF352DRR<br>ENCFF414LXZ<br>ENCFF592LPO |

| | | | | |
|---|---|---|---|---|
| BRD4 | ENCAB782ZNQ | 0.01 | EFO:0002067<br>EFO:0001187 | ENCFF806CQB<br>ENCFF736GHL |
| CBX5 | ENCAB150NID | 0.0 | EFO:0002067<br>EFO:0002784 | ENCFF403TAE<br>ENCFF417SVR |
| CBX8 | ENCAB000AEX | 0.08 | EFO:0001086<br>EFO:0002067 | ENCFF330OCU<br>ENCFF210GJE |
| CEBPB | ENCAB000AFB | 0.11 | EFO:0001086<br>EFO:0002067<br>EFO:0001187<br>EFO:0001196<br>EFO:0002784 | ENCFF786YYI<br>ENCFF047UIF<br>ENCFF757KYL<br>ENCFF862DXR<br>ENCFF915ZYE<br>ENCFF813LOW<br>ENCFF321KQD |
| CEBPG | ENCAB728YTO | 0.11 | EFO:0002067<br>EFO:0001203 | ENCFF930PBH<br>ENCFF086CSF<br>ENCFF797MRW |
| CHD1 | ENCAB000AFE | 0.08 | EFO:0002784<br>EFO:0003042 | ENCFF863CTN<br>ENCFF806HXY<br>ENCFF549ODQ |
| CHD1 | ENCAB000AFF | 0.08 | EFO:0001196<br>EFO:0001203 | ENCFF510QXG<br>ENCFF730UAD |
| CHD2 | ENCAB000AFG | 0.1 | EFO:0001086<br>EFO:0001187<br>EFO:0003072<br>EFO:0002784<br>EFO:0003042 | ENCFF310IDS<br>ENCFF546AYN<br>ENCFF181XMM<br>ENCFF245UXM<br>ENCFF068MEO |
| CHD4 | ENCAB276UJU | 0.07 | EFO:0001086<br>EFO:0001187 | ENCFF766YPH<br>ENCFF148ABR |
| CREB1 | ENCAB279BHV | 0.06 | EFO:0001187<br>EFO:0001203 | ENCFF495PCJ<br>ENCFF550TXR |
| CREM | ENCAB000AAT | 0.17 | EFO:0002067<br>EFO:0001187<br>EFO:0002784 | ENCFF091YID<br>ENCFF290UGF<br>ENCFF021XJN |
| CTBP1 | ENCAB000BAY | 0.23 | EFO:0002067<br>EFO:0001203 | ENCFF456MGR<br>ENCFF349UTF |

| | | | | |
|---|---|---|---|---|
| CTCF | ENCAB000AFQ | 0.01 | EFO:0001086<br>EFO:0002067<br>EFO:0003072<br>EFO:0001196<br>EFO:0002784 | ENCFF396BZQ<br>ENCFF307XFM<br>ENCFF960ZGP<br>ENCFF540DWT<br>ENCFF646TUX |
| CTCF | ENCAB000AFR | 0.04 | EFO:0002824<br>EFO:0001203<br>EFO:0002067<br>EFO:0001187<br>EFO:0003042 | ENCFF549PGC<br>ENCFF543WTP<br>ENCFF821AQO<br>ENCFF119XFJ<br>ENCFF942TCG |
| CTCF | ENCAB000AXU | 0.01 | UBERON:0002107<br>CL:0002319<br>EFO:0003072 | ENCFF049UCF<br>ENCFF143HEE<br>ENCFF372JOV |
| CTCF | ENCAB000AXX | 0.01 | CL:0000103<br>NTR:0000711<br>CL:0000192<br>CL:0002551<br>CL:0002372<br>EFO:0002824<br>CL:0000062<br>EFO:0003072<br>CL:0000182<br>EFO:0006711<br>EFO:0002074<br>EFO:0007950<br>CL:0000127<br>EFO:0001203 | ENCFF203ZIS<br>ENCFF798RFA<br>ENCFF719TNH<br>ENCFF560GGY<br>ENCFF232FXZ<br>ENCFF186NOM<br>ENCFF476DVJ<br>ENCFF685KTA<br>ENCFF141MTA<br>ENCFF518MQA<br>ENCFF744PXO<br>ENCFF148BSH<br>ENCFF960XTR<br>ENCFF846FYU |

| | | | | |
|---|---|---|---|---|
| CTCF | ENCAB000AXY | 0.02 | EFO:0001086<br>CL:0000236<br>EFO:0002067<br>CL:0001054<br>EFO:0001187<br>CL:0000312<br>UBERON:0002106<br>UBERON:0001264<br>EFO:0002791<br>UBERON:0002048<br>CL:0002327<br>CL:0002553<br>EFO:0002784<br>EFO:0001203 | ENCFF505MGI<br>ENCFF068YLN<br>ENCFF139RCX<br>ENCFF502CZS<br>ENCFF861NDU<br>ENCFF954FAQ<br>ENCFF028IIR<br>ENCFF356LIU<br>ENCFF612ZUY<br>ENCFF519CXF<br>ENCFF300XXC<br>ENCFF910TER<br>ENCFF785NTC<br>ENCFF535MZG<br>ENCFF628EUU<br>ENCFF685HMV<br>ENCFF615GTV |
| CTCF | ENCAB247TYO | 0.11 | CL:0000103<br>EFO:0007950 | ENCFF322WKG<br>ENCFF904CNB |
| DDX20 | ENCAB398QAO | 0.16 | EFO:0002067<br>EFO:0001203 | ENCFF536LKB<br>ENCFF089GNH |
| DPF2 | ENCAB076EMA | 0.24 | EFO:0002067<br>EFO:0001203 | ENCFF217ZTP<br>ENCFF042AWM |
| DPF2 | ENCAB384ZPY | 0.25 | EFO:0002067<br>EFO:0002784 | ENCFF537VKZ<br>ENCFF771IAW |
| E2F4 | ENCAB000AFV | 0.2 | EFO:0002067<br>EFO:0002784 | ENCFF687SFB<br>ENCFF225TLP |
| E2F8 | ENCAB224FFQ | 0.1 | EFO:0002067<br>EFO:0002784<br>EFO:0001203 | ENCFF171WWF<br>ENCFF412GFI<br>ENCFF072VGV |

| | | | | |
|---|---|---|---|---|
| E4F1 | ENCAB944PGR | 0.23 | EFO:0002067<br>EFO:0002784<br>EFO:0001203 | ENCFF347USC<br>ENCFF752KNU<br>ENCFF035GFS |
| EGR1 | ENCAB000ASX | 0.0 | UBERON:0002107<br>EFO:0002067<br>EFO:0003042 | ENCFF477ANT<br>ENCFF561OGS<br>ENCFF617JQS<br>ENCFF808WST |
| EHMT2 | ENCAB282XQE | 0.13 | EFO:0001086<br>EFO:0002067<br>EFO:0001187 | ENCFF413RQL<br>ENCFF682XPD<br>ENCFF199OOU |
| ELF1 | ENCAB000AGA | 0.1 | EFO:0001086<br>EFO:0002067<br>EFO:0001187 | ENCFF935ZUW<br>ENCFF463GCH<br>ENCFF840RWO |
| ELF1 | ENCAB778OCV | 0.08 | EFO:0002067<br>EFO:0002784<br>EFO:0001203 | ENCFF617ZLL<br>ENCFF020UCD<br>ENCFF948CPI |
| ELK1 | ENCAB000AGB | 0.02 | EFO:0002067<br>EFO:0002784<br>EFO:0001203 | ENCFF119SCQ<br>ENCFF408TWV<br>ENCFF432AQP |
| EP300 | ENCAB000AJM | 0.08 | EFO:0002067<br>CL:0002319<br>EFO:0001187<br>EFO:0002791<br>EFO:0002784<br>EFO:0003042 | ENCFF755HCK<br>ENCFF924KFU<br>ENCFF459ARL<br>ENCFF674QCU<br>ENCFF834UVX<br>ENCFF865UDD<br>ENCFF510FUM |
| EP300 | ENCAB000AJO | 0.3 | EFO:0001187<br>EFO:0002784 | ENCFF080HJX<br>ENCFF806JJS |
| ESRRA | ENCAB000AGE | 0.15 | EFO:0001086<br>EFO:0002067<br>EFO:0002784<br>EFO:0001203 | ENCFF722LJP<br>ENCFF592GWM<br>ENCFF541DRZ<br>ENCFF558UWY |
| ETS1 | ENCAB000AGG | 0.06 | EFO:0007950<br>EFO:0001086<br>EFO:0002067<br>EFO:0002784 | ENCFF896WFR<br>ENCFF461PRP<br>ENCFF511AZU<br>ENCFF980VOD |

| | | | | |
|---|---|---|---|---|
| ETV6 | ENCAB000ABD | -0.0 | EFO:0002067<br>EFO:0002784 | ENCFF426GSY<br>ENCFF745ANU |
| ETV6 | ENCAB997CJG | 0.37 | EFO:0002067<br>EFO:0002784 | ENCFF658SGJ<br>ENCFF116AMK |
| EZH2 | ENCAB000AGH | 0.12 | CL:0002551<br>EFO:0001187<br>CL:0000515<br>EFO:0002791<br>CL:0002327<br>CL:0002553<br>EFO:0002784<br>CL:0000236 | ENCFF340LPI<br>ENCFF260KLJ<br>ENCFF504QZJ<br>ENCFF279MNV<br>ENCFF420KMT<br>ENCFF615NYO<br>ENCFF128OWK<br>ENCFF434OEY |
| EZH2 | ENCAB913HCF | 0.03 | CL:0000182<br>EFO:0005723 | ENCFF976SAN<br>ENCFF324UNA |
| EZH2phosphoT487 | ENCAB000BKV | 0.07 | NTR:0000711<br>EFO:0002074<br>CL:0000103<br>EFO:0005723 | ENCFF314ZKR<br>ENCFF320REA<br>ENCFF689GPW<br>ENCFF687VHB |
| FIP1L1 | ENCAB511DBP | 0.12 | EFO:0002067<br>EFO:0001187 | ENCFF084DTV<br>ENCFF031LBW |
| FOS | ENCAB000AEQ | 0.13 | CL:0002618<br>EFO:0001196<br>EFO:0001203 | ENCFF217ZMF<br>ENCFF327GZX<br>ENCFF170POB |
| FOSL2 | ENCAB000AGL | 0.16 | EFO:0001086<br>EFO:0001187 | ENCFF808RWZ<br>ENCFF054ESU |
| FOXA1 | ENCAB000AGM | 0.16 | EFO:0001086<br>EFO:0001187 | ENCFF152BOT<br>ENCFF297HAX<br>ENCFF167BKY |
| FOXA1 | ENCAB000AGN | 0.2 | EFO:0001187<br>UBERON:0002107 | ENCFF872MGU<br>ENCFF324QGE<br>ENCFF951VPZ |
| FOXA1 | ENCAB301QLE | 0.1 | EFO:0001187<br>EFO:0001203 | ENCFF160RLI<br>ENCFF367TQC |
| FOXA2 | ENCAB000AGO | 0.11 | EFO:0001187<br>UBERON:0002107 | ENCFF184NAC<br>ENCFF168JLI<br>ENCFF293LRQ |

| | | | | |
|---|---|---|---|---|
| FOXK2 | ENCAB625ERS | 0.17 | EFO:0002067<br>EFO:0001187<br>EFO:0002784<br>EFO:0001203 | ENCFF899MQW<br>ENCFF990MTR<br>ENCFF490EQR<br>ENCFF315CHX |
| FUS | ENCAB598VSJ | 0.03 | EFO:0002067<br>EFO:0001187 | ENCFF216YZI<br>ENCFF688ARM |
| GABPA | ENCAB000AGR | 0.06 | EFO:0001086<br>EFO:0002067<br>EFO:0001187<br>UBERON:0002107<br>EFO:0002791<br>EFO:0002784<br>EFO:0003042 | ENCFF520GJC<br>ENCFF124HAC<br>ENCFF225GFQ<br>ENCFF054HJA<br>ENCFF091UDB<br>ENCFF946ACA<br>ENCFF280YAF<br>ENCFF344XWK |
| GABPA | ENCAB728YTO | 0.05 | EFO:0002067<br>EFO:0001203 | ENCFF620TOF<br>ENCFF535BMV |
| GATA2 | ENCAB000AGT | 0.3 | EFO:0002067<br>CL:0002618 | ENCFF173TXA<br>ENCFF987YIJ |
| GATAD2B | ENCAB939ONI | 0.17 | EFO:0002067<br>EFO:0002784<br>EFO:0001203 | ENCFF046BRP<br>ENCFF298AIX<br>ENCFF569CMJ |
| GTF2F1 | ENCAB000AHE | 0.11 | EFO:0002067<br>EFO:0001187<br>EFO:0002791<br>EFO:0003042<br>EFO:0001203 | ENCFF876GXQ<br>ENCFF394JHN<br>ENCFF493PRB<br>ENCFF478HYJ<br>ENCFF343QQE |
| GTF2F1 | ENCAB108WPN | 0.33 | EFO:0002067<br>EFO:0001187 | ENCFF843UHP<br>ENCFF599TWF |
| HCFC1 | ENCAB000AHG | 0.02 | EFO:0002067<br>EFO:0001187<br>EFO:0002784<br>EFO:0001203 | ENCFF722QBB<br>ENCFF167RXK<br>ENCFF401IAI<br>ENCFF485SRU |
| HDAC1 | ENCAB649LYF | 0.16 | EFO:0002067<br>EFO:0001187 | ENCFF069KPS<br>ENCFF557WXK |

| | | | | |
|---|---|---|---|---|
| HDAC2 | ENCAB000AHI | 0.1 | EFO:0001086<br>EFO:0002067<br>EFO:0003042 | ENCFF363GSV<br>ENCFF497YNJ<br>ENCFF814DAF |
| HDAC2 | ENCAB000AHJ | 0.15 | EFO:0002067<br>EFO:0001187<br>EFO:0003042 | ENCFF589GSN<br>ENCFF009IVJ<br>ENCFF741IMY |
| HDGF | ENCAB173GGB | 0.17 | EFO:0002067<br>EFO:0002784<br>EFO:0001203 | ENCFF442WRJ<br>ENCFF161SFU<br>ENCFF575WFB |
| HES1 | ENCAB724OQW | 0.31 | EFO:0002067<br>EFO:0001203 | ENCFF010OOE<br>ENCFF144OPN |
| HNF4A | ENCAB000AHP | 0.15 | EFO:0001187<br>UBERON:0002107 | ENCFF072CXB<br>ENCFF837QHJ<br>ENCFF905JAC |
| HNF4G | ENCAB000AHQ | 0.09 | EFO:0001187<br>UBERON:0002107 | ENCFF497MUF<br>ENCFF086CTA |
| HNRNPK | ENCAB000BFN | 0.09 | EFO:0002067<br>EFO:0001187 | ENCFF828KXG<br>ENCFF984QUV |
| HNRNPL | ENCAB000AVZ | 0.23 | EFO:0002067<br>EFO:0001187 | ENCFF984ESZ<br>ENCFF039CUI |
| HNRNPLL | ENCAB698BTS | 0.12 | EFO:0002067<br>EFO:0001187 | ENCFF662WPN<br>ENCFF890KTX |
| IKZF1 | ENCAB144WPX | 0.32 | EFO:0002067<br>EFO:0001187<br>EFO:0002784 | ENCFF969BZA<br>ENCFF968NOG<br>ENCFF994OQH |
| IKZF1 | ENCAB590IRI | 0.27 | EFO:0002067<br>EFO:0002784 | ENCFF018NNF<br>ENCFF785BTP |
| JUN | ENCAB000AER | 0.12 | EFO:0002067<br>EFO:0003042<br>EFO:0001187<br>EFO:0001203 | ENCFF907UNK<br>ENCFF331IUK<br>ENCFF312GEN<br>ENCFF394CEC<br>ENCFF167WUZ<br>ENCFF881AVX<br>ENCFF672LKE<br>ENCFF032UMW |

| | | | | |
|---|---|---|---|---|
| JUNB | ENCAB000BQG | 0.0 | EFO:0002067<br>EFO:0002784 | ENCFF478XNA<br>ENCFF739XTO |
| JUND | ENCAB000AID | 0.02 | EFO:0001086<br>EFO:0002824<br>EFO:0002067<br>EFO:0001187<br>UBERON:0002107<br>EFO:0003072<br>EFO:0003042<br>EFO:0002784<br>EFO:0001203 | ENCFF587VEY<br>ENCFF569ZCY<br>ENCFF213EYD<br>ENCFF873DJD<br>ENCFF998KDQ<br>ENCFF246HKM<br>ENCFF539GRW<br>ENCFF420PED<br>ENCFF187QQB<br>ENCFF430PEI<br>ENCFF443HNU<br>ENCFF646IUA<br>ENCFF229COM |
| KDM1A | ENCAB000AIH | 0.11 | EFO:0001086<br>EFO:0001187<br>EFO:0003042 | ENCFF768FGG<br>ENCFF316CBQ<br>ENCFF562OAN |
| KDM5A | ENCAB914RSH | 0.06 | EFO:0001086<br>EFO:0001187 | ENCFF149INM<br>ENCFF334HKG |
| LARP7 | ENCAB568DRS | 0.32 | EFO:0002067<br>EFO:0002784 | ENCFF305SLO<br>ENCFF668ZTJ |
| MAFF | ENCAB717EYF | 0.14 | EFO:0002067<br>EFO:0002791<br>EFO:0001187 | ENCFF498MGH<br>ENCFF493TIR<br>ENCFF672LKL |
| MAFK | ENCAB000AIJ | 0.11 | EFO:0001086<br>EFO:0001203<br>EFO:0002067<br>EFO:0001187<br>EFO:0002791<br>EFO:0001196<br>EFO:0002784<br>EFO:0003042 | ENCFF813WJW<br>ENCFF171OJF<br>ENCFF873SVI<br>ENCFF351VGZ<br>ENCFF186AWV<br>ENCFF712RIS<br>ENCFF328IZQ<br>ENCFF893SCL |

| | | | | |
|---|---|---|---|---|
| MAX | ENCAB000AIL | 0.15 | EFO:0002067<br>EFO:0001187<br>UBERON:0002107<br>EFO:0002784<br>EFO:0003042 | ENCFF762LKG<br>ENCFF140PUO<br>ENCFF270NAL<br>ENCFF493ZMX<br>ENCFF618VMC<br>ENCFF669BQN<br>ENCFF900NVQ |
| MAZ | ENCAB000AIM | 0.05 | EFO:0001086<br>EFO:0001187<br>EFO:0001196<br>EFO:0002784<br>EFO:0001203 | ENCFF666YGQ<br>ENCFF144TBQ<br>ENCFF661NNJ<br>ENCFF916AJB<br>ENCFF348STZ |
| MBD2 | ENCAB000BQP | 0.13 | EFO:0002067<br>EFO:0001203 | ENCFF464QAL<br>ENCFF617QSK |
| MLLT1 | ENCAB650PBW | 0.16 | EFO:0002067<br>EFO:0002784<br>EFO:0001203 | ENCFF578NMN<br>ENCFF125MEN<br>ENCFF010AIG |
| MNT | ENCAB000BCL | 0.19 | EFO:0002067<br>EFO:0001187<br>EFO:0001203 | ENCFF562FMQ<br>ENCFF432GSK<br>ENCFF459DYU |
| MNT | ENCAB887GAG | 0.06 | EFO:0002067<br>EFO:0001187 | ENCFF454QQD<br>ENCFF482JSR |
| MTA1 | ENCAB000BCN | 0.24 | EFO:0002067<br>EFO:0001203 | ENCFF801KEW<br>ENCFF225VFR |
| MTA3 | ENCAB000BML | 0.24 | EFO:0002067<br>EFO:0001203 | ENCFF083AZM<br>ENCFF459XLR |
| MXI1 | ENCAB000AIT | 0.08 | EFO:0002067<br>CL:0002319<br>EFO:0002784<br>EFO:0003072 | ENCFF255WJM<br>ENCFF199HGX<br>ENCFF243QTL<br>ENCFF116RCK |

| | | | | |
|---|---|---|---|---|
| MYC | ENCAB000AET | 0.15 | EFO:0001086<br>EFO:0002067<br>EFO:0003042<br>EFO:0001203 | ENCFF392JJN<br>ENCFF542GMN<br>ENCFF339AQP<br>ENCFF605WXD<br>ENCFF370EQJ<br>ENCFF492XUU<br>ENCFF300OKR<br>ENCFF527EGF<br>ENCFF658XME<br>ENCFF700TLG |
| NANOG | ENCAB000AIX | 0.28 | EFO:0007950<br>EFO:0003042 | ENCFF621PFM<br>ENCFF794GVQ |
| NCOR1 | ENCAB805KAO | 0.26 | EFO:0002067<br>EFO:0001187 | ENCFF638IIC<br>ENCFF616RSZ |
| NEUROD1 | ENCAB000BDX | 0.28 | EFO:0002067<br>EFO:0001203 | ENCFF059LJD<br>ENCFF755APC |
| NFATC3 | ENCAB375FHW | 0.15 | EFO:0002067<br>EFO:0002784 | ENCFF430JFH<br>ENCFF704PDA |
| NFE2L2 | ENCAB800OND | 0.1 | EFO:0001086<br>EFO:0002791<br>EFO:0001187<br>EFO:0001196 | ENCFF882YLO<br>ENCFF305KIK<br>ENCFF474PPT<br>ENCFF418TUX |
| NFXL1 | ENCAB208FGX | 0.26 | EFO:0002067<br>EFO:0002784<br>EFO:0001203 | ENCFF329STX<br>ENCFF860IXB<br>ENCFF927DIO |
| NFYB | ENCAB000AJD | 0.11 | EFO:0002067<br>EFO:0002784 | ENCFF510NDO<br>ENCFF009NXK |
| NKRF | ENCAB893CIV | 0.14 | EFO:0002067<br>EFO:0002784 | ENCFF084NXU<br>ENCFF520QSR |
| NONO | ENCAB097MPS | 0.11 | EFO:0002067<br>EFO:0001187 | ENCFF823CQK<br>ENCFF420QKI |
| NONO | ENCAB349QGP | 0.34 | EFO:0002067<br>EFO:0001203 | ENCFF515YFU<br>ENCFF800CDQ |
| NR2C1 | ENCAB324ARN | 0.18 | EFO:0002067<br>EFO:0002784 | ENCFF462AKP<br>ENCFF023XHV |
| NR2F1 | ENCAB000ATZ | 0.2 | EFO:0002067<br>EFO:0002784 | ENCFF363IQN<br>ENCFF531KOV |

| | | | | |
|---|---|---|---|---|
| NR2F2 | ENCAB000AJH | 0.24 | EFO:0002067 UBERON:0002107 | ENCFF118HUH ENCFF819WNB ENCFF379TVQ |
| NR2F6 | ENCAB854ATP | 0.19 | EFO:0002067 EFO:0001187 | ENCFF350CKI ENCFF194VBK |
| NRF1 | ENCAB000AJI | 0.05 | EFO:0001203 EFO:0001187 EFO:0002784 EFO:0003042 | ENCFF652BRY ENCFF407IVS ENCFF418DKQ ENCFF269RME |
| NRF1 | ENCAB000BLM | 0.01 | EFO:0002067 EFO:0001187 | ENCFF313RFR ENCFF626VDA ENCFF543STN |
| PAX5 | ENCAB000AJS | 0.18 | EFO:0002785 EFO:0002786 EFO:0002784 | ENCFF997VAB ENCFF196JGP ENCFF987CQF |
| PAX8 | ENCAB000BOS | 0.34 | EFO:0002784 EFO:0001203 | ENCFF473UHQ ENCFF992JWY |
| PBX2 | ENCAB697XQW | 0.11 | EFO:0002067 EFO:0001187 | ENCFF925OBR ENCFF709YWO |
| PCBP1 | ENCAB000BFX | 0.0 | EFO:0002067 EFO:0001187 | ENCFF467RYH ENCFF487WAN |
| PCBP2 | ENCAB000BFY | 0.33 | EFO:0002067 EFO:0001187 | ENCFF642XRH ENCFF941XZW |
| PHF8 | ENCAB757LUG | 0.06 | EFO:0001086 EFO:0001187 | ENCFF907WHF ENCFF202WIO |

| | | | | |
|---|---|---|---|---|
| POLR2A | ENCAB000AOC | 0.02 | EFO:0001086<br>EFO:0002824<br>EFO:0001203<br>EFO:0002067<br>EFO:0001187<br>EFO:0002786<br>EFO:0002791<br>CL:0002618<br>EFO:0002785<br>EFO:0002784<br>EFO:0003042 | ENCFF246QVY<br>ENCFF271RGE<br>ENCFF403ZEO<br>ENCFF422HDN<br>ENCFF964EVA<br>ENCFF565SUC<br>ENCFF387VGY<br>ENCFF455ZLJ<br>ENCFF021HUZ<br>ENCFF099NYA<br>ENCFF741JES<br>ENCFF798PUX<br>ENCFF730DLS<br>ENCFF664KTN<br>ENCFF668VIK<br>ENCFF915LKZ<br>ENCFF182YZG |
| POLR2AphosphoS2 | ENCAB000AOB | 0.02 | EFO:0001086<br>EFO:0002067<br>EFO:0001187<br>EFO:0002784 | ENCFF652YVO<br>ENCFF156MIR<br>ENCFF266OPF<br>ENCFF847DXY |
| POLR2G | ENCAB392AZR | 0.19 | EFO:0002067<br>EFO:0001187 | ENCFF283CUY<br>ENCFF551IJP |
| PRPF4 | ENCAB624FBH | 0.4 | EFO:0002067<br>EFO:0001187 | ENCFF417RQZ<br>ENCFF908QCS |
| PTBP1 | ENCAB553IRN | 0.25 | EFO:0002067<br>EFO:0001187 | ENCFF875ZPV<br>ENCFF917HXV |

| | | | | |
|---|---|---|---|---|
| RAD21 | ENCAB000AKG | 0.04 | EFO:0001086<br>CL:0002319<br>EFO:0001187<br>UBERON:0002107<br>EFO:0003072<br>EFO:0001196<br>EFO:0002784<br>EFO:0003042 | ENCFF557OCR<br>ENCFF897QCA<br>ENCFF654EGO<br>ENCFF454TRL<br>ENCFF895JAW<br>ENCFF295GOD<br>ENCFF255FRL<br>ENCFF874VFZ<br>ENCFF060IVS<br>ENCFF229WFR<br>ENCFF093XOJ<br>ENCFF315BSV |
| RAD21 | ENCAB697XQW | 0.09 | EFO:0001187<br>EFO:0001203 | ENCFF330VPL<br>ENCFF081TVG |
| RAD51 | ENCAB406ZKT | 0.13 | EFO:0002067<br>EFO:0001187<br>EFO:0002784<br>EFO:0001203 | ENCFF859MBC<br>ENCFF996NBR<br>ENCFF091AYX<br>ENCFF740OPF |
| RB1 | ENCAB000BFB | 0.02 | EFO:0002067<br>EFO:0002784 | ENCFF034OSV<br>ENCFF328QZM |
| RBBP5 | ENCAB000AKH | 0.03 | EFO:0002067<br>EFO:0003042 | ENCFF607WCG<br>ENCFF666PCE |
| RBFOX2 | ENCAB592TEY | 0.05 | EFO:0002067<br>EFO:0001187 | ENCFF871YRG<br>ENCFF232ASB |
| RBM22 | ENCAB476CFS | 0.32 | EFO:0002067<br>EFO:0001187 | ENCFF420IBN<br>ENCFF305WYD |
| RBM39 | ENCAB975CWB | 0.25 | EFO:0002067<br>EFO:0001187 | ENCFF420ALF<br>ENCFF503DIK |
| RCOR1 | ENCAB000AFK | 0.15 | EFO:0003072<br>EFO:0001187<br>EFO:0002784<br>EFO:0001196 | ENCFF073ADA<br>ENCFF470ZMK<br>ENCFF987VKU<br>ENCFF139EBY |

| | | | | |
|---|---|---|---|---|
| REST | ENCAB000AJK | 0.05 | EFO:0001086<br>EFO:0002067<br>EFO:0001187<br>EFO:0003072<br>UBERON:0002107<br>EFO:0002791<br>EFO:0002784<br>EFO:0003042 | ENCFF023ZUW<br>ENCFF208NUB<br>ENCFF107EWI<br>ENCFF403CAJ<br>ENCFF540FXB<br>ENCFF313CII<br>ENCFF669XCW<br>ENCFF986RRJ<br>ENCFF796YFZ<br>ENCFF274BBE<br>ENCFF288XHG<br>ENCFF178WRO |
| RFX5 | ENCAB000AKJ | 0.04 | EFO:0001086<br>EFO:0001203<br>EFO:0002067<br>EFO:0001187<br>EFO:0003072<br>EFO:0002784<br>EFO:0003042 | ENCFF502JJJ<br>ENCFF062WBN<br>ENCFF259LNG<br>ENCFF201YKU<br>ENCFF179WDI<br>ENCFF059GWW<br>ENCFF103MPW |
| RNF2 | ENCAB000BEA | 0.06 | EFO:0002067<br>EFO:0001187 | ENCFF380SYL<br>ENCFF820LKT |
| RNF2 | ENCAB790JUW | 0.26 | EFO:0002067<br>EFO:0003042 | ENCFF462AZY<br>ENCFF283MNG |
| RXRA | ENCAB000AKN | 0.2 | EFO:0003042<br>EFO:0001187<br>EFO:0002784<br>UBERON:0002107 | ENCFF313BDA<br>ENCFF105TFM<br>ENCFF430SIE<br>ENCFF572MCI<br>ENCFF201KGJ |
| SAP30 | ENCAB000AKO | 0.15 | EFO:0002067<br>EFO:0003042 | ENCFF103RHL<br>ENCFF193TFR |

| | | | | |
|---|---|---|---|---|
| SIN3A | ENCAB000AKR | 0.05 | EFO:0001086<br>EFO:0001203<br>EFO:0002067<br>EFO:0002784<br>EFO:0003042 | ENCFF514BGQ<br>ENCFF802JAN<br>ENCFF050CYK<br>ENCFF567BJI<br>ENCFF220RUS |
| SIN3A | ENCAB000AKS | 0.08 | EFO:0001086<br>EFO:0002067<br>EFO:0001187<br>EFO:0003072<br>EFO:0003042 | ENCFF663RUS<br>ENCFF407VGB<br>ENCFF635YMI<br>ENCFF708HTR<br>ENCFF905VZD |
| SIN3B | ENCAB194LUB | 0.19 | EFO:0002067<br>EFO:0001187 | ENCFF193DQZ<br>ENCFF543INR |
| SIX5 | ENCAB000AKV | 0.07 | EFO:0001086<br>EFO:0002067<br>EFO:0002784<br>EFO:0003042 | ENCFF864TFH<br>ENCFF189NMX<br>ENCFF644BNN<br>ENCFF247LOF |
| SKIL | ENCAB000BNW | 0.3 | EFO:0002067<br>EFO:0002784 | ENCFF254QDM<br>ENCFF903KEI |
| SMAD1 | ENCAB000AYH | 0.16 | EFO:0002067<br>EFO:0002784 | ENCFF987PGY<br>ENCFF084BUP |
| SMAD5 | ENCAB000AAG | 0.02 | EFO:0002067<br>EFO:0002784 | ENCFF855SJG<br>ENCFF069AAY |
| SMARCA4 | ENCAB000AEO | 0.26 | CL:0000103<br>EFO:0002067 | ENCFF703NAE<br>ENCFF482JUI |
| SMARCA5 | ENCAB528BVW | 0.21 | EFO:0002067<br>EFO:0002784<br>EFO:0001203 | ENCFF052STI<br>ENCFF481TNF<br>ENCFF618JNX |
| SMARCC2 | ENCAB313DWJ | 0.18 | EFO:0002067<br>EFO:0001187 | ENCFF150NHK<br>ENCFF751ZVX |
| SMARCE1 | ENCAB550CKA | 0.27 | EFO:0002067<br>EFO:0001203 | ENCFF435SZS<br>ENCFF761NKP |
| SMC3 | ENCAB000AKX | 0.02 | EFO:0001086<br>EFO:0002067<br>CL:0002319<br>EFO:0001187<br>EFO:0001196<br>EFO:0002784 | ENCFF035YWE<br>ENCFF572RPI<br>ENCFF944KJO<br>ENCFF380ZXB<br>ENCFF256LDD<br>ENCFF175UEE |

| | | | | |
|---|---|---|---|---|
| SNIP1 | ENCAB027LVC | 0.14 | EFO:0002067<br>EFO:0001203 | ENCFF529BDW<br>ENCFF455HWV |
| SOX6 | ENCAB000BOG | 0.43 | EFO:0002067<br>EFO:0001187 | ENCFF944LNI<br>ENCFF431STY |
| SP1 | ENCAB000AKY | 0.16 | UBERON:0002107<br>EFO:0001086<br>EFO:0001187<br>EFO:0003042 | ENCFF404OSB<br>ENCFF500JFI<br>ENCFF175VXL<br>ENCFF433EFF<br>ENCFF978TMH |
| SP1 | ENCAB000BAV | 0.25 | EFO:0002067<br>EFO:0001187<br>EFO:0001203 | ENCFF577EMC<br>ENCFF452LDK<br>ENCFF735WMX |
| SPI1 | ENCAB000AKF | 0.15 | EFO:0002067<br>EFO:0002785<br>EFO:0002784 | ENCFF414ECK<br>ENCFF744AGB<br>ENCFF071ZMW |
| SREBF1 | ENCAB000ALC | 0.14 | EFO:0001086<br>EFO:0002067<br>EFO:0001203 | ENCFF275WAD<br>ENCFF777MYW<br>ENCFF624DDK |
| SRF | ENCAB000ALE | 0.14 | EFO:0002784<br>EFO:0003042 | ENCFF345IDL<br>ENCFF182IFE<br>ENCFF829SEJ |
| STAT1 | ENCAB000ALF | 0.16 | EFO:0002067<br>EFO:0002784 | ENCFF323QQU<br>ENCFF747ICD<br>ENCFF431NLF<br>ENCFF646MXG |
| STAT5A | ENCAB000ALI | 0.29 | EFO:0002067<br>EFO:0002784 | ENCFF383YEA<br>ENCFF517IXK |
| SUZ12 | ENCAB000BEB | -0.02 | EFO:0002067<br>EFO:0001187 | ENCFF856HYC<br>ENCFF239LRW |

| | | | | |
|---|---|---|---|---|
| TAF1 | ENCAB000ALM | 0.02 | EFO:0001086<br>EFO:0001203<br>EFO:0002067<br>EFO:0001187<br>EFO:0002786<br>EFO:0003072<br>UBERON:0002107<br>EFO:0002791<br>EFO:0002785<br>EFO:0002784<br>EFO:0003042 | ENCFF033PLJ<br>ENCFF762MGC<br>ENCFF234TBW<br>ENCFF471NIK<br>ENCFF453TIB<br>ENCFF278XOE<br>ENCFF540AAP<br>ENCFF870SFJ<br>ENCFF886KDK<br>ENCFF214OJW<br>ENCFF423CTO |
| TARDBP | ENCAB000AUF | 0.02 | EFO:0002067<br>EFO:0002784 | ENCFF641AXD<br>ENCFF871LZM |
| TARDBP | ENCAB000BAX | 0.16 | EFO:0002067<br>EFO:0002784<br>EFO:0001203 | ENCFF909RMQ<br>ENCFF668JHK<br>ENCFF233RBO |
| TARDBP | ENCAB057RGG | 0.18 | EFO:0002067<br>EFO:0001187 | ENCFF448YOS<br>ENCFF696QPP |
| TBL1XR1 | ENCAB000ALP | 0.1 | EFO:0002067<br>EFO:0001187<br>EFO:0002784 | ENCFF126KGW<br>ENCFF392JWA<br>ENCFF239WFN |
| TBP | ENCAB000ALR | 0.08 | EFO:0002067<br>EFO:0001187<br>EFO:0002791<br>EFO:0002784<br>EFO:0003042 | ENCFF370YGS<br>ENCFF748YXF<br>ENCFF302RQH<br>ENCFF534GKQ<br>ENCFF896UZB |
| TCF12 | ENCAB000ALT | 0.22 | EFO:0001086<br>EFO:0001187<br>EFO:0002784<br>EFO:0003042 | ENCFF740HPV<br>ENCFF768VSH<br>ENCFF299JYV<br>ENCFF228CDD |
| TCF12 | ENCAB506UYG | 0.38 | EFO:0002067<br>EFO:0002784 | ENCFF897RYA<br>ENCFF912LXU |

| | | | | |
|---|---|---|---|---|
| TCF7 | ENCAB000ACR | 0.27 | EFO:0002067<br>EFO:0001187<br>EFO:0002784 | ENCFF152RNE<br>ENCFF512IAI<br>ENCFF928MIN |
| TOE1 | ENCAB755SML | 0.19 | EFO:0002067<br>EFO:0001187<br>EFO:0001203 | ENCFF539FVQ<br>ENCFF014WCO<br>ENCFF144VMM |
| TRIM22 | ENCAB000BNM | 0.33 | EFO:0001187<br>EFO:0002784<br>EFO:0001203 | ENCFF063GDN<br>ENCFF452VLA<br>ENCFF830TFU<br>ENCFF552WAH |
| U2AF1 | ENCAB298WVV | 0.33 | EFO:0002067<br>EFO:0001187 | ENCFF034KUO<br>ENCFF482DRO |
| USF1 | ENCAB000AMF | 0.08 | EFO:0002067<br>EFO:0001187<br>EFO:0002784<br>EFO:0003042 | ENCFF701QXK<br>ENCFF717KGR<br>ENCFF914IFQ<br>ENCFF699HXL |
| USF2 | ENCAB000AMH | 0.07 | EFO:0001086<br>EFO:0002067<br>EFO:0001196<br>EFO:0002784<br>EFO:0003042 | ENCFF514SWA<br>ENCFF425FVY<br>ENCFF593EOW<br>ENCFF938BOJ<br>ENCFF710JBU |
| XRCC5 | ENCAB308AOH | 0.04 | EFO:0002067<br>EFO:0001187 | ENCFF929TWP<br>ENCFF790ZAQ |
| YBX1 | ENCAB493UWX | 0.17 | EFO:0001187<br>EFO:0002784<br>EFO:0001203 | ENCFF332FUE<br>ENCFF247VVK<br>ENCFF500RBO |
| YY1 | ENCAB000ANS | 0.03 | EFO:0002067<br>EFO:0002786 | ENCFF072IHJ<br>ENCFF635XCI<br>ENCFF024TJO |

| | | | | |
|---|---|---|---|---|
| YY1 | ENCAB000ANT | 0.04 | EFO:0001086<br>EFO:0002824<br>EFO:0002067<br>EFO:0001187<br>EFO:0003072<br>UBERON:0002107<br>EFO:0002785<br>EFO:0002784<br>EFO:0003042 | ENCFF953BTB<br>ENCFF613DTQ<br>ENCFF363UWP<br>ENCFF094BQZ<br>ENCFF177YDT<br>ENCFF509GYP<br>ENCFF538VYU<br>ENCFF223MUF<br>ENCFF838VFX<br>ENCFF459TWF |
| ZBED1 | ENCAB000AAK | 0.21 | EFO:0002067<br>EFO:0002784 | ENCFF388TYU<br>ENCFF630FLK |
| ZBTB33 | ENCAB000AML | 0.15 | EFO:0001086<br>EFO:0002824<br>EFO:0001187<br>UBERON:0002107<br>EFO:0002784 | ENCFF422MCZ<br>ENCFF593ZJA<br>ENCFF943WRA<br>ENCFF773OQL<br>ENCFF727ZIT<br>ENCFF882UHR |
| ZBTB33 | ENCAB292USO | 0.21 | EFO:0002067<br>EFO:0002784<br>EFO:0001203 | ENCFF556STK<br>ENCFF780WLS<br>ENCFF475DID |
| ZBTB40 | ENCAB373DVF | 0.08 | EFO:0002067<br>EFO:0001187<br>EFO:0002784<br>EFO:0001203 | ENCFF932XEU<br>ENCFF624WDI<br>ENCFF088LZZ<br>ENCFF084IUW |
| ZBTB7A | ENCAB000AMM | 0.07 | EFO:0002067<br>EFO:0001187 | ENCFF953JQD<br>ENCFF245LRG |
| ZC3H11A | ENCAB000AMN | 0.44 | EFO:0001086<br>EFO:0002067 | ENCFF478PGJ<br>ENCFF415SIS |

| | | | | |
|---|---|---|---|---|
| ZFP36 | ENCAB118PND | 0.09 | EFO:0001086<br>EFO:0002067<br>EFO:0001187<br>EFO:0002791<br>EFO:0002784 | ENCFF166GKK<br>ENCFF429XQI<br>ENCFF137JHO<br>ENCFF224WII<br>ENCFF763HPQ |
| ZFX | ENCAB657HDP | 0.05 | EFO:0001203<br>EFO:0002824 | ENCFF215SIC<br>ENCFF775BWJ |
| ZHX1 | ENCAB361RPF | 0.04 | EFO:0002067<br>EFO:0002791 | ENCFF267DZF<br>ENCFF495BPY |
| ZHX2 | ENCAB000ATW | 0.17 | EFO:0001187<br>EFO:0001203 | ENCFF694ZRC<br>ENCFF964KDQ |
| ZKSCAN1 | ENCAB000AMP | 0.26 | EFO:0002067<br>EFO:0001187<br>EFO:0001203 | ENCFF687REM<br>ENCFF704VDI<br>ENCFF721NEC |
| ZMYM3 | ENCAB426WVA | 0.19 | EFO:0002067<br>EFO:0001187 | ENCFF195IFB<br>ENCFF769SEZ |
| ZNF143 | ENCAB000AMR | 0.06 | EFO:0002067<br>EFO:0002784<br>EFO:0003042 | ENCFF933WSP<br>ENCFF700GZI<br>ENCFF193POQ<br>ENCFF153TQR |
| ZNF207 | ENCAB000BNU | 0.24 | EFO:0002784<br>EFO:0001203 | ENCFF676BIG<br>ENCFF621ZSK |
| ZNF217 | ENCAB182PZR | 0.15 | EFO:0002784<br>EFO:0001203 | ENCFF200SLC<br>ENCFF620RPM |
| ZNF24 | ENCAB060JJI | 0.1 | EFO:0002067<br>EFO:0001187 | ENCFF858WPR<br>ENCFF723JDW |
| ZNF24 | ENCAB198YAJ | 0.08 | EFO:0002067<br>EFO:0001187<br>EFO:0002784<br>EFO:0001203 | ENCFF619BFO<br>ENCFF313HBL<br>ENCFF260CBQ<br>ENCFF904QAD |
| ZNF282 | ENCAB503NQV | 0.0 | EFO:0002067<br>EFO:0001187 | ENCFF596JDS<br>ENCFF482XNG |
| ZNF592 | ENCAB438BKV | 0.1 | EFO:0002067<br>EFO:0001203 | ENCFF972UGK<br>ENCFF541HRT |
| ZNF687 | ENCAB146FZU | 0.14 | EFO:0002784<br>EFO:0001203 | ENCFF137BRA<br>ENCFF329QYZ |
| ZSCAN29 | ENCAB211EDR | 0.1 | EFO:0002067<br>EFO:0002784 | ENCFF214NJL<br>ENCFF979GFF |

| ENCODE Name | Cell Type |
|---|---|
| CL:0000062 | osteoblast |
| CL:0000103 | bipolar neuron |
| CL:0000127 | astrocyte |
| CL:0000182 | hepatocyte |
| CL:0000192 | smooth muscle cell |
| CL:0002319 | neural cell |
| CL:0002372 | myotube |
| CL:0002551 | fibroblast of dermis |
| CL:0002618 | endothelial cell of umbilical vein |
| EFO:0001086 | A549 |
| EFO:0001187 | HepG2 |
| EFO:0001196 | IMR-90 |
| EFO:0001203 | MCF-7 |
| EFO:0002067 | K562 |
| EFO:0002074 | PC-3 |
| EFO:0002784 | GM12878 |
| EFO:0002785 | GM12891 |
| EFO:0002791 | HeLa-S3 |
| EFO:0002824 | HCT116 |
| EFO:0003042 | H1 |
| EFO:0003072 | SK-N-SH |
| EFO:0006711 | OCI-LY7 |
| EFO:0007950 | GM23338 |
| UBERON:0002107 | liver |
| NTR:0000711 | neural progenitor cell |

Table 2: Cell type names and their corresponding ENCODE representations.
